# Human genetic adaptation related to cellular zinc homeostasis

**DOI:** 10.1101/2022.06.29.498106

**Authors:** Ana Roca-Umbert, Jorge Garcia-Calleja, Marina Vogel-González, Alejandro Fierro-Villegas, Gerard Ill-Raga, Víctor Herrera-Fernández, Anja Bosnjak, Gerard Muntané, Esteban Gutiérrez, Felix Campelo, Rubén Vicente, Elena Bosch

## Abstract

*SLC30A9* encodes a ubiquitously zinc transporter (ZnT9) and has been consistently suggested as a candidate for positive selection in humans. However, no direct adaptive molecular phenotype has been demonstrated. Our results provide evidence for directional selection operating in two major complementary haplotypes in Africa and East Asia. These haplotypes are associated with differential gene expression but also differ in the Met50Val substitution (rs1047626) in ZnT9, which we show is found in homozygosis in the Denisovan genome and displays accompanying signatures suggestive of archaic introgression. Although we found no significant differences in systemic zinc content between individuals with different rs1047626 genotypes, we demonstrate that the expression of the derived isoform (ZnT9 50Val) in HEK293 cells shows a gain of function when compared with the ancestral (ZnT9 50Met) variant. Notably, the ZnT9 50Val variant was found associated with differences in zinc handling by the mitochondria and endoplasmic reticulum, with an impact on mitochondrial metabolism. Given the essential role of the mitochondria in skeletal muscle and since the derived allele at rs1047626 is known to be associated with greater susceptibility to several neuropsychiatric traits, we propose that adaptation to cold may have driven this selection event, while also impacting predisposition to neuropsychiatric disorders in modern humans.

**Author Summary:** Contrasting continental signatures of positive natural selection have been previously found in the human *SLC30A9* gene encoding the protein ZnT9, which transports zinc across cell membranes. Here we investigate the genetic variants that have been targeted by natural selection in the surrounding region of this gene and which molecular and whole-body changes may have brought about. We found that two major *SLC30A9* variant combinations (haplotypes) that are extremely frequent in Africa and East Asia, respectively, are expressed differentially. These two haplotypes also differ at one site that creates an amino acid difference at ZnT9; the version most often found outside Africa avoiding zinc overload in the endoplasmic reticulum and mitochondria and directly influencing mitochondrial activity. Moreover, we found that this substitution, which is known to be associated with greater susceptibility to several neuropsychiatric disorders, is present in the Denisova and displays accompanying patterns of variation that could be suggestive of adaptive introgression. Since mitochondria play an important role in skeletal muscle energy metabolism, we speculate that adaptation to cold may have driven this selection event outside Africa, while also impacting predisposition to neuropsychiatric disorders in modern humans.

## Introduction

How adaptation has shaped current genetic diversity in human populations is a long-standing question in evolutionary genetics. In turn, it has been shown that the identification and functional deciphering of adaptive variants in our genome can reveal biologically relevant variation with important phenotypic consequences in current populations [1,2]. Recent access to whole-genome sequencing data from diverse human populations coupled with improvements in methods that detect the molecular signatures of natural selection in the genome have facilitated the identification of hundreds of candidate genes for positive selection in humans [3]. As a result, several candidate regions for selection have been successfully associated with diverse adaptive phenotypes and their corresponding selective pressures, which were mostly related to our diet, immune response, high altitude environment, and UV radiation [2,4,5]. However, our understanding of the adaptive phenotypes and functional variants underlying most of the signatures of positive selection in the human genome is still limited.

Zinc is an essential micronutrient with different structural, catalytic, and regulatory roles in the human body [6–8]. As adequate zinc levels are fundamental for maintaining good health, the systemic and cellular homeostasis of zinc must be tightly regulated. The proteins controlling the import and export of zinc across cell membranes are known as Zinc Transporters (ZT) [7,9]. The 24 ZTs identified in humans can be classified in two families: the ZIP family, whose members import zinc into the cytosol, and the ZnT family, whose function is to export zinc outside the cell or into the cell organelles. The 14 ZIP transporters are encoded by the *SLC39A1-14* genes, whereas the 10 ZnT transporters are encoded by the *SLC30A1-10* genes [7,10–12]. Variation in this set of 24 ZT encoding genes can lead to zinc dysregulation at a cellular or systemic level and, given the fundamental role of zinc in the human organism, such zinc imbalance might cause alterations in distinct phenotypes upon which adaptive selection could act [13]. In this context, a non-synonymous substitution at the *SLC39A4* gene, which encodes the most important intestinal zinc uptake transporter (ZIP4), has been shown to produce differential cellular zinc uptake and presents extreme population differentiation because of a local positive selection event in Sub-Saharan Africa [14,15].

The *SLC30A9* gene (chr4: 41,990,502-42,090,461; GRCh38) has been repeatedly reported as a top candidate region for positive selection in several genome-wide scans of selection, mostly in East Asian populations [14,16–20]. This gene encodes the ZnT9 protein, a recently characterized ZT in the mitochondria [21,22]. Moreover, the patterns of genetic variation and population differentiation at *SLC30A9* have also been investigated in candidate-based selection studies analyzing human adaptation in relation to zinc homeostasis [13,23,24]. In most of these previous studies, the adaptive signals in *SLC30A9* have been attributed to a non-synonymous single nucleotide polymorphism (SNP) (rs1047626, A>G at chr4:42001654; GRCh38) resulting in the substitution of a methionine by a valine in the N-terminus of the ZnT9 transporter (Met50Val) [13,17,23,24]. In agreement with the strong signatures of recent positive selection detected in East Asia, this non-synonymous SNP presents extremely high levels of population differentiation, where the ancestral A-allele (encoding Met) is at high frequency in African populations, whereas the derived G-allele (encoding Val) is nearly fixed in East Asian populations and at intermediate frequencies in European, South Asian, and American populations. Moreover, contrasting extended haplotypes and selection signals along the *SLC30A9* region have been reported between Africans and East Asians [13]. Furthermore, the derived G-allele has been shown to correlate with the distribution of human zinc deficiency worldwide [13] and to be in moderate linkage disequilibrium with three variants in the 3’ flanking region of the *SLC30A9* gene, which are QTLs influencing zinc content in the liver [23]. However, to date, no experimental validation has been performed to demonstrate the functional relevance of this putative adaptive variant regarding zinc transport.

Here, we first compiled evidence for a selective sweep in the *SLC30A9* gene region, inferred the past allele trajectories and selection coefficients for two putative adaptive SNP variants characterizing the two major *SLC30A9* haplotypes in humans, traced back their genetic origins and then investigated the phenotypical relevance of the Met50Val substitution (rs1047626) in ZnT9. Accordingly, we overexpressed ZnT9 in HEK293 cells and explored whether the two variants of the Met50Val substitution induce differences in the expression levels and localization of ZnT9 and other ZTs, and whether they differentially affect zinc transport in the cytosol, mitochondria, and endoplasmic reticulum (ER). Our results show that the ZnT9 Met50Val substitution sits in an unusual haplotype, which shows suggestive signatures of archaic introgression, and demonstrate relevant differences in mitochondrial and ER zinc homeostasis directly influencing mitochondrial activity and cell viability.

## Results

### Signals of positive selection outside Africa

We first explored the genomic signatures of the selective sweep previously reported in the *SLC30A9* gene region [14,16–18,20,25]. Hence, we compiled evidence from different summary statistics and selection tests available at the PopHuman browser for the 26 human populations from the 1000 Genomes Project (1000 GP; http://pophuman.uab.cat) [26]. To summarize and visualize the continental patterns of positive selection along the *SLC30A9* region, we extracted the F_ST_, XP-EHH, and Fay and Wu’s H values in three populations used as a reference for Africa (Yoruba: YRI), Europe (Residents of Utah with North and Western European ancestry: CEU), and Asia (Han Chinese: CHB) (Fig. 1A). Unusual high levels of population differentiation were confirmed along the *SLC30A9* region when comparing CEU and CHB with YRI. Similarly, when applying the XP-EHH statistic, we detected genome-wide departures for long haplotypes in CEU and above all in CHB (using YRI as the reference). Furthermore, a significant excess of derived alleles was detected around the *SLC30A9* region in both CHB and CEU populations when analyzing the site frequency spectrum with the Fay and Wu’s H statistic. Notably, most of these signatures of selection replicate for the remaining 1000 GP populations available at the PopHuman browser in all continents except Africa.

**Figure 1.**
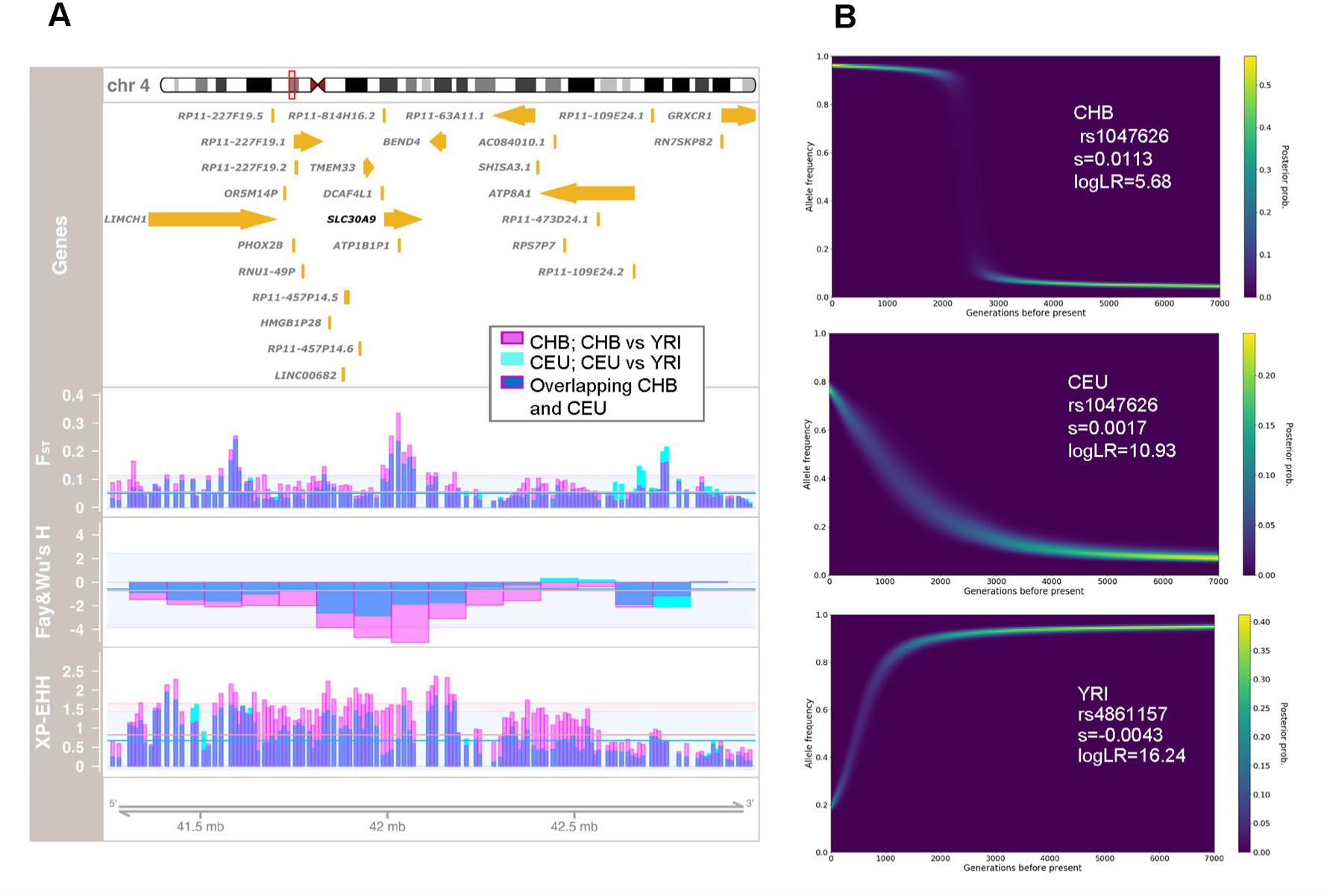
Signals of positive selection at *SLC30A9*. (A) Signatures of adaptation in the *SLC30A9* region. Chromosomal location and genes are displayed at the top. In the first selection panel, window-based F_ST_ values when comparing CEU versus YRI and CHB versus YRI are represented by cyan and pink bars, respectively. Similarly, window-based Fay and Wu’s H values and XP-EHH scores (using YRI as the reference population) in CEU and CHB are shown with the same color scheme in the middle and bottom selection panels, respectively. Overlapping population scores can result in blue shades. Horizontal cyan and pink lines represent the genomic average of each test for each population, whereas 2 standard deviations are represented by the background shade in each case as a mean to interpret the genome-wide significance of each selection signature. All selection test values were extracted from the corresponding 1000 GP populations as available at the PopHuman browser (http://pophuman.uab.cat) [26] and represented using the Gviz package [61]. (B) Allele frequency trajectories and selection coefficients inferred with CLUES for rs1047626 in CHB and CEU and for rs4861157 in YRI.

We next investigated which variants along the *SLC30A9* gene could underlie the detected signals of selection (see complete list in Table S1). From a total of 354 biallelic SNPs with MAF ≥ 0.02 in *SLC30A9* gene, we initially selected as candidates those variants with high population differentiation when comparing YRI with CEU and CHB. From the resulting 235 variants, we further considered as putatively adaptive those that were either non-synonymous, present at the UTR regions of the *SLC30A9* gene, or had a CADD Phred score ≥ 10, as such a value predicts that the variant falls within the top 10% of the most deleterious (i.e., functional) variants of the genome [27]. A total of 18 SNPs matched our filtering criteria (Table 1). Among these, we identified a non-synonymous SNP (rs1047626, resulting in the Met50Val substitution), two synonymous variants, five variants at the 3’ UTR, and ten intronic variants. According to the GTEX portal (https://www.gtexportal.org/home/; last accessed 13/04/2022), all 18 putative adaptive variants are eQTLs, with the most common allele in CHB and CEU associated with reduced *SLC30A9* expression mostly in brain (Figs. S1-S19). Moreover, iSAFE [28] similarly pinpointed several eQTLs (including rs1047626) among the most likely candidate mutations favored by selection along the region (chr4: 41,853,671-42,153,671; GRCh37) in both CEU and CHB (Table 1, Tables S2-3, Fig. S20).

**Table 1.** Putative adaptive variants in the *SLC30A9* region.

| SNP | Ref | Alt | SNP type | AFR | EUR | EAS | SAS | AMR | CADD<br>Phred score | iSAFE<br>CHB | iSAFE<br>CEU | eQTLs <sup>a</sup><br>(Tissue; NES; p-value) |
| --- | --- | --- | --- | --- | --- | --- | --- | --- | --- | --- | --- | --- |
| rs2581434 | C | G | synonymous | 0.246 | 0.815 | 0.959 | 0.860 | 0.840 | 12.00 |  |  | Brain-NA (basal ganglia); -0.23; 7.9 x 10<sup>-10</sup> |
| rs1047626 | A | G | non-synonymous | 0.105 | 0.757 | 0.948 | 0.780 | 0.768 | 0.001 | 1.302 | 0.588 | Brain-Cortex; -0.24; 4.4 x 10<sup>-10</sup> |
| rs2581452 | C | T | intronic | 0.193 | 0.776 | 0.958 | 0.800 | 0.803 | 11.38 |  |  | Brain-NA (basal ganglia); -0.24; 2.5 x 10<sup>-11</sup> |
| rs2660319 | G | A | intronic | 0.146 | 0.756 | 0.949 | 0.780 | 0.770 | 12.93 | 1.209 | 0.588 | Brain-NA (basal ganglia); -0.22; 5.0 x 10<sup>-10</sup> |
| rs1848182 | A | G | intronic | 0.193 | 0.776 | 0.958 | 0.800 | 0.803 | 10.10 |  |  | Brain-NA (basal ganglia); -0.24; 2.5 x 10<sup>-11</sup> |
| rs2581424 | G | A | intronic | 0.188 | 0.776 | 0.958 | 0.800 | 0.804 | 10.78 |  |  | Brain-NA (basal ganglia); -0.23; 3.1 x 10<sup>-10</sup> |
| rs15857 | C | A | synonymous | 0.188 | 0.776 | 0.958 | 0.800 | 0.804 | 14.93 |  |  | Brain-NA (basal ganglia); -0.23; 3.1 x 10<sup>-10</sup> |
| rs7439806 | G | A | intronic | 0.188 | 0.776 | 0.958 | 0.800 | 0.804 | 10.63 |  |  | Brain-NA (basal ganglia); -0.23; 3.1 x 10<sup>-10</sup> |
| rs55835604 | T | A | intronic | 0.191 | 0.776 | 0.958 | 0.800 | 0.804 | 13.81 |  |  | Brain-NA (basal ganglia); -0.23; 3.1 x 10<sup>-10</sup> |
| rs12510574 | G | A | intronic | 0.393 | 0.123 | 0.037 | 0.100 | 0.134 | 12.13 |  |  | Adipose - Subcutaneous; -0.16; 1.9 x 10<sup>-6</sup> |
| rs4861014 | A | G | intronic | 0.067 | 0.752 | 0.951 | 0.780 | 0.759 | 12.23 | 1.343 |  | Brain-NA (basal ganglia); -0.25; 1.2 x 10<sup>-12</sup> |
| rs10019356 | A | G | intronic | 0.176 | 0.789 | 0.960 | 0.820 | 0.811 | 15.41 |  |  | Brain-NA (basal ganglia); -0.24; 4.2 x 10<sup>-10</sup> |
| rs7660223 | C | T | intronic | 0.153 | 0.789 | 0.960 | 0.820 | 0.808 | 15.33 |  |  | Brain-NA (basal ganglia); -0.24; 4.2 x 10<sup>-10</sup> |
| rs11051 | G | A | 3' UTR | 0.051 | 0.751 | 0.951 | 0.780 | 0.758 | 10.81 | 1.343 |  | Brain-NA (basal ganglia); -0.25; 1.2 x 10<sup>-12</sup> |
| rs11935648 | G | A | 3' UTR | 0.176 | 0.788 | 0.960 | 0.820 | 0.808 | 1.44 |  |  | Brain-NA (basal ganglia); -0.24; 4.2 x 10<sup>-10</sup> |
| rs12511999 | C | T | 3' UTR | 0.087 | 0.751 | 0.951 | 0.780 | 0.764 | 0.24 | 1.302 |  | Brain-NA (basal ganglia); -0.26; 9.9 x 10<sup>-14</sup> |
| rs10938178 | G | C | 3' UTR | 0.067 | 0.751 | 0.951 | 0.780 | 0.759 | 1.32 | 1.343 |  | Brain-NA (basal ganglia); -0.25; 1.2 x 10<sup>-12</sup> |
| rs12512101 | C | T | 3' UTR | 0.086 | 0.751 | 0.951 | 0.780 | 0.764 | 4.31 | 1.302 |  | Brain-NA (basal ganglia); -0.26; 9.9 x 10<sup>-14</sup> |

<sup>a</sup>eQTLs described for *SLC30A9*. Extracted from GTEx database V7. NA: Nucleus accumbens. NES: Normalized effect size. Only the most significant tissue eQTL is shown. See multi-tissue eQTL comparison from GTEx in Figs S2-S19. See top 40 iSAFE values in Tables S2-S3.

We then took advantage of the previously inferred genome-wide genealogies available for the 1000 GP individuals and used Relate [29] to explore whether the lineages carrying the derived allele of the Met50Val substitution had spread at an unusually fast rate in comparison with competing lineages in the YRI, CHB and CEU populations. The marginal tree corresponding to the surrounding region of rs1047626 displayed the Val-derived variant in multiple lineages concentrated in CHB and CEU, although it was also present in some YRI haplotypes (Fig. S21). In contrast, most of the lineages carrying the ancestral variant were mainly found in YRI. The subsequent analysis with CLUEs estimated a drastic allele frequency shift compatible with the action of positive selection acting on rs1047626 at ∼56-77 kya (i.e., 2,000-2,750 generations before the present; using 28 years per generation) with selection coefficients s=0.0113 (logLR=5.68) and s=0.0017 (logLR=10.93) in CHB and CEU, respectively (Fig.1B). When analyzing other non-African populations such as FIN (European), JPT (East Asian), PJL (South Asian) and PEL (American), we obtained similar allelic trajectories and selection coefficients (Fig. S22). Thus, we seem to be detecting a strong signal of positive selection common to all non-African populations probably dating back before the expansion of modern humans though Europe and Asia.

### Potential signals of archaic introgression

We next explored whether such an early selection event could be explained by adaptive introgression. For that, we first investigated whether the Met50Val substitution was present in publicly available Neanderthal and Denisovan genomes. Remarkably, the Denisovan genome presents the homozygous genotype for the derived allele at the rs1047626 position, whereas three high coverage Neanderthal genomes (Altai, Vindija, and Chagyrskaya) are homozygotes for the ancestral allele. We then analyzed the corresponding Neanderthal and Denisovan base positions for a total of 170 SNPs found in high LD (r^2^>0.8) with rs1047626 in CHB and CEU and spanning a 98.74 kb region (Table S4). For 164 of these SNPs (96%), the most frequent allele in EAS, EUR, SAS, and AMR differs from the most prevalent in AFR, confirming the existence of two major contrasting Yin/Yang human haplotypes (i.e., two haplotypes differing at each polymorphic site; [30]). On the corresponding 98 covered SNP positions, the Neanderthal and Denisovan genomes were found to display higher allelic similarity and greater sharing of derived alleles with the allelic combinations most frequently found in non-Africans (Fig. S23; Table S4). Given these results, we next investigated whether this region had been identified in genome scans of Neanderthal and Denisovan introgression into modern humans. No hit was found within the most probable candidate genomic regions for Neanderthal introgression [31–33]. However, two independent studies recently reported two overlapping candidate regions for Denisovan introgression in Melanesians [34] and Papua New Guineans [35], respectively (Figs. S23-24; Table S4). The two most distant SNP positions where the archaic genomes were found to carry derived alleles shared with the most frequent allele combinations in present-day humans and coinciding with at least one of these introgression maps delimitate a region of 70.61 kb (chr4: 41,977,828-42,048,441; GRCh38). Consequently, we further explored the haplotype structure of this putatively introgressed region using all the SNP variants in the CHB, CEU and YRI populations from the 1000GP [36] together with those of Oceanian individuals available in the HGDP dataset [37] presenting a homozygous genotype in the Denisovan genome, and including the corresponding nucleotide positions in the three high-coverage Neanderthal genomes previously used (see details in Materials and Methods). The resulting set of 112 SNPs defined 4 archaic haplotypes and a total of 61 modern human haplotypes with two major allelic configurations differing at 66 SNP positions: htI, present in 53% of the YRI, 13% in CEU and 2% in CHB chromosomes, and ht II, found in 95% of the CHB, 67% of the CEU, and in 3% of the YRI chromosomes. Notably, the two most frequent haplotypes in Oceanians (ht LX and ht LXI found in 52% and 30% of the Oceanian chromosomes, respectively) only show 2 and 3 SNP differences when compared with ht II (Tables S5-6 and see haplotype network in Fig S25). We then used haplostrips [38] to better explore the haplotype structure of this region (chr4: 41,977,828-42,048,441; GRCh38) and genetic differences between modern human or Neanderthal haplotypes and the Denisovan genome. Notably, most of the Oceanian, CHB and CEU chromosomes (but also 8 out of 216 YRI chromosomes that carry the derived allele at rs1047626) displayed between 11 and 13 differences to the Denisova. In contrast, while the three Neanderthal haplotypes present between 17 to 28 differences to the Denisovan genome, the most common haplotype in YRI displays a total of 78 changes (Fig. 2, Table S7). Moreover, the most frequent allelic configuration in CHB and CEU contains 77 derived alleles shared with the Denisovan genome, while the most prevalent in YRI only presents 11 derived alleles shared with archaic humans (Fig. 2A, and Table S5). Similarly, the Neanderthal haplotypes, which all present the ancestral allele for rs1047626, share a minimum of 72 derived alleles with the Denisovan genome. To gain further insight into the genetic structure of such high-frequency Denisovan-like haplotype in modern humans and understand its presence in up to 7% of the YRI chromosomes, we repeated the same analyses including 4 additional Sub-Saharan African populations from the 1000 GP (i.e., ESN, MSL, GWD, and LWK; Fig S26, Table S8) and extending the explored region (chr4: 41,905,811-42,146,424; GRCh37; Fig S27, Table S9). Visual inspection of the corresponding haplostrips plots confirms that the Denisovan-like haplotypes carrying the derived Val allele in ESN (found in 4% of the ESN chromosomes), MSL (11%), GWD (10%) and LWK (8%) present the same archaic-like haplotype structure (and length ∼70,614 bp) found in CHB and CEU (see Figs. S26-27). By contrast, the African haplotypes first clustered to the Denisovan and Altai Neanderthal configurations but presenting the ancestral allele at rs1047626, display fewer shared derived alleles with the Denisovan genome (besides that of the rs1047626 position) and seem to present a much-dispersed pattern of private and shared derived alleles with both archaic genomes that expands outside the *SLC30A9* region and appears reshuffled in other African haplotypes (see Fig. S27, Table S9). As for the Denisovan-like modern human haplotypes containing the Met to Val substitution, since the probability of observing one haplotype of 70,614 bp in length shared by ancestry without being broken by recombination given the local recombination rate of the region (1.58 cM/Mb) and the time since divergence between modern and archaic humans (see details in Materials and Methods) is highly unlikely (p-value < 2.07 × 10^−8^ when considering a mutation rate µ = 1 × 10^−9^; Table S10), we can reject an scenario of incomplete lineage sorting (ILS) for causing the unusual haplotype configuration observed around the derived allele at rs1047626.

**Figure 2.**
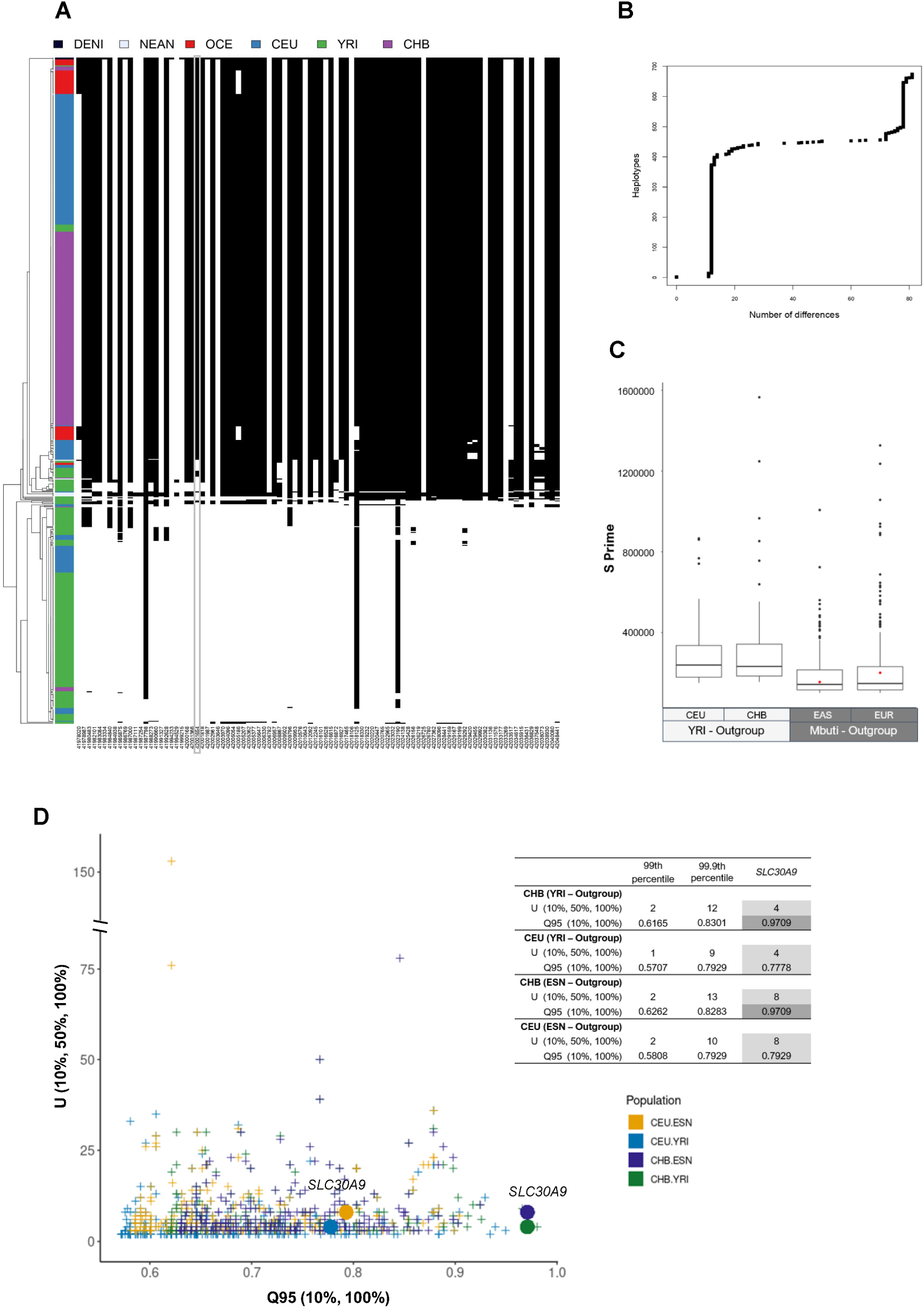
Potential signatures of adaptive introgression at *SLC30A9*. (A) Haplotypes along the putatively introgressed region (chr4: 41,977,828-42,048,441; GRCh38) clustered and sorted by increasing distance with the Denisova genome shown in the first row. Each column corresponds to a polymorphic position with a homozygous genotype in the Denisovan (112 in total; see details in Tables S4-5), each row is a phased haplotype, colored according to the population of origin of the individuals shown in the top. Note that all sites with a maximum within-population minor allele frequency below 0.05 were removed. Black cells represent the presence of the derived allele and white cells the ancestral allele (see Materials and Methods). The grey box highlights the location of rs1047626. (B) Number of SNP differences that the Neanderthal and modern human haplotypes present to the Denisovan genome sorted by decreasing similarity as obtained by Haplostrips [38]. (C) Genomewide distribution of S prime introgression values in Europeans and East Asians when using either YRI (values obtained from [41]) or Mbuti as an unadmixed group. The red dot denoted the *SLC30A9* region. (D) Joint distribution of the U (10%, 50%, 100%) and Q95 (10%, 100%) statistics in CEU and CHB when using either YRI or ESN as unadmixed group. Circles denote the U (10%, 50%, 100%) and Q95 (10%, 100%) values for the *SLC30A9* region in each corresponding analysis, which are also shown in the inserted table together with the 99.9th and 99th genomewide percentiles.

The presence of the putatively introgressed haplotype in YRI may explain why previous scans of Denisovan introgression have not detected *SLC30A9* as a candidate region for archaic introgression in Eurasians. Since Mbuti do not present the derived allele of the Met to Val substitution (Fig. S28) and probably have not been reached by the back to Africa migration [39,40], we next computed the S prime statistic [41] in the Europeans and East Asian samples available in the Human Genome Diversity Panel (HGDP-CEPH) [42], using the Mbuti as an outgroup, and detected the *SLC30A9* as a candidate for introgression in both groupings (Fig. 2C). Finally, we also investigated the scenario of archaic introgression in CHB and CEU with the Q95 and U statistics [43] using both the African YRI and ESN population as outgroups and considering the 10% outgroup cutoff to account for the known presence of this putatively introgressed haplotype in Africa (Fig. 2D, Table S11). In CHB, Q95_YRI, DEN_ (10%, 100%) and Q95_ESN, DEN_ (10%, 100%) detected unusual frequencies for the derived alleles that are shared with the Denisovan genome (and at <10% frequency in YRI or ESN, respectively) when considering the 99.9th percentile of the statistic computed genomewide; while in CEU, Q95_YRI, DEN_ (10%, 100%) and Q95_ESN, DEN_ (10%, 100%) were only significant when considering the 99th percentile (Fig. 2D). Similarly, the U statistic indicated an excess of shared alleles with the Denisova at more than 50% frequency in CHB and CEU (while being found at <10% frequency in YRI or ESN, respectively) only when considering the 99th percentile (Fig. 2D, Table S11, and see joint U and Q95 distributions indicating the *SLC30A9* region together with two previously described candidate regions for Denisovan introgression in CHB in Fig. S29 for comparison). Thus, overall, the numbers and frequencies of shared Denisovan alleles found in CHB and CEU provide further suggestive evidence for a likely scenario of adaptive introgression in the *SCL30A9* region.

### Signals of positive selection in Africa

As signatures of positive selection had been previously reported for the most abundant haplotype around *SLC30A9* in YRI [13], we also searched for putative candidate variants displaying unusual past allelic trajectories in Africa. Notably, several derived allele variants characterizing the African-like haplotype of *SLC30A9* were found to show marginal genome-wide p-values for positive selection with Relate (Table S2). After their corresponding functional annotation (Tables S13-16), we selected one of the strongest eQTLs (i.e., rs4861157 at chr4:42046074, GRCh38) that also presented one of the highest iSAFE values along the region for further analysis with CLUEs, which estimated a selection coefficient s=-0.00425 and a logLR=16.2401 for an adaptive event occurring within the last 1,000 generations (i.e., 28,000 years before present) in the YRI population (Fig. 1B, Figs. S30-33). Although the genotype for rs4861157 was not available in the high coverage archaic genomes and therefore it was not included in the computation of the haplotype network, we checked that its derived allele (A) is found in all chromosomes presenting either the most prevalent haplotype in YRI (ht I) or their corresponding derived haplotypes in the network (Fig. S25). These results confirm a pattern of contrasting directional selection between African and non-African populations acting on two major differentiated Yin/Yang haplotype configurations at *SLC30A9*. Moreover, as expected, the derived allele of rs4861157 has high frequencies in all African populations from the 1000 GP and associates with higher *SLC30A9* expression (Figs. S1B and S33).

### Molecular phenotype for Met50Val

We then investigated whether the Met50Val substitution in ZnT9 could determine any further differential molecular phenotype besides that of gene expression to explain this case of adaptative (positive) selection outside Africa. We first verified the predicted protein structure of ZnT9 using the AlphaFold algorithm. We observed that the polymorphic amino acid residue affected by rs1047626 is located at the cytosolic N-terminus domain of the ZnT9 transporter (Fig. 3A-B). It has been suggested that the N-terminus of the ZnT family modulates the transporter activity by facilitating either protein interactions or zinc binding [44,45]. Thus, we proceeded to analyze whether the Met50Val substitution could cause any differential phenotype on the ZnT9 protein expression levels, cellular location, or zinc homeostasis at cellular level. Accordingly, we used a heterologous expression system in HEK293 cells overexpressing each of the two ZnT9 forms resulting from the Met50Val substitution, carrying either the ancestral allele (ZnT9-50Met) or the derived allele (ZnT9-50Val). Our experiments showed no difference in protein expression between the ZnT9-50Met and ZnT9-50Val variants after their corresponding 24h transient expression (Fig. 3C). We next characterized the localization of the two ZnT9-50Met and ZnT9-50Val variants. For that, we performed immunostaining in non-permeabilizing and permeabilizing conditions. The results, shown in Fig. 3D, indicate that the ZnT9 N-terminus is located intracellularly and follows a reticular pattern independently of the overexpressed variant. We further explored the subcellular distribution of the variants co-transfecting the ZnT9 constructs with the FEMP probe, a double fluorescent reporter plasmid for the ER (in CFP) and the mitochondria (in YFP). Our colocalization analysis gave similar results for the two ZnT9 variants, both showing a higher colocalization degree with the ER compartment (Fig. S34). We then used super-resolution STED microscopy to confirm the presence of ZnT9 in mitochondria as described by others [21,22] (Fig. 4). ZnT9-HA was co-stained with anti-TOM20 (Fig. 4A), a mitochondrial outer membrane resident protein. Both ZnT9-50Met and ZnT9-50Val variants showed co-localization with mitochondria. Remarkably, by analyzing the mean distance between mitochondria (Fig. 4B) and the relative area occupied by mitochondria in the cell (Fig. 4C), we observed that the overexpression of ZnT9 generated mitochondrial clustering in comparison with non-transfected cells. To check whether the distance between ER and mitochondria might be altered we carried out FRET analysis using the FEMP probe together with the ZnT9 variants or empty vector. Our results showed no differences in any of the tested conditions (Fig. 4D). Overall, this initial characterization revealed no differences in cellular localization for either variant but suggests that ZnT9 has a role at the ER-mitochondria interface.

**Figure 3.**
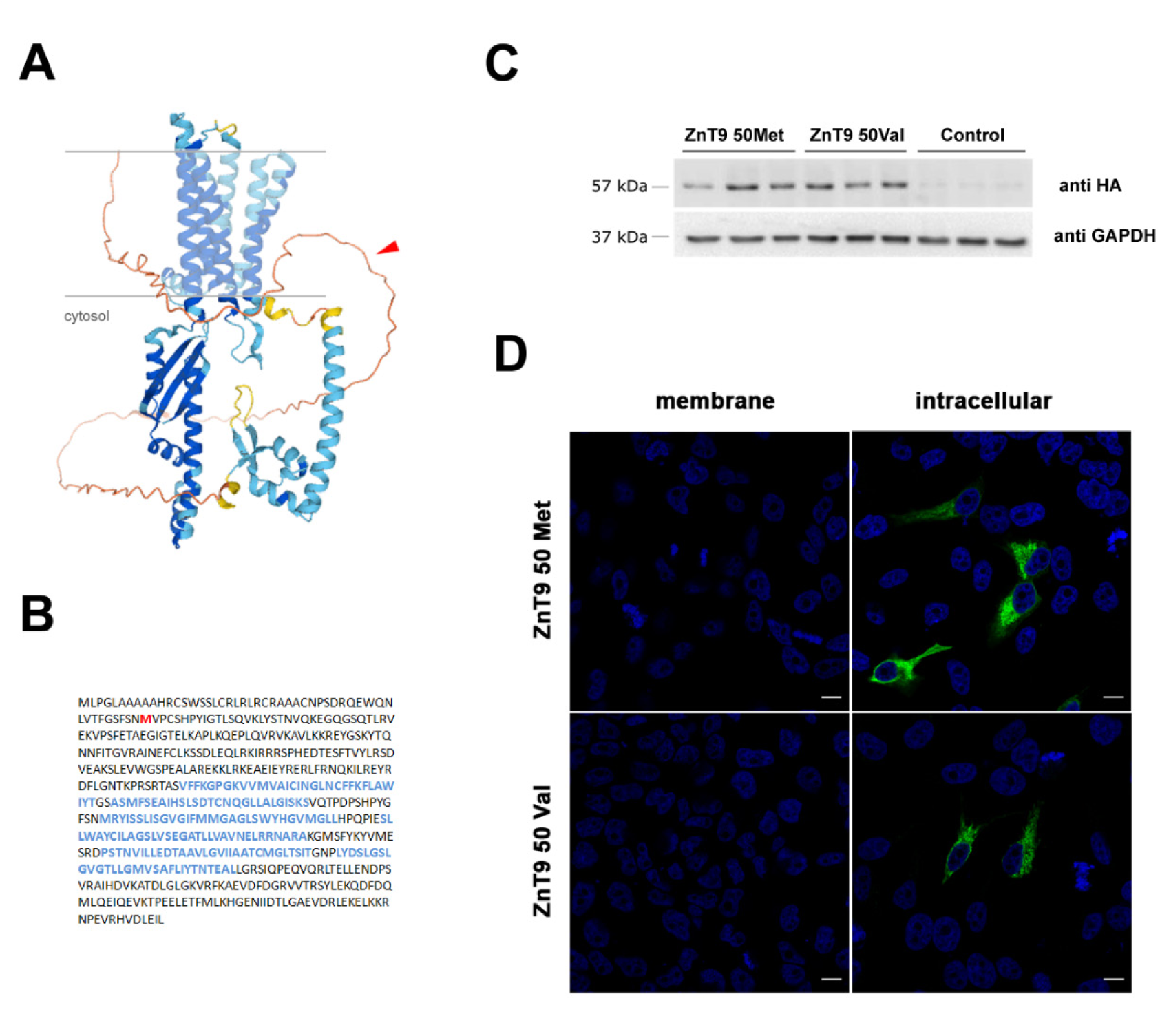
Characterization of the structure and overexpression of ZnT9 in HEK293 cells. (A) ZnT9 protein structure prediction by AlphaFold, showing 6 transmembrane domains and a cytosolic N-terminus, where the Met50Val substitution is indicated by a red arrow. (B) Protein sequence of the ZnT9 transporter with the Met50Val substitution in red and the predicted alpha-helices corresponding to transmembrane domains in blue. (C) Representative Western blot against HA and GAPDH in cells transfected with ZnT9-50Met, ZnT9-50Val, or an empty vector. (D) Representative pictures of the membrane and intracellular immunostaining of HEK293 cells expressing ZnT9-50Met and ZnT9-50Val isoforms (both in green). Nucleae were stained with DAPI (blue). Scale bar = 10 µm.

**Figure 4.**
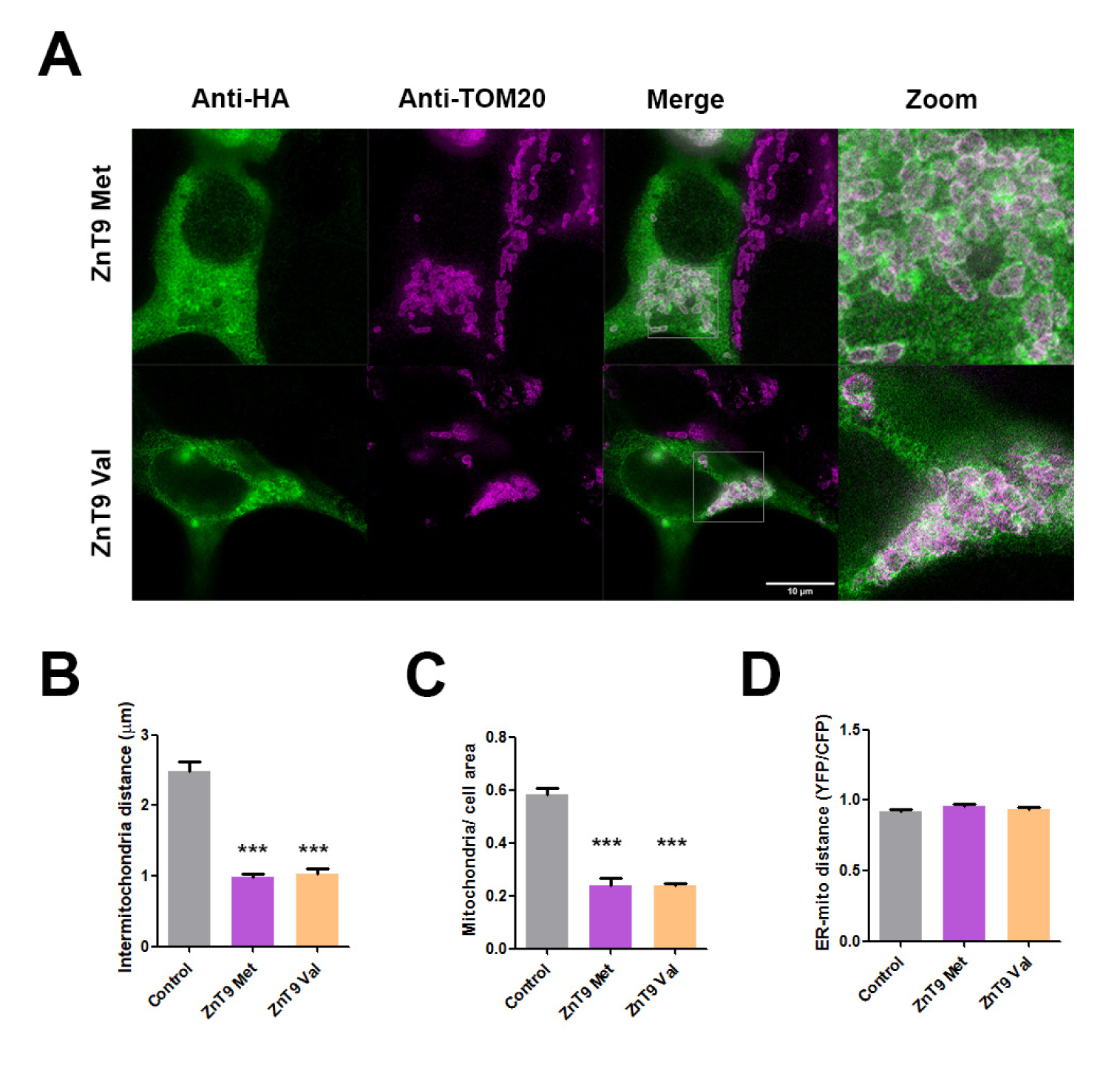
Subcellular localization of ZnT9 variants. (A) Superresolution STED microscopy in cells transfected with ZnT9-50Met and ZnT9-50Val immunoassayed with anti-HA (green) and with the mitochondrial marker anti-TOM20. Scale bar = 10 µm. (B-C) Bar graph representing mean inter-mitochondrial distance (B) and relative mitochondria area (C) in cells transfected with ZnT9-50Met or ZnT9-50Val (n=10). *** p<0.001 using Bonferroni test between conditions. (D) Bar graph measuring FRET between the endoplasmic reticulum and mitochondria transfected with an empty vector (control), ZnT9-50Met or ZnT9-50Val together with the FEMP probe (n=46-65). See further statistical details in File S1.

We then studied the impact of ZnT9 overexpression on cellular zinc homeostasis, comparing cells transiently transfected with either of the ZnT9 variants or an empty vector. First, the mRNA expression of the different ZIP transporters was characterized to analyze whether the overexpression of ZnT9, a transporter reported to promote cytosolic zinc efflux, is somehow compensated for by a modification in a zinc importer expression (Fig. 5A). Our results showed alterations in ZIP12 and ZIP14 expression, although in the case of ZIP12 CT values were above 30 and ZIP14 displayed a reduction in cells overexpressing either of the two ZnT9 variants. We then used flow cytometry with Zinquin to measure the cytosolic zinc content in basal conditions or with excess zinc for 30 min and observed no differences in cytosolic zinc handling between vectors in the two tested conditions (Fig. 5B).

**Figure 5.**
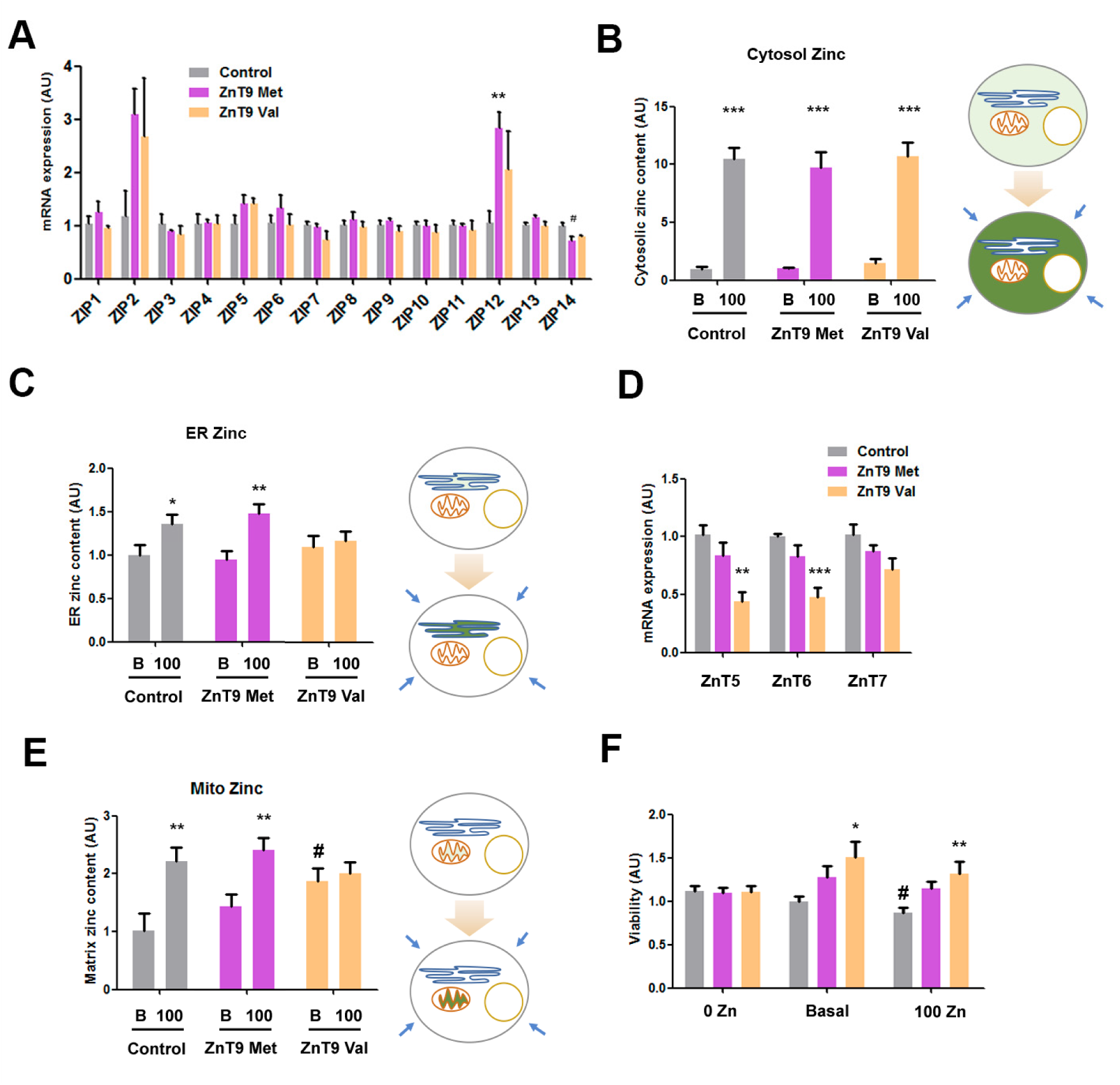
Characterization of the impact on cellular zinc homeostasis of ZnT9 variant overexpression in HEK293 cells. (A) RT-PCR comparing the expression of several ZTs in basal conditions in cells transfected with ZnT9-50Met, ZnT9-50Val, or an empty vector. 2^−(DDCT)^ plotted using GAPDH as the housekeeping gene (ZIP1, ZIP6, and ZIP7 n = 6; Rest of ZTs n = 3); ** p<0.01 versus control using t-test, # p<0.05 versus control using ANOVA with Bonferroni correction. (B) Evaluation of zinc content by flow cytometry using Zinquin in 10 µM and 100 µM of ZnSO_4_ (n = 9); *** p<0.001 using t-test. (C) Evaluation of endoplasmic zinc content using an endoplasmic reticulum fluorescent zinc sensor (ER-ZapCY1) in basal and 100 µM ZnSO_4_ conditions (n = 14-20). * p<0.05 using t-test, ** p<0.01 using t-test. (D) RT-PCR comparing the expression of several ZnT transporters in basal conditions in cells transfected with ZnT9-50Met, ZnT9-50Val, or an empty vector. 2^−(DDCT)^ plotted using GAPDH as the housekeeping gene (n = 5-6) ** p<0.01, *** p<0.001 using Bonferroni-corrected ANOVA. (E) Evaluation of zinc mitochondrial matrix content using Mito-cCherry-Gn2Zn incubating 40min with basal and 100 µM ZnSO_4_ conditions (n = 9-12); ** p<0.01 using t-test between basal and 100 µM zinc conditions, # p<0.05 using t-test between transfection conditions. (F) Viability MTT assay in cells transfected with ZnT9-50Met, ZnT9-50Val, or an empty vector incubated for 24h at 0, basal and 100 µM ZnSO_4_ conditions (n=8-15) * p<0.05, ** p<0.01 using ANOVA with Bonferroni correction between transfection conditions, # p<0.05 using Bonferroni-corrected ANOVA between zinc conditions. See further statistical details in File S1.

Given the presence of the ZnT9 transporter in ER membranes, we measured the endoplasmic zinc content using ER-ZapCY1 probe. No major alterations were observed in cells transfected with ZnT9-50Val, ZnT9-50Met, or an empty vector in basal conditions. However, upon incubation with 100 µM external zinc medium, only the cells transfected with ZnT9-50Met, or an empty vector had an increased ER zinc content, indicating that the overexpression of ZnT9-50Val was modifying the ER zinc homeostasis (Fig. 5C). When analyzing the RNA expression of the known ER zinc importers ZnT5, ZnT6 and ZnT7, we found that the overexpression of ZnT9-50Val causes a downregulation of ZnT5 and ZnT6 expression (Fig. 5D). Considering that ZIP7 is the major zinc exporter in the ER, we further explored its activity in conditions of ZnT9 overexpression. After 24h of transient expression, the results showed no difference in ZIP7 protein expression between the ZnT9 variants (Fig. S35A). We also measured the ER zinc content in the presence of a ZIP7 blocker, finding a strong increase in the fluorescence of the ER zinc reporter. However, differences between basal and 100 µM external zinc conditions were no longer observed, nor between ZnT9 variants and the empty vector (Fig. S35B), indicating that in the absence of ZIP7 activity the ER suffers a zinc overload.

As ZnT9 has been shown to regulate mitochondrial zinc homeostasis [21,22], we also interrogated the functional relevance of the Met50Val substitution by measuring the zinc content in the intermembrane space and the mitochondrial matrix, using SMAC-mCherry-GZnP2 and mito-Cherry-GZnP2 probes, respectively. In the intermembrane space, overexpression of ZnT9-50Val resulted in a slightly higher zinc content than cells overexpressing ZnT9-50Met or an empty vector. However, zinc increased equally in all conditions upon incubation with 100 µM zinc (Fig. S35C).

In the mitochondrial matrix, ZnT9-50Val-transfected cells had a higher zinc content in basal conditions as well but that was not further affected by incubating with 100 µM zinc, contrary to cells overexpressing ZnT9-50Met, or an empty vector (Fig. 5E). Given that mitochondrial zinc content is reported to depend on ER zinc transport [21,22], similar experiments were carried out, but this time blocking the ER zinc exporter ZIP7 for 30 min to accumulate zinc in this organelle. The result was an increase of zinc levels in the mitochondrial matrix of control cells but not in ZnT9 overexpressing cells (Fig. S35D). Moreover, the accumulation of zinc in the mitochondrial matrix of ZnT9-50Val-transfected cells is not restricted to HEK293. We observed similar results in the myoblast cell line C2C12 (Fig. S35E). As a summary, we have generated a schematic model with the main findings related with the impact ZnT9 variants on intracellular zinc homeostasis (Fig. S35F). In order to gain further insight into the ZnT9 transport activity in the mitochondria, ZnT9 was silenced with siRNA and the zinc content of the mitochondrial matrix was measured. As the data showed that incubation with 100 µM zinc increases zinc levels despite silencing ZnT9, we discarded the possibility that ZnT9 is a major zinc importer in the mitochondria (Fig. S36A). Our expression data confirmed the silencing strategy and showed no alterations in the expression of ZnT5, the zinc importer in the ER, when ZnT9 was silenced (Fig. S36B). Finally, we also investigated the impact of the alteration of zinc homeostasis caused by the ZnT9 variants overexpression on mitochondrial metabolism. MTT experiments were carried out in cells incubated for 24h with 0, basal and 100 µM zinc content. Higher metabolic activity signal was found in ZnT9-50Val-overexpressing cells both in basal and zinc excess conditions compared with the control (Fig. 5F). Considering that at basal conditions cell number was not altered after 24h transfection with different plasmids (Fig. S37), MTT assays are revealing differences in mitochondrial metabolism. The absence of difference in the 0 zinc media supports the idea that the impact on metabolism is due to a zinc-dependent process.

### Exploring an adaptive phenotype for Met50Val

As three SNPs in the 3’ flanking region of the *SLC30A9* gene (rs2880666, rs6447133, rs7659700) have been previously identified as QTLs influencing zinc content in the liver [23] and they are found in moderate linkage disequilibrium (0.66 < r^2^ < 0.76) with rs1047626 (Table S17), we examined whether rs1047626 could be associated with such a systemic phenotype. No significant differences in liver zinc content were detected between individuals carrying the A (n=74) or G alleles (n=212; p-value=0.107), nor between the AA (n=11), AG (n=52), and GG (n=80) genotypes at rs1047626 (p-value = 0.322). However, in agreement with the known effects of the surrounding nutriQTLs with which the alleles at rs1047626 associate, the homozygous individuals for the G allele showed a tendency towards higher zinc concentrations in the liver than those homozygous for the A allele, whereas the heterozygotes AG were associated with an intermediate zinc content (Table S18, Fig. S38).

Additionally, we searched for GWAS annotations linked to any of the 170 SNPs in high linkage disequilibrium (r^2^ > 0.8) with rs1047626. We found reported GWAS hits for helping behavior (rs1507086, rs2660319, rs11051), total PHF-tau SNP interaction (rs4861153), and one SNP (rs34215985) associated with multiple psychiatric phenotypes, such as anorexia nervosa, hyperactivity disorder, autism spectrum disorder, bipolar disorder, major depression, obsessive compulsive disorder, and schizophrenia (Table S4). A phenome-wide association study (PheWAS) for rs1047626 retrieved seven entries with a corrected p-value ≤ 1.5 × 10^−5^ (0.05/3,302 unique traits), including traits within the psychiatric (i.e., neuroticism and major depressive disorder, being the derived G allele of the rs1047626 polymorphism the allele that increases the risk of both phenotypes), metabolic (i.e., impedance, with the G allele associated with lower impedance measures), activities (i.e., fish oil, and glucosamine uptake, with the G allele increasing the phenotype value) and skeletal (i.e., height, with the G allele decreasing the phenotype value) domains, respectively (Table S19).

## Discussion

The motivation for this study was the previous detection of strong signatures of positive selection in the *SLC30A9* region [13,14,16,17,20,25]. In this context, we first re-examined the genetic evidence for a hard-selective sweep using the F_ST_, XP-EHH, and Fay and Wu’s H selection statistics and found several functional variants on *SLC30A9* that could explain the strong adaptive signals detected. In accordance with Zhang et al. (2015) [13], we confirmed contrasting signals of positive selection between two major haplotypes in Africa and East Asia, the latter also being present at intermediate-high frequencies in South Asia, Europe, and America. The major African haplotype carries alleles at several eQTLs associated with higher *SLC30A9* expression. In contrast, the most prevalent allelic combinations outside Africa are associated with lower gene expression and are linked to the derived allele of the rs1047626 polymorphism, which causes a Met to Val substitution at ZnT9 codon position 50. Although no experimental characterization of its corresponding molecular phenotype has been reported to date, previous studies have attributed the strong signals of adaptation found in East Asians to this non-synonymous variant [13,17,23]. In this work, we describe the presence of the derived Val allele in the Denisovan genome and an unusually high sharing of derived alleles between the major haplotype outside Africa and archaic humans (both Neanderthals and Denisovans). Moreover, we demonstrate that the persistence of such similar allelic configuration around the Met50Val substitution comprising a segment of 70,614 bp in length between modern and archaic humans cannot be explained by ILS when considering the recombination rate of the *SLC30A9* region. These results are compatible with two independent studies reporting a Denisovan adaptive introgressed region in modern Melanesians [34] and Papua New Guineans [35] overlapping with the *SLC30A9* gene and comprising the rs1047626 polymorphism. Interestingly, the putatively introgressed Oceanian chromosomes not only present the derived allele at rs1047626 but display haplotypes just 2-3 mutational steps away from those positively selected in CHB and CEU. However, some African chromosomes (4% in ESN, 7% in YRI, 8% in LWK, 10% in GWD, and 11% in MSL) carry the derived rs1047626 allele and present similar allelic configurations to those suggested as putatively introgressed. The presence of such Denisovan-like haplotypes in Africa could result from back-to-Africa migrations [46] and, as shown by our S prime analysis when using Mbuti as unadmixed group, it might have easily prevented the detection of the *SLC30A9* gene in previous maps of archaic introgression in modern humans. Similarly, the use of the U and Q95 introgression statistics with the 10% unadmixed outgroup frequency cut-off confirmed that the *SLC30A9* region displays unusual sharing of derived alleles with the Denisova as well as unusual numbers of shared Denisovan alleles at frequencies >50% in both CHB (above the 99.9th genomewide percentile for Q95) and CEU (when considering the 99th percentile), suggesting again a likely scenario for adaptive introgression. However, we also describe high similarity (but greater SNP differences) between the Denisovan genome and (at least two) Neanderthal haplotypes and other much less frequent modern human haplotypes carrying the ancestral allele at rs1047626. But while the Denisovan-like haplotypes carrying the derived allele at rs1047626 present very similar specific allelic configurations and lengths across populations, these other archaic-like configurations are more diverse and appear reshuffled in different chromosomal backgrounds, especially in Africa. In the future, the availability of new high-coverage archaic genomes and the consideration of multiple evolutionary event models may help to better explain such unusual sharing of archaic-like haplotypes in the *SLC30A9* region and test the likelihood of adaptive introgression from either Denisovans, Neanderthals, or other archaic humans. For example, although we have rejected the possibility of ILS, we cannot discard a scenario of shared ancestral variation coupled with two contrasting selecting sweeps in and outside Africa creating such an unusual putative Denisovan-like configuration outside Africa. Similarly, we cannot discard other more complex scenarios including a first introgression event from the ancestors of modern humans into archaic humans followed by a second archaic introgression event into non-Africans and back to Africa migrations, as similarly suggested to explain the finding of Neanderthal introgression sequences in the 1000 GP African populations or the presence of a putative modern human haplotype at the *KNL1* gene in some Neanderthals [47,48].

The protein structure of ZnT9 predicted by AlphaFold [49] shows a total of 6 transmembrane domains and a cytosolic N-terminus for ZnT9, a structure similar to that predicted for the other members of the ZnT family. The location of the Met50Val substitution in the N-terminus may indicate a putative effect of this non-synonymous SNP on zinc homeostasis through a possible accessory role of the N-terminus domain of ZnT9 in facilitating zinc transport or protein interactions. When exploring the consequences of the Met50Val substitution at the molecular level, the protein expression analyses revealed a similar partial localization of both ZnT9 variants in the ER and mitochondria. Moreover, we observed mitochondrial clustering near the ER when ZnT9 was overexpressed what could imply a specific role in mitochondria-ER contact sites and architecture. The presence of the transporter in both cellular compartments has been previously described [50], and the ionic fluxes between the two organelles are highly interconnected. In this context, it has been recently described that the ZnT9KO phenotype can be rescued by targeting transporters in the ER and mitochondria [22]. The ER plays a major role in protein folding, maturation, quality control, and trafficking. Many of these processes require zinc to function correctly, and its deficit or excess often leads to protein malformations that induce ER stress. The most well-known ZTs implicated in ER zinc homeostasis are the ZnT5-ZnT6 heterodimer and ZnT7 homodimer, which are involved in zinc importation [51,52], and ZIP7, which is the main zinc-releasing transporter in the ER [53]. Considering the effect on ER zinc homeostasis of the overexpression of either ZnT9 isoform, we propose that the 50Val variant avoids ER zinc accumulation due to the reduced ZnT5-ZnT6 heterodimer expression. Additionally, we observed that ZIP7 inhibition generates an excess of zinc in the ER, supporting the idea that the zinc exporting activity of ZIP7 is key for maintaining endoplasmic zinc homeostasis. Therefore, it is likely that the observed absence of differences in ER zinc content in basal conditions is due to the master role of ZIP7 in fine-tuning ER zinc content.

It has been proposed that ZnT9 in the mitochondria together with ZnT5 in the ER determine the zinc content in the mitochondrial matrix [22]. ZnT9 has been identified as a zinc exporter from the mitochondria based on the zinc accumulation observed in the mitochondria in ZnT9KO models [21,22,50]. We only detected a mild increase in zinc in ZnT9-silenced cells, which was not statistically significant. Probably, the zinc accumulation observed in KO models is the result of a complete and chronic effect. Nevertheless, this transport model implies that ZnT9 works in the opposite direction from the rest of the ZnT transport family. Our results showed that ZnT9 50Val expression increases the zinc content of the mitochondrial matrix but avoids zinc overload. The intermembrane space, however, showed no differences at high zinc concentrations between the tested conditions. Zinc homeostasis in this space has been previously reported to follow cytosolic zinc concentrations [54]. On the other hand, it is likely that the repression of ZnT5 expression observed in ZnT9 50Val cells might be a negative feedback to avoid a further zinc increase in the mitochondrial matrix. Having ruled out that ZnT9 is the main mitochondria zinc importer, we speculate that it might influence a wider machinery of proteins involved in mitochondrial zinc homeostasis. In this respect, ZnT9 has been recently connected to the expression of mitochondrial ribosomes and OxPhos components [55]. Importantly, we show that the ZnT9 50Val variant increases mitochondrial metabolism in a zinc-dependent manner. ZnT9 has been previously reported to affect mitochondrial function [21,22] and in this work we show that it also affects zinc homeostasis in the ER. Both organelles are interconnected and essential for cell fate and metabolism [56].

Previously, a mutation in the fourth TMD in *SLC30A9* was found to cause acute dysregulation of zinc homeostasis in all family members affected by a novel cerebro-renal syndrome with important early neurological deterioration [57]. Given that the Met50Val substitution sits in what is expected to be an auxiliary region for the activity of the transporter, we hypothesize that the effects of rs1047626 on zinc homeostasis are milder, but somehow became adaptive in the past. Accordingly, when analyzing whether the genotype for rs1047626 influenced zinc content in the liver as a proxy for systemic zinc homeostasis, we only detected a tendency towards increased zinc in the liver when the derived G allele (50Val) was present. Even if larger sample sizes might produce results with significant differences, any putative effect of rs1047626 on liver zinc content would be lower than that of the QTLs previously identified in the 3’region of the *SLC30A9* gene. Although *SLC30A9* is moderately expressed in most tissues, its expression is higher in the fetal brain, cerebellum, skeletal muscle, pituitary, thyroid, and kidneys [57]. Hence, greater effects of the Met50Val substitution on other tissues or organs cannot be ruled out, for instance, related to adjustments in renal excretion, brain function or even skeletal muscle metabolism. The major haplotype outside Africa was found significantly associated with greater susceptibility to major depression and other related psychiatric disorders [58,59] as well as with higher values for self-reported helping behavior [60]. Moreover, our PheWAS analysis confirms that the derived G-allele of rs1047626 contributes not only to a greater risk for both major depressive disorder and neuroticism but also lower impedance (i.e., a higher lean body mass), among others.

In the last years it is emerging the crucial function of ZnT9 in core mitochondrial processes. The function of the mitochondrial ribosomes (mitoribosomes) and, consequently, the expression of proteins encoded by mitochondrial DNA, such as several OxPhos subunits, has been shown to depend on ZnT9 expression [55]. In this context, our data proves that the Met50Val substitution influences mitochondrial metabolism. This effect might have an impact on systemic metabolism that could explain the association of rs1047626 with higher lean body mass. However, despite all the data we have compiled, it is not obvious how the observed differential molecular phenotype for the Met50Val substitution can be translated into an adaptive phenotype at the organism level or might relate to environmental zinc [13]. Given the essential role of the mitochondria in skeletal muscle thermogenesis, we propose that the derived G allele, which we have shown provides better protection to the mitochondria, may have been positively selected to facilitate adaptation to cold while providing higher susceptibility to neuropsychiatric traits. This is a plausible scenario for a case of archaic adaptive introgression, as Denisovans and Neanderthals were probably already well adapted to the local environmental conditions modern humans encountered when expanding across Eurasia.

Some limitations in this study should be noted. Although several promising human traits have been associated with rs1047626, further work is required to understand how the molecular phenotype of differential zinc handling between the mitochondrial and the endoplasmic reticulum described here for the Met50Val substitution is refined into an adaptive phenotype at the organismal level. Similarly, we note that it cannot be ruled out the possible adaptive role of regulatory variants, influencing both *SLC30A9* expression and liver zinc content (and thus systemic homeostasis of zinc), also contributing to the adaptive phenotype we proposed for the Met50Val substitution. After understanding where in the cell ZnT9 activity is having an impact and demonstrating that there are zinc cellular homeostasis differences between the Met50Val ZnT9 variants using an overexpression approach, other experimental assays working with endogenous expression ZnT9 levels might help to better characterize the putative disruption role of the Met50Val substitution in particular cell types and tissues of interest, such as muscle and adipose tissue or neuronal cells, among others. Finally, additional work is necessary to ascertain the exact timing, place, and potential admixture events experienced with archaic humans to explain the current patterns of variation and contrasting signatures of positive selection observed along the *SLC30A9* gene region among modern human populations.

## Materials and Methods

### Signals of positive selection in the SLC30A9 region

The PopHuman genome browser (http://pophuman.uab.cat) [26] was consulted to search for signatures of recent positive selection along the *SLC30A9* region. This browser contains a broad range of genomic metrics and neutrality tests for all 26 human populations of the 1000 Genomes Project (1000 GP) [36]. Window-based computed values for F_ST_, XP-EHH, and Fay and Wu’s H statistics were extracted from three geographically differentiated populations from Africa, Europe, and Asia: the Yoruba from Ibadan (YRI), Utah residents with Northern and Western European ancestry (CEU), and Han Chinese from Bejing (CHB). Gene annotations surrounding the *SLC30A9* region were obtained from the Ensembl genome browser (http://grch37.ensembl.org/). The collected information was then represented using the Gviz package [61]. For each selection statistic and population, deviations from neutrality were considered as those greater than two standard deviations from the genomic mean.

### Analysis of putative adaptive variants in SLC30A9

All SNPs for the longest *SLC30A9* gene transcript (ENST00000264451) were extracted from the Ensembl Genome Browser (GRCh37 assembly). Only biallelic variants with a MAF ≥ 0.02 were kept for subsequent analysis and annotated using ANNOVAR [62] to obtain their corresponding genomic location, SNP type classification (i.e., coding, non-coding, synonymous, non-synonymous), and allele frequency information. All compiled variants were explored for associations in the GWAS Catalog (v1.0) and eQTLS in the GTEx Portal Dataset (V7) and they were further annotated with different *in silico* function predictors such as the score of Combined Annotation-Dependent Depletion (CADD score) [27], the Eigen score [63], and the FitCons score [64]. SNPs in high linkage disequilibrium (r^2^>0.8) with rs1047626 in CEU or CHB and their corresponding allele frequencies by continental region (AMR, EAS, EUR, SAS and AFR) were extracted from Ensembl. We run iSAFE [28] using a 300 kb region centered on rs1047626 (chr4: 41,853,671-42,153,671; GRCh37) from the CHB, CEU and YRI phase 3 1000 GP sequencing data and inferring the corresponding human ancestral allelic states from the Ensembl EPO multiple alignments.

### Analysis of positive selection on candidate variants

We used Relate (https://myersgroup.github.io/relate) [29] to test whether the derived alleles at rs1047626 and rs4861157 were detected as fast spreading variants along multiple lineages when compared to competing non-carriers. For that, we first obtained the Relate-estimated coalescence rates, haplotypes and genealogies available for the 1000 GP (kindly provided by Leo Speidel), removed all sample haplotypes not belonging to CEU, CHB or YRI (with RelateFileFormats -- mode RemoveSamples) and annotated SNPs as recommended in order to extract trees (with RelateFileFormats --mode GenerateSNPAnnotations) and the inferred genealogies for rs1047626 and rs4861157 in CHB, CEU and YRI (with RelateExtract --mode SubTreesForSubpopulation -- years_per_gen 28), which were then visualized (with RelateTreeViewMutation -m 1.25E-8). Subsequently, we ran CLUES [65] to estimate the corresponding selection coefficients and allelic trajectories in either CEU and CHB or YRI using the SampleBranchLengths output from Relate as obtained with the Inference.py with N 30.000. The same procedure was repeated to run CLUES on FIN, PEL, JPT, and PJL.

### Introgression

Allelic states in the Neanderthal and Denisovan genomes for those SNPs displaying high linkage disequilibrium (r^2^>0.8) with rs1047626 in CEU and CHB were queried from the genotypes available in four high coverage archaic genomes (https://cdna.eva.mpg.de/neanderthal/). We initially inferred as putatively introgressed the genomic region comprised between the two more distant SNP positions found at high linkage disequilibrium (r^2^>0.8) with rs1047626 in CEU and CHB where the archaic genomes carried derived alleles shared with the most frequent allele combinations in present-day humans while also coinciding with at least one of the Denisovan introgression maps available for Melanesians [34] and Papua New Guineans [35]. To further explore a scenario of archaic introgression, all SNP genotypes along this putatively introgressed region (hg 38, chr4: 41977828-42048441) were extracted for all the Oceanians and for the CHB, CEU, and YRI populations from the corresponding VCF files of the HGDP [37] and 1000GP [66] sequencing projects, merged, and phased with Shapeit4 [67] using the recombination maps available (https://github.com/eyherabh/genetic_map_comparisons). In turn, VCF files for the Denisova, AltaiNeandertal, Vindija33.19, and Chagyrskaya-Phalanx archaic genomes were downloaded (https://cdna.eva.mpg.de/neanderthal/), lifted over to hg38, and merged. We then crossed all the Denisovan positions found with a homozygous genotype in the putatively introgressed region with the corresponding Neanderthal positions and phased genotypes across the YRI, CHB, CEU and Oceanian populations. Only three positions in the Altai Neanderthal were found as heterozygotic positions and were kept in the analysis. From this complete set of inferred modern and archaic haplotypes (65 in total defined by 112 polymorphic positions, see Table S5), we then computed nucleotide pairwise distances and plotted the haplotype network using the ape and pegas R packages [68]. The probability of observing a segment of 70,614 pb in-length shared with archaic humans due to incomplete lineage sorting (ILS) was calculated following the same procedure as in Huerta-Sánchez et al. (2014) [69] considering the generation (g=25 years) and split times as in Dannemann et al. (2016) [70] and the average recombination rate of the putatively introgressed *SLC30A9* segment (r = 1.58 × 10^−8^; extracted from https://github.com/eyherabh/genetic_map_comparisons). Briefly, we first calculated which is the expected length (L) for a segment shared by ILS using the formula L = 1/(r(t1+t2)/g) considering the split times for the divergence between humans and hominids (t1) and between Denisovan and Neanderthals (t2) given in Prüfer et al. (2014) [71] when considering two different mutation rates. Subsequently, the probability of observing an archaic segment of 70,614 bp in-length that persists due to ILS was computed as 1-GammaCDF (observed length, k, I/L), being 1/L the rate of the gamma distribution, L the expected length and k, the shape (k=2) (see details in Table S10). S prime was computed pooling all the whole genome sequences for the European and East Asian populations available at the HGDP dataset [42], respectively, and considering Mbuti as unadmixed outgroup. Singletons were excluded from the S prime analysis, and we only used biallelic sites without missing variants. S prime values estimated for CHB and CEU were directly extracted from Browning et al. (2018) [41]. U and Q95 statistics [43] testing for archaic introgression in CHB and CEU (phase 3 1000 GP sequencing data) were computed twice in the form U (10%, 50%, 100%) and Q95 (10%, 100%) using either YRI or ESN as outgroup (phase 3 1000 GP sequencing data), respectively, and considering only biallelic sites with no missing data. Significance was evaluated considering the 99th and 99.9th percentiles obtained from a genomewide distribution of the corresponding U and Q95 statistics computed in 40 kb windows as in Racimo et al. (2017) [43]

### GWAS analysis

We used the GWAS catalog (https://www.ebi.ac.uk/gwas/) [72] and the PheWAS option in the GWAS atlas (https://atlas.ctglab.nl/) [73] to look for trait associations with rs1047626 and SNPs in high LD (r^2^ >0.8). In the PheWAS analysis, a total of 3,302 unique traits as available at the GWAS Atlas database (last accession on 16/05/2022) were considered for Bonferroni multiple test correction (FDR=1.5 × 10^−5^).

### Generation of polymorphic sites

The plasmid containing the human *SLC30A9* gene codifying for the ZnT9 transporter with an N-HA tag was obtained from Sino Biological Inc. (Catalog nr. HG22621-NY). The Met50Val polymorphism was generated via site-directed mutagenesis following standard conditions (QuikChange Lightning; Agilent) with the following primers: forward primer (5’-GACATTTGGAAGCTTTTCAAACGTGGTTCCCTGTAGTCA-3’) and reverse primer (5’-TGACTACAGGGAACCACGTTTGAAAAGCTTCCAAATGTC-3’). Before being used for cell transfection, both ZnT9-50Met and ZnT9-50Val isoforms were confirmed by sequencing. The LightRun sequencing service of Eurofins Genomics was used following the standard conditions for purified plasmid DNA samples with the following primers: forward primer 5’-GGCACAGAACTCAAAGCT-3’ and reverse primer 5’-TCTTCATGGGGACTTCGT-3’.

### Genotyping of DNA liver samples

DNA samples and information regarding zinc concentration in the liver were obtained from the project of Engelken et al. (2016) [23], which determined micronutrient concentrations in liver samples from healthy individuals of western European origin, originally gathered by the UKHTB (for details, see Engelken et al. (2016) [23]). DNA concentrations for 143 available samples from the original study were quantified using a NanoDrop One spectrophotometer (Thermo Fisher Scientific, Waltham, MA, USA). A genomic region of 226 bp comprising the rs1047626 position was amplified, purified, and sequenced using the LightRun sequencing service (Eurofins Genomics) following standard conditions for purified PCR products. The primers used for amplification were: forward primer 5’-GAAGCAGTGAAACACCTCTGG-3’, reverse primer 5’-TGTTTGTGATCCCTGTCCTTC-3’. For sequencing we used primer 5’-TGCAGCTAGGACTTGGTTTG-3’.

### Cell culture and transfection procedure

HEK293 and C2C12 cells were cultured in DMEM supplemented with 10% FBS, 1% L-glutamine, and 1% penicillin and streptomycin (Basal medium) at 37 °C in a humidified 5% CO_2_ atmosphere. ZnSO_4_ was added as needed to the final medium to generate specific Zn^2+^ concentration conditions. Cells were transiently transfected with ZnT9-50Met, ZnT9-50Val, pEGFP, or empty pCDNA3 vectors depending on the experiment. Polyethyleneimine (PEI) was used as the transfection reagent using 3 μg DNA for a 6-well plate or 1 μg DNA for a 24-well plate. In RNA interference experiments we used siRNA control (1027310, Qiagen) and siRNA ZnT9 (114789, Eupheria Biotech), and the transfection reagent was Lipofectamine 3000. Cells were incubated with the transfection solution for 3 hours, then washed and replaced with normal media. The experiments were performed 24h after transfection. To characterize the genotype of cells used for all the experiments, the DNA of HEK293 cells was extracted and purified following the NucleoSpin Tissue protocol (Macherey-Nagel, Düren, Germany). The genotype at rs1047626 was verified to be heterozygote (AG) in HEK293 cells with the same sequencing procedure previously described for the liver DNA samples.

### Western Blotting

Transfected cells were grown in 6-well plates and incubated for 24 hours. Cell lysis to detect ZnT9 protein was performed with 30 μL of lysis buffer containing 50 mM Tris-HCl at pH 7.4, 150 mM NaCl, 0.5% Nonidet P-40, and EDTA-free protease inhibition cocktail (Roche). Cell lysates were vortexed for 30 min at 4 °C and centrifuged at 10,000× *g* to remove aggregates and boiled for 5 min at 95 °C to lastly be placed on ice for 1 min. After electrophoresis in 12.5% polyacrylamide gel, proteins were transferred to nitrocellulose membranes using the iBlot system (Invitrogen, Waltham, MA, USA). Membranes were blocked with either 5% milk in TBS-Tween 0.1%, for GAPDH, or 5% BSA for HA-ZnT9, for 1 hour at room temperature. Primary antibodies were diluted in the corresponding blocking solution anti-HA for ZnT9 (H3663, Sigma-Merck), and anti-GAPDH (ab8245, Abcam). HRP secondary antibodies (1:1000; GE Healthcare) were used depending on the primary antibody. The ChemiDoc XRS+ system (Bio-Rad, Hercules, CA, USA) was used to obtain high-quality images. Quantity One Software (Bio-Rad) was used to analyze the results.

### Immunostaining

For immunodetection of the two isoforms of ZnT9, for both cell surface and total cell expression, immunostaining was performed 24 hours after transfection with ZnT9-50Met or ZnT9-50Val plasmids in 24-well plates with coverslips coated with collagen. For cell surface expression, cells were incubated with anti HA (1:1000) in DMEM for 1h at 37°C. All samples were fixed with 4% paraformaldehyde (PFA) and only for total cell expression experiments were cells permeabilized with 0.1% Triton x-100 in PBS for 10 minutes. For all experiments, samples were blocked overnight with 1% BSA and 2% FBS in PBS. For total cell expression experiments, samples were incubated for 1h with an anti-HA antibody (1:1000 in blocking solution). For all experiments, cells were incubated with the secondary antibody (1:2000 in blocking solution), a goat anti-mouse Alexa Fluor 488 (Molecular Probes). Images were acquired using an inverted Leica SP8 Confocal Microscope with a 63× Oil objective and analyzed using ImageJ software.

In the colocalization analysis, to determine the subcellular distribution of the variants, immunostaining was performed 24 hours after transfection in 24-well plates with coverslips. Cells were co-transfected with ZnT9-50Met or ZnT9-50Val plasmids and a modified FRET-based ER-mitochondria proximity probe named FEMP [74]. The immunostaining procedure was the same as the one described above, except for the secondary antibody, which in this case was a goat anti-mouse Alexa Fluor 555 (Molecular Probes). The analysis of colocalization of ZnT9 with either the ER or mitochondria was performed with both Pearson’s correlation coefficient and Mander’s overlap coefficient using ImageJ software.

For STED imaging, cells were grown and seeded on 1.5H-thickness cover glasses. Primary antibodies were diluted in 2% BSA blocking buffer reagent. We used the rabbit anti-human TOM20 and the rabbit anti-human Calreticulin antibodies (1:1000) and secondary antibodies Abberior STAR RED or ORANGE (1:350). STED images were taken with a commercial Leica TCS SP8 STED 3× microscope equipped with a pulsed supercontinuum white light laser excitation source, using a 100× 1.4 NA oil HC PL APO CS2 objective. Post-analysis was performed on ImageJ. To measure the area occupied by the mitochondria with the Freehanded selection tool, we measured the cell region on the HA channel, subtracted the nucleus area and selected the area occupied by these organelles. The distribution of the mitochondria was quantified tracing a segmented line that connected all single mitochondria in one cell and then compare the length of these lines against the length in control cells.

### Real-Time RT PCR

Cells transfected with either ZnT9-50Met, ZnT9-50Val, or empty pCDNA3 vectors were grown in 6-well plates and incubated for 24 hours. Total RNA was extracted from cells using Macherey-Nagel total RNA extraction kit. RNA was measured using a NanoDrop 1000 spectrophotometer (Thermo Fisher). cDNA was generated using a SuperScript Reverse Transcriptase system (Invitrogen). Quantitative PCR was performed using SYBR Green (Applied Biosystems) in the QuantStudio 12K system (Applied Biosystems). Primers are listed in Table S20.

### Zinc Measurements

For the detection of zinc at the cytosolic level, cells were co-transfected with ZnT9-50Met or ZnT9-50Val vectors and pEGFP in 24-well plates. 24 hours after transfection, cells were incubated with 25 µM of Zinquin (Sigma-Aldrich, Darmstadt, Germany) for 30 min at 37 °C (5% CO_2_) in isotonic solution (ISO) containing 140 mM NaCl, 2.5 mM KCl, 1.2 mM CaCl_2_, 0.5 mM MgCl_2_, 5 mM glucose, and 10 mM Hepes (300 milliosmoles/liter, pH 7.4), and different concentrations of Zn^2+^. Cells were then dissociated with Trypsin 0.05% in 0.53 mM EDTA and washed with PBS. Fluorescence was quantified using an LSRII flow cytometer. Further analysis was performed using Flowing software to quantify Zinquin fluorescence of live transfected cells (Perttu Terho, Turun yliopisto, Turku, Finland).

To determine ER zinc levels, *in vivo* confocal imaging was used in cells co-transfected with the FRET-based ER-ZapCY1 probe (Addgene, Catalog nr. 36321)[75] and ZnT9-50Met, ZnT9-50Val, or empty pCDNA3 vectors in 6-well plates with 22 mm coverslips. 24 hours after transfection, cells were incubated with basal medium or supplemented with 100 µM ZnSO_4_ for 3 hours. Then, samples were placed under the microscope in ISO for imaging with an SP8 Leica microscope (Wetzlar, Germany). When blocking ZIP7 activity, 1 µM of the ZIP7 inhibitor NVS-ZP7-4; HY-114395, MedChemExpress (Quimigen) or 1 µM of DMSO used as a control were added at 1:1000 in the treatment 3 hours before imaging. The measured zinc content is dependent on the FRET signal, which is expressed as the mean of YFP/CFP ratios from each condition normalized across experiments to an empty vector in basal zinc conditions. Images were analyzed using ImageJ software.

Mitochondria zinc concentration was measured *in vivo* using a plate reader (VICTOR Nivo, Perkin Elmer). Cells were co-transfected with Mito-cCherry-Gn2Zn or SMAC-Gn2Zn probes [54] and ZnT9-50Met, ZnT9-50Val, or empty pCDNA3 vectors in 24-well plates. 24 hours after transfection, cells were incubated for 40 min with basal medium or supplemented with 100 µM ZnSO_4_. Then the media was changed to ISO solution and samples were placed in the plate reader for measuring. When blocking the ZIP7 activity, 1µM of the ZIP7 inhibitor NVS-ZP7-4 or DMSO was used as a control. Zinc content is expressed as the mean of GFP/Cherry ratios from each condition normalized to an empty vector in basal zinc conditions.

### MTT assays

Cells were transfected with ZnT9-50Met, ZnT9-50Val, or empty pCDNA3 vectors in 24-well plates and incubated for 24h with normal medium, supplemented with 100 µM ZnSO4 or Zn^2+^-free growth medium generated with Chelex 100 resin. Then, 3-(4,5-dimethylthiazol-2-yl)-2,5-diphenyltetrazolium bromide (MTT) reagent was added (0.5 mg/mL) for 2 h at 37 °C. After that, the supernatant was removed, and cells were resuspended in 400 µl of DMSO. The absorbance was read at 570 nm.

## Data and material availability

All genotyping and experimental data generated in the present study is provided as Supporting Information. Additionally, data from the following public resources was used: PopHuman genome browser, http://pophuman.uab.cat; Ensembl genome browser, http://grch37.ensembl.org/; UCSC Genome Browser, https://genome.ucsc.edu/; Denisova, AltaiNeandertal, Vindija33.19, and Chagyrskaya-Phalanx archaic VCF files, https://cdna.eva.mpg.de/neanderthal/; ARGweaver UCSC Genome browser track, http://compgen.cshl.edu/ARGweaver/introgressionHub/hub.txt; recombination maps, https://github.com/eyherabh/genetic_map_comparisons; Geography of Genetic Variants Browser, https://popgen.uchicago.edu/ggv/; ALFRED database, https://alfred.med.yale.edu/alfred/index.asp; GTEX portal, https://www.gtexportal.org/home/; GWAS catalog, https://www.ebi.ac.uk/gwas/; GWAS atlas, https://atlas.ctglab.nl/. Cell lines, probes and clones used in the reported experiments will be available upon request.

## Acknowledgments

We thank Martin Kuhlwilm for his comments and help. This work was supported by Ministerio de Ciencia e Innovación (MCIN) and Agencia Estatal de Investigación (AEI; DOI:10.13039/501100011033) with project grants PID2019-110933GB-I00 (to EB), PID2019-106755RB-I00 (to RV) and Unidad de Excelencia María de Maeztu CEX2018-000792-M (to EB and RV); and by Direcció General de Recerca, Generalitat de Catalunya with project grant 2017SGR00702 (to EB). JGC was supported with an FPI-MCIN/AEI PhD contract (PRE2020-095762).

## Author contributions

Conceptualization: RV, EB; Methodology: FC, GM, RV, EB; Formal analysis and investigation: ARU, MVG, AF, JGC, EG, AB, GIR, VHF, FC, GM, RV, EB; Supervision: RV, EB; Writing original draft: ARU and EB; Writing review & editing: GM, RV, EB; Funding acquisition: RV, EB.

## Supporting Information

**Figure S1.**
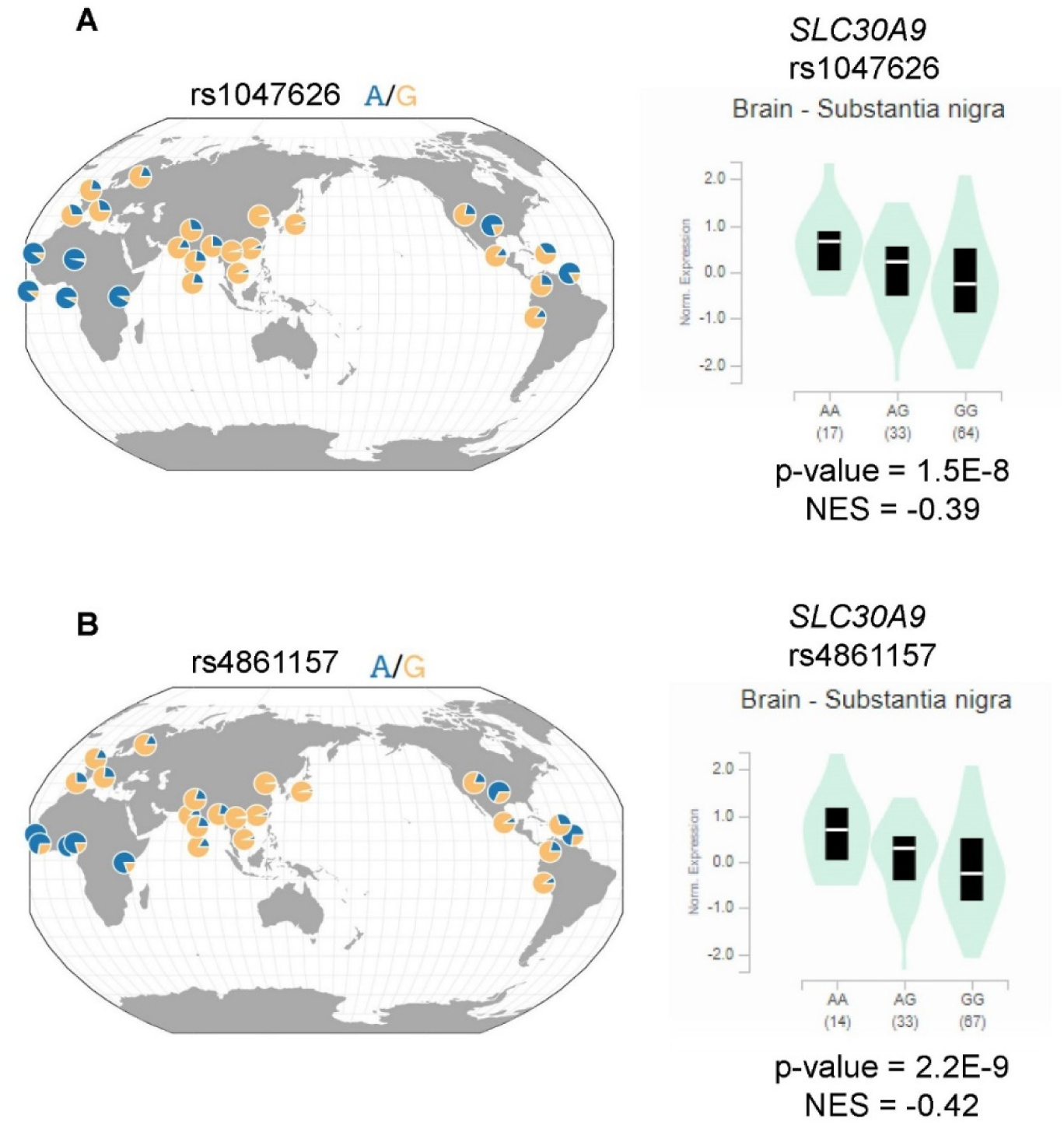
Worldwide allele frequencies and differential gene expression between genotypes at rs1047626 and rs4861157. (A) Allele frequencies at rs1047626 and rs4861157 across human populations in the 1000 Genomes Project [1]. Frequency plots were downloaded from the Geography of Genetic Variants Browser (https://popgen.uchicago.edu/ggv/) [2] (B) Differential *SLC30A9* expression in the substantia nigra according to the rs1047626 and rs4861157 genotypes as available at the GTEX portal (https://www.gtexportal.org/home/). NES, normalized effect sizes.

**Figure S2.**
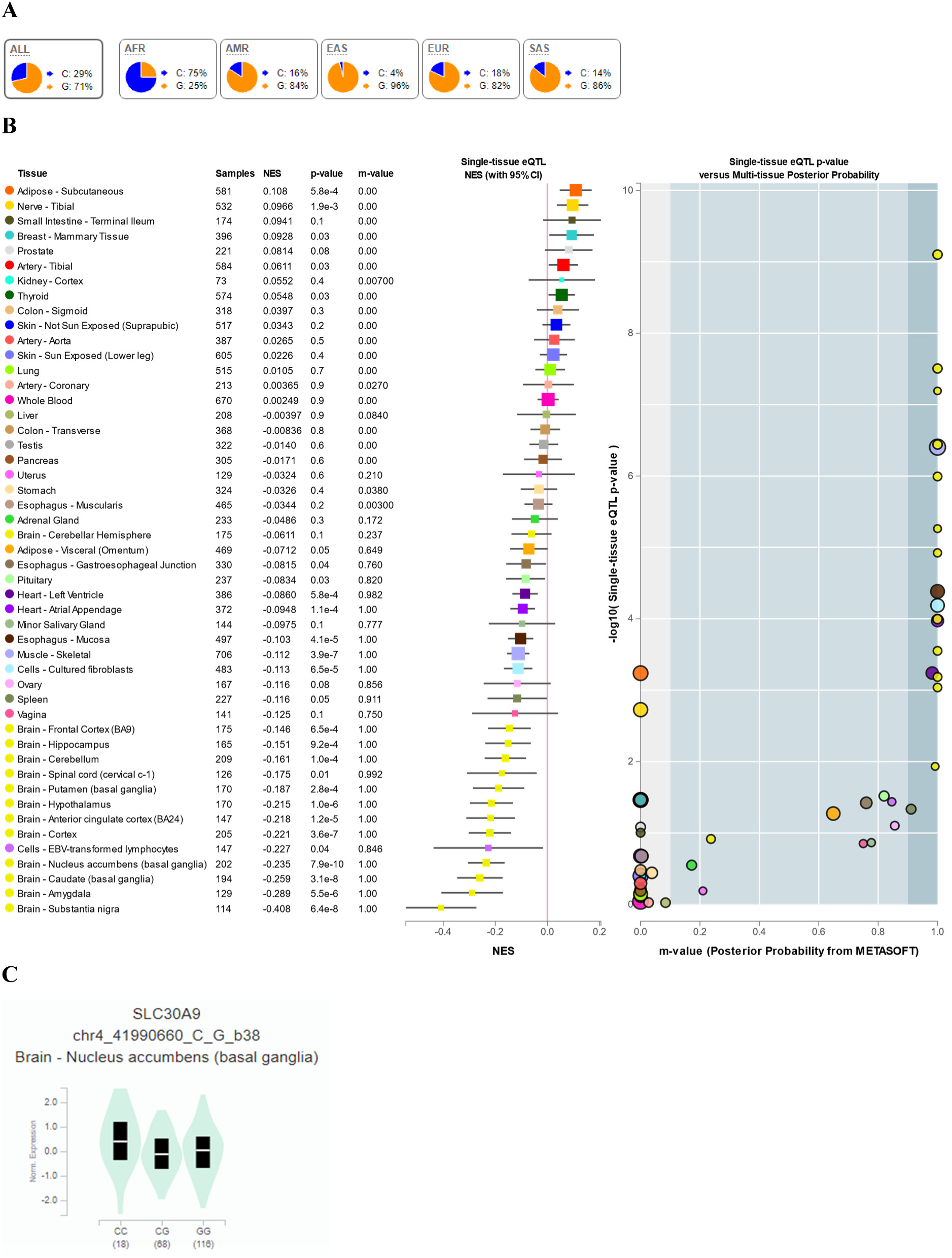
Continental allele frequencies and GTEX data for rs2581434. (A) Continental 1000 Genomes Project Phase 3 allele frequencies as retrieved from Ensembl (https://www.ensembl.org/index.html). (B) Multi-tissue eQTL comparison for rs2581434. (C) Differential *SLC30A9* expression in the substantia nigra according to the rs2581434 genotypes as available at the GTEX portal (https://www.gtexportal.org/home/). NES, normalized effect sizes.

**Figure S3.**
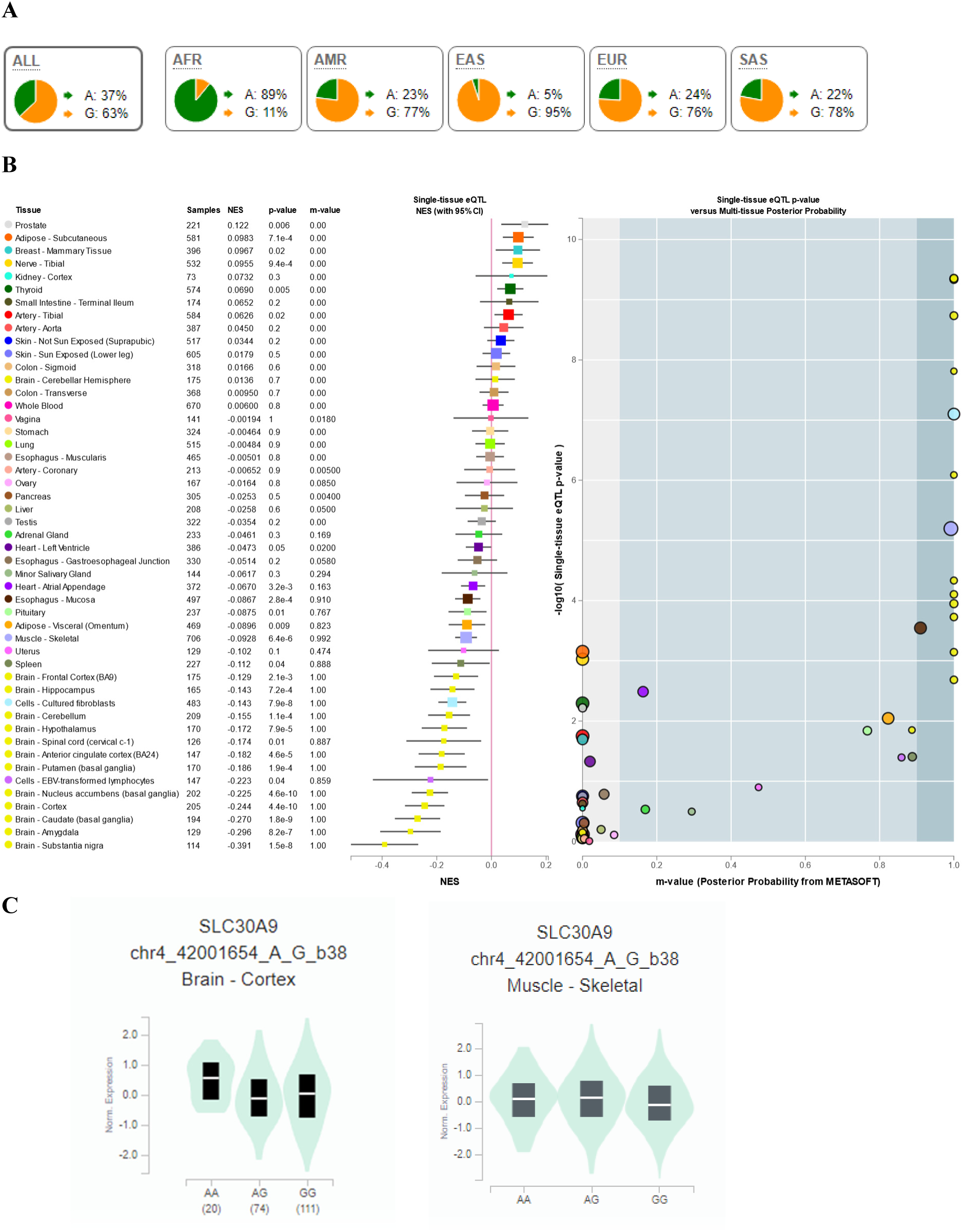
Continental allele frequencies and GTEX data for rs1047626. (A) Continental 1000 Genomes Project Phase 3 allele frequencies as retrieved from Ensembl (https://www.ensembl.org/index.html). (B) Multi-tissue eQTL comparison for rs1047626. (C) Differential *SLC30A9* expression in brain cortex and skeletal muscle according to the rs1047626 genotypes as available at the GTEX portal (https://www.gtexportal.org/home/). NES, normalized effect sizes.

**Figure S4.**
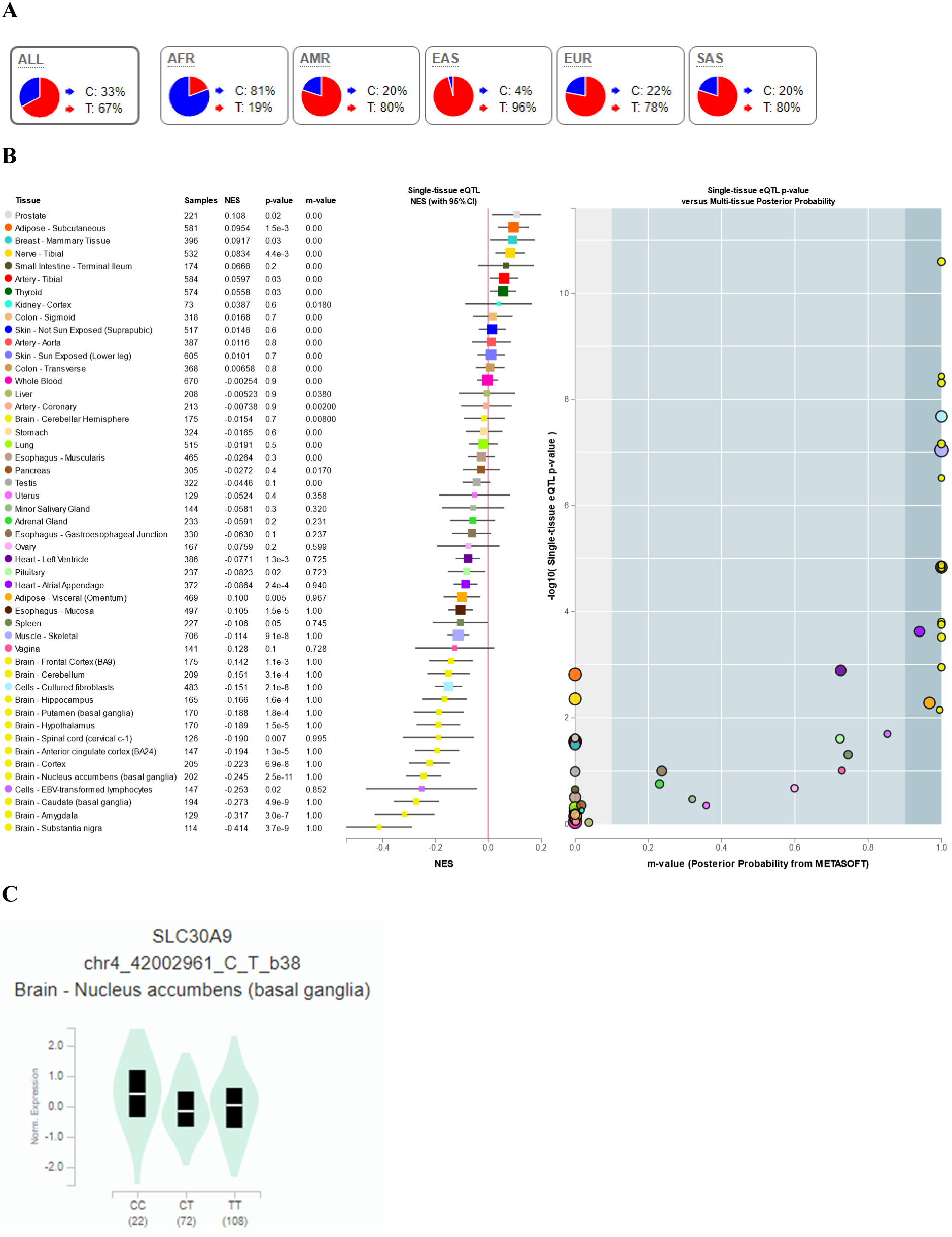
Continental allele frequencies and GTEX data for rs2581452. (A) Continental 1000 Genomes Project Phase 3 allele frequencies as retrieved from Ensembl (https://www.ensembl.org/index.html). (B) Multi-tissue eQTL comparison for rs2581452. (C) Differential *SLC30A9* expression in the nucleus accumbens according to the rs2581452 genotypes as available at the GTEX portal (https://www.gtexportal.org/home/). NES, normalized effect sizes.

**Figure S5.**
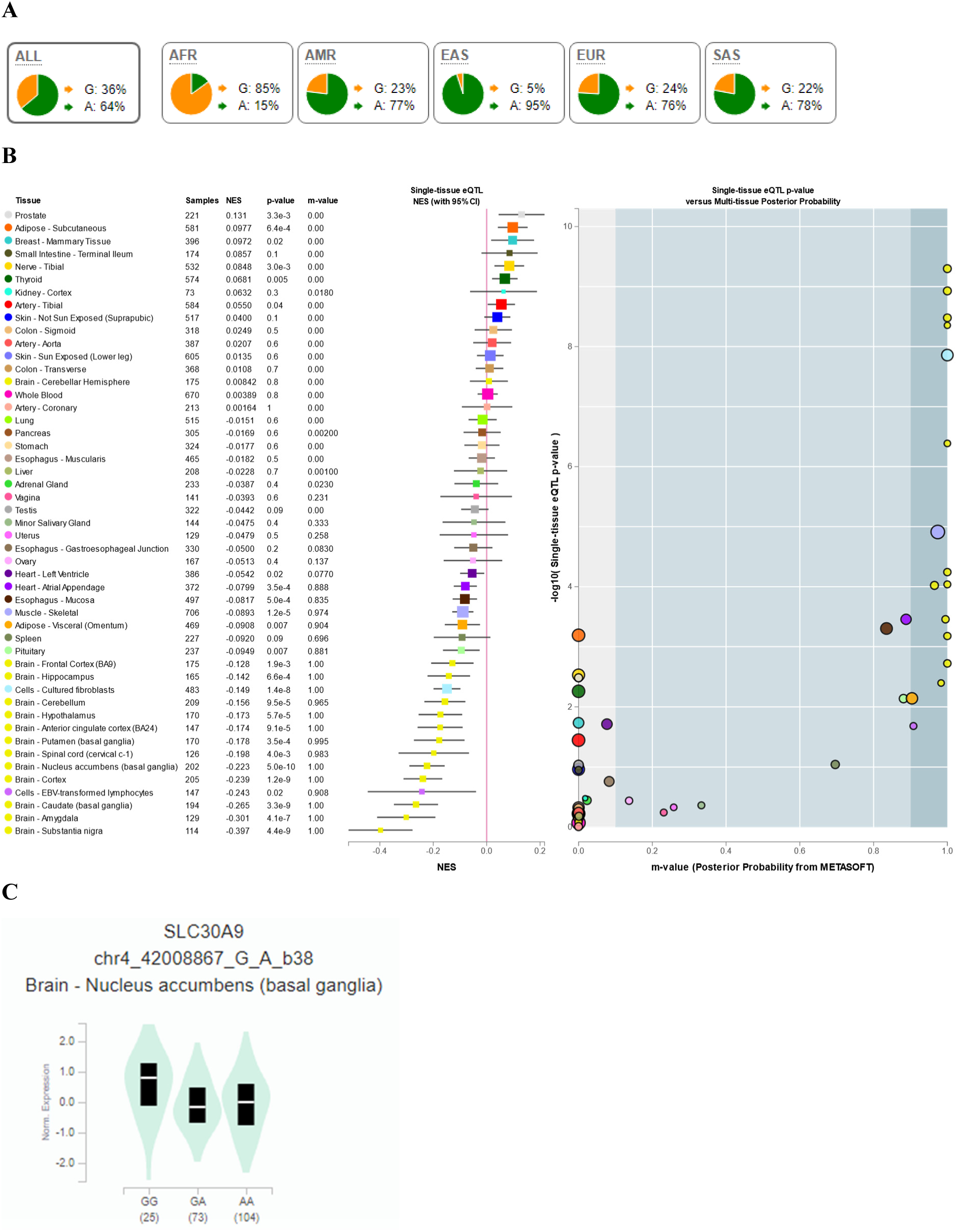
Continental allele frequencies and GTEX data for rs2660319. (A) Continental 1000 Genomes Project Phase 3 allele frequencies as retrieved from Ensembl (https://www.ensembl.org/index.html). (B) Multi-tissue eQTL comparison for rs2660319. (C) Differential *SLC30A9* expression in the nucleus accumbens according to the rs2660319 genotypes as available at the GTEX portal (https://www.gtexportal.org/home/). NES, normalized effect sizes.

**Figure S6.**
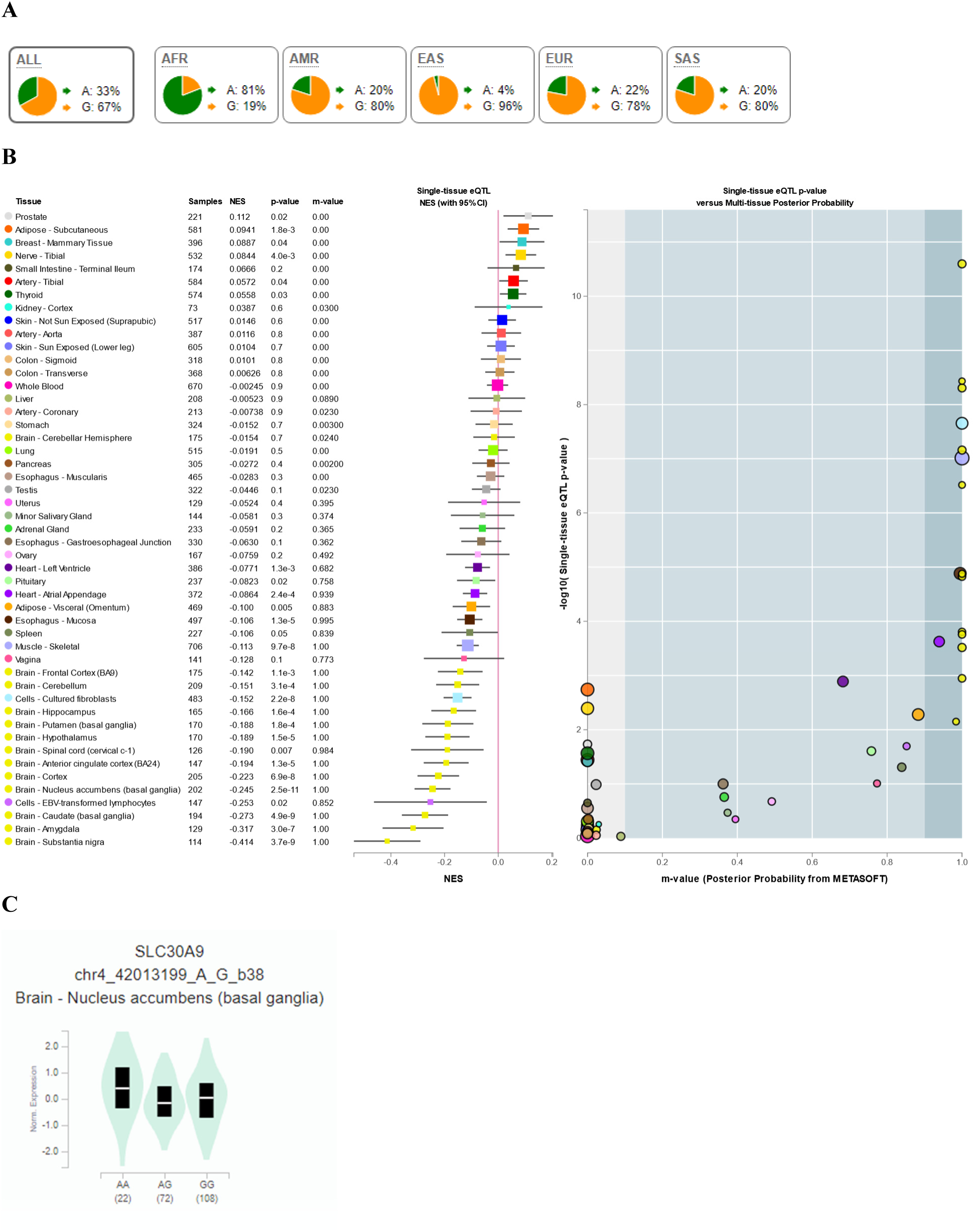
Continental allele frequencies and GTEX data for rs1848182. (A) Continental 1000 Genomes Project Phase 3 allele frequencies as retrieved from Ensembl (https://www.ensembl.org/index.html). (B) Multi-tissue eQTL comparison for rs1848182. (C) Differential *SLC30A9* expression in the nucleus accumbens according to the rs1848182 genotypes as available at the GTEX portal (https://www.gtexportal.org/home/). NES, normalized effect sizes.

**Figure S7.**
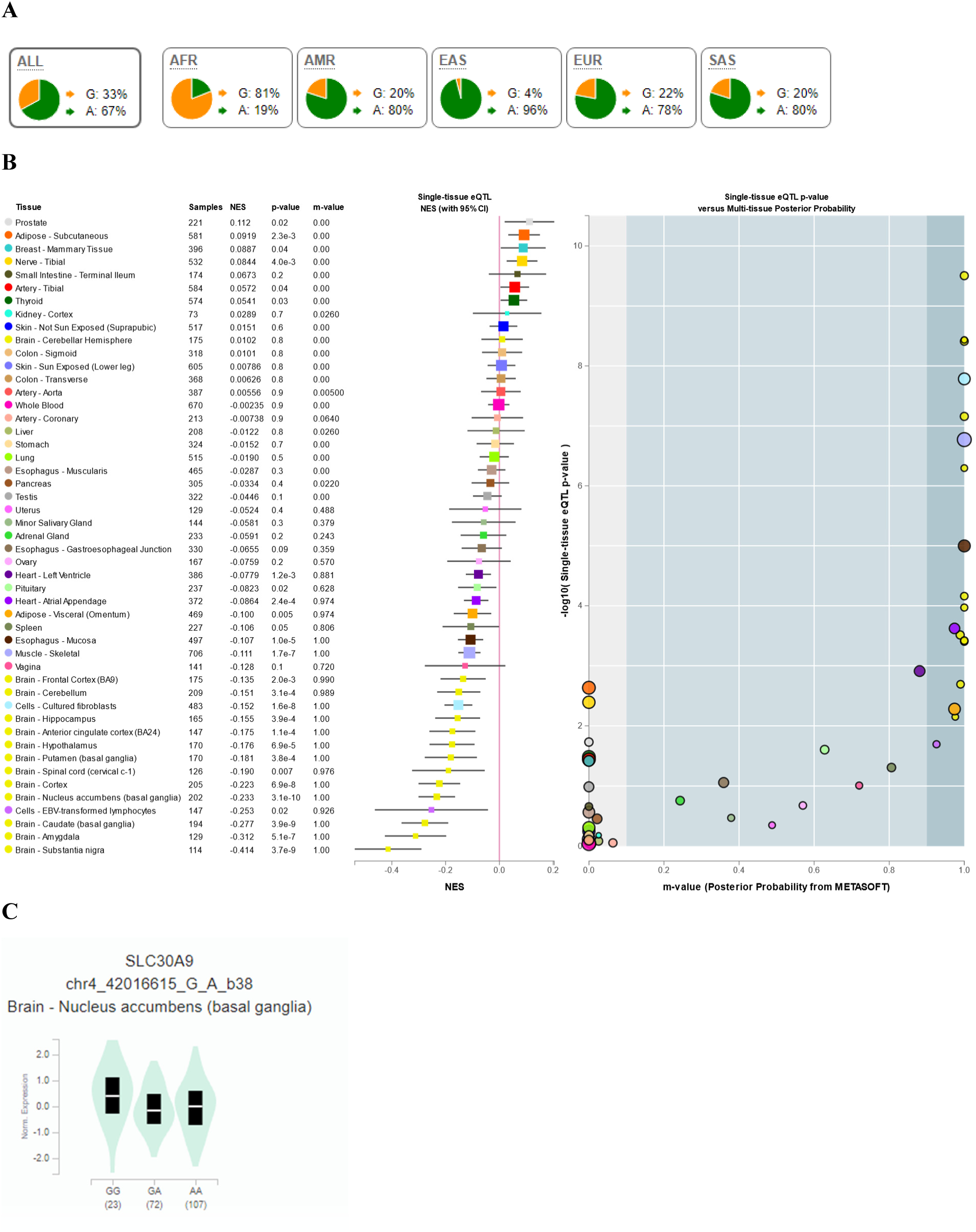
Continental allele frequencies and GTEX data for rs2581424. (A) Continental 1000 Genomes Project Phase 3 allele frequencies as retrieved from Ensembl (https://www.ensembl.org/index.html). (B) Multi-tissue eQTL comparison for rs2581424. (C) Differential *SLC30A9* expression in the nucleus accumbens according to the rs2581424 genotypes as available at the GTEX portal (https://www.gtexportal.org/home/). NES, normalized effect sizes.

**Figure S8.**
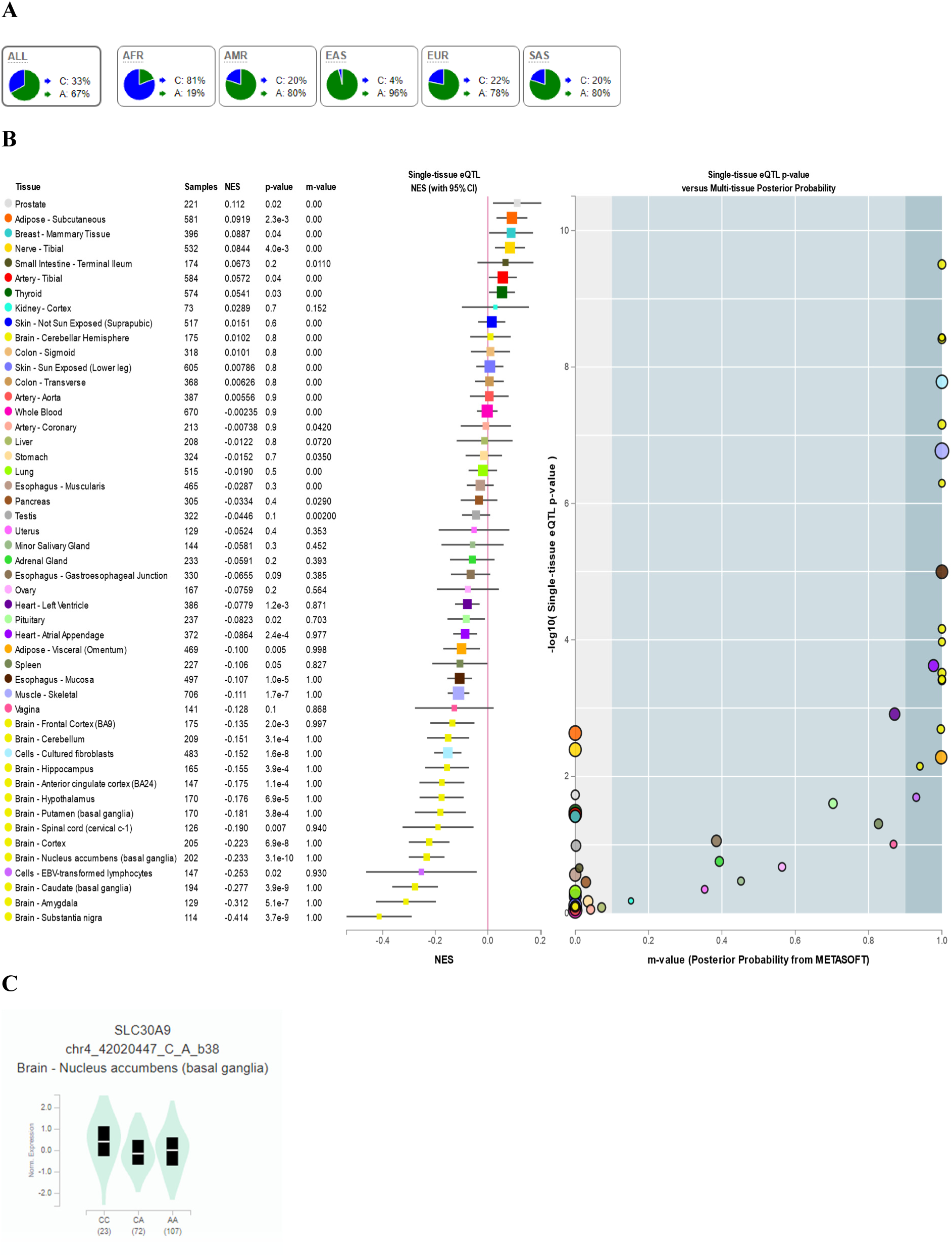
Continental allele frequencies and GTEX data for rs15857. (A) Continental 1000 Genomes Project Phase 3 allele frequencies as retrieved from Ensembl (https://www.ensembl.org/index.html). (B) Multi-tissue eQTL comparison for rs15857. (C) Differential *SLC30A9* expression in the nucleus accumbens according to the rs15857 genotypes as

**Figure S9.**
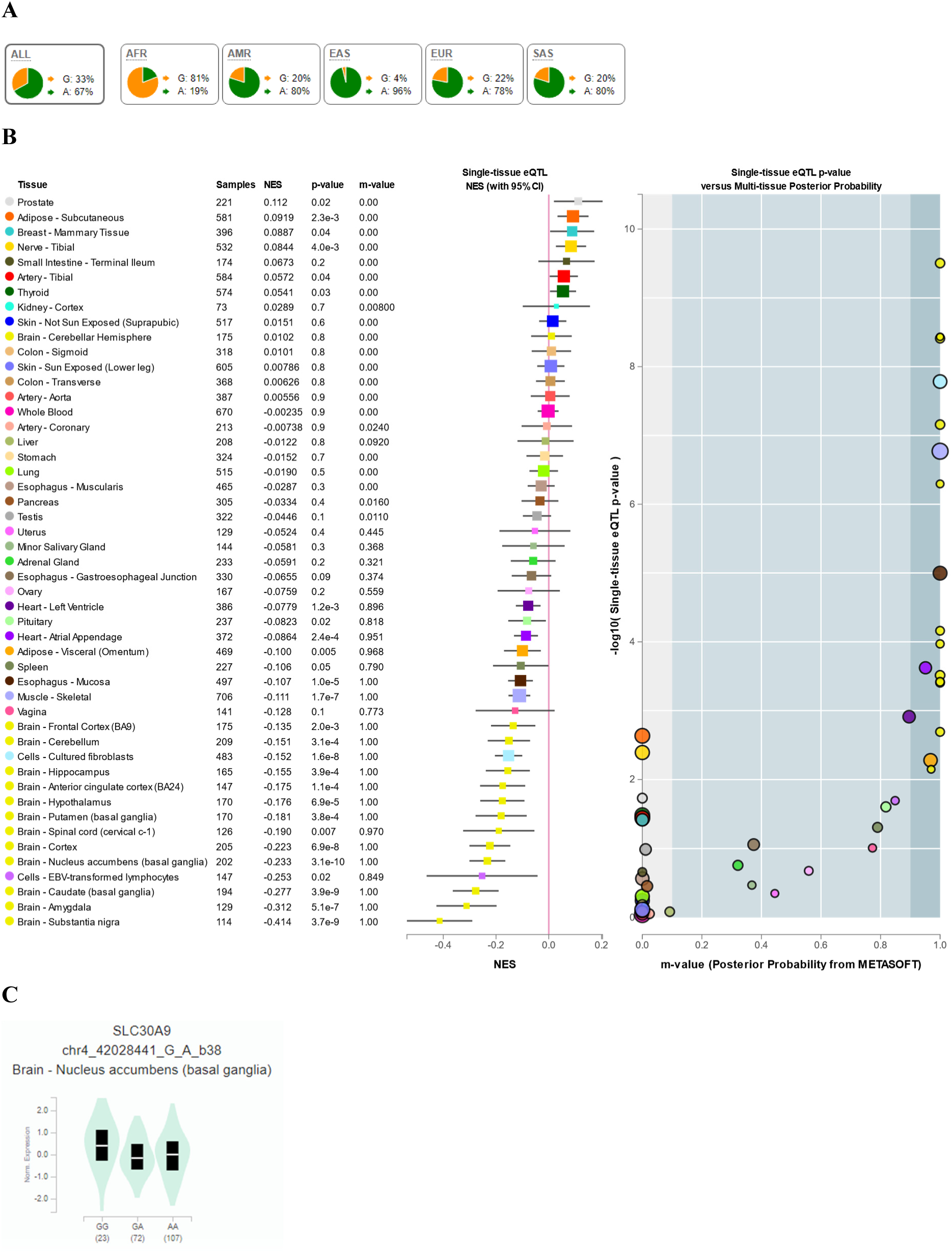
Continental allele frequencies and GTEX data for rs7439806. (A) Continental 1000 Genomes Project Phase 3 allele frequencies as retrieved from Ensembl (https://www.ensembl.org/index.html). (B) Multi-tissue eQTL comparison for rs7439806. (C) Differential *SLC30A9* expression in the nucleus accumbens according to the rs7439806 genotypes as

**Figure S10.**
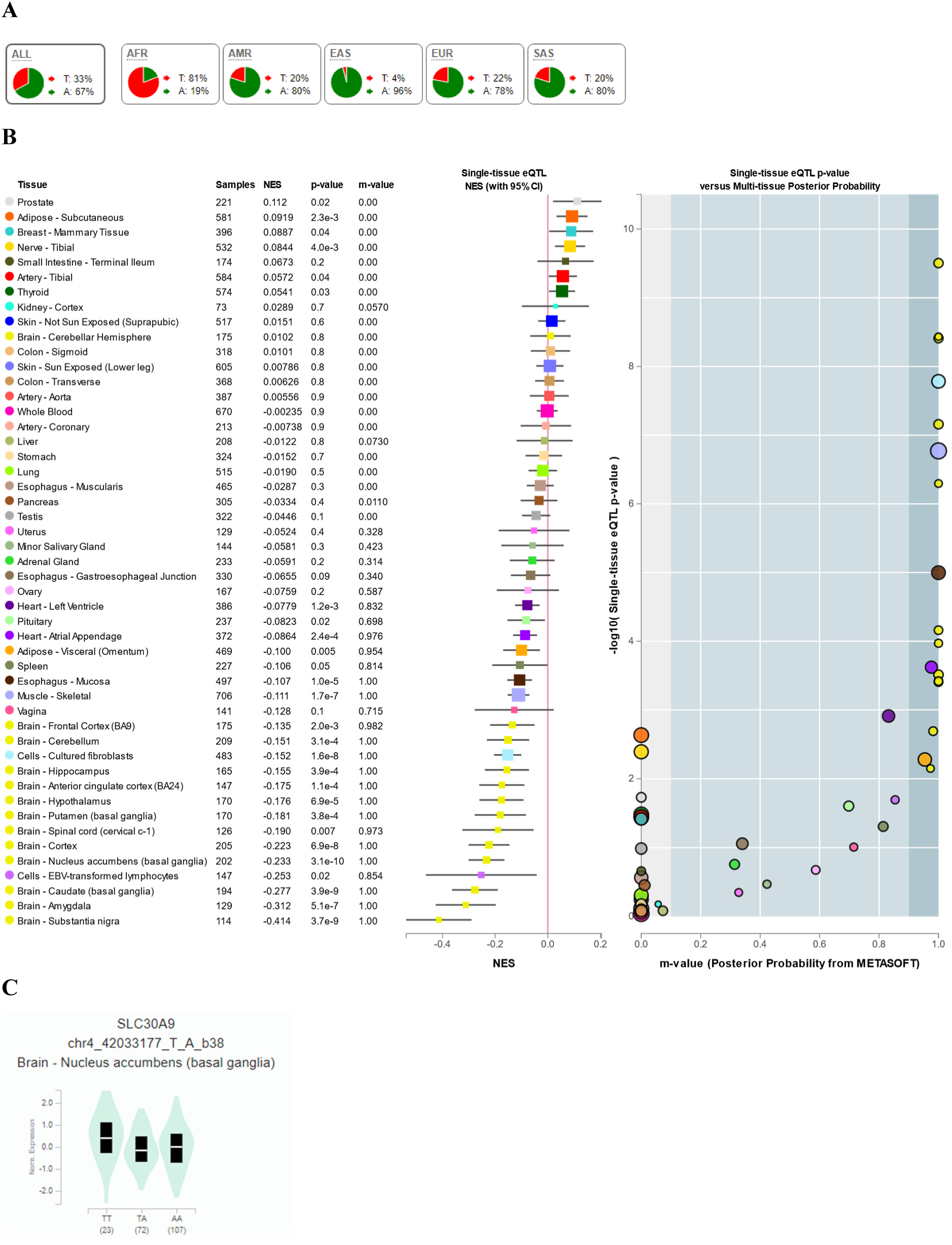
Continental allele frequencies and GTEX data for rs55835604. (A) Continental 1000 Genomes Project Phase 3 allele frequencies as retrieved from Ensembl (https://www.ensembl.org/index.html). (B) Multi-tissue eQTL comparison for rs55835604. (C) Differential *SLC30A9* expression in the nucleus accumbens according to the rs55835604 genotypes as

**Figure S11.**
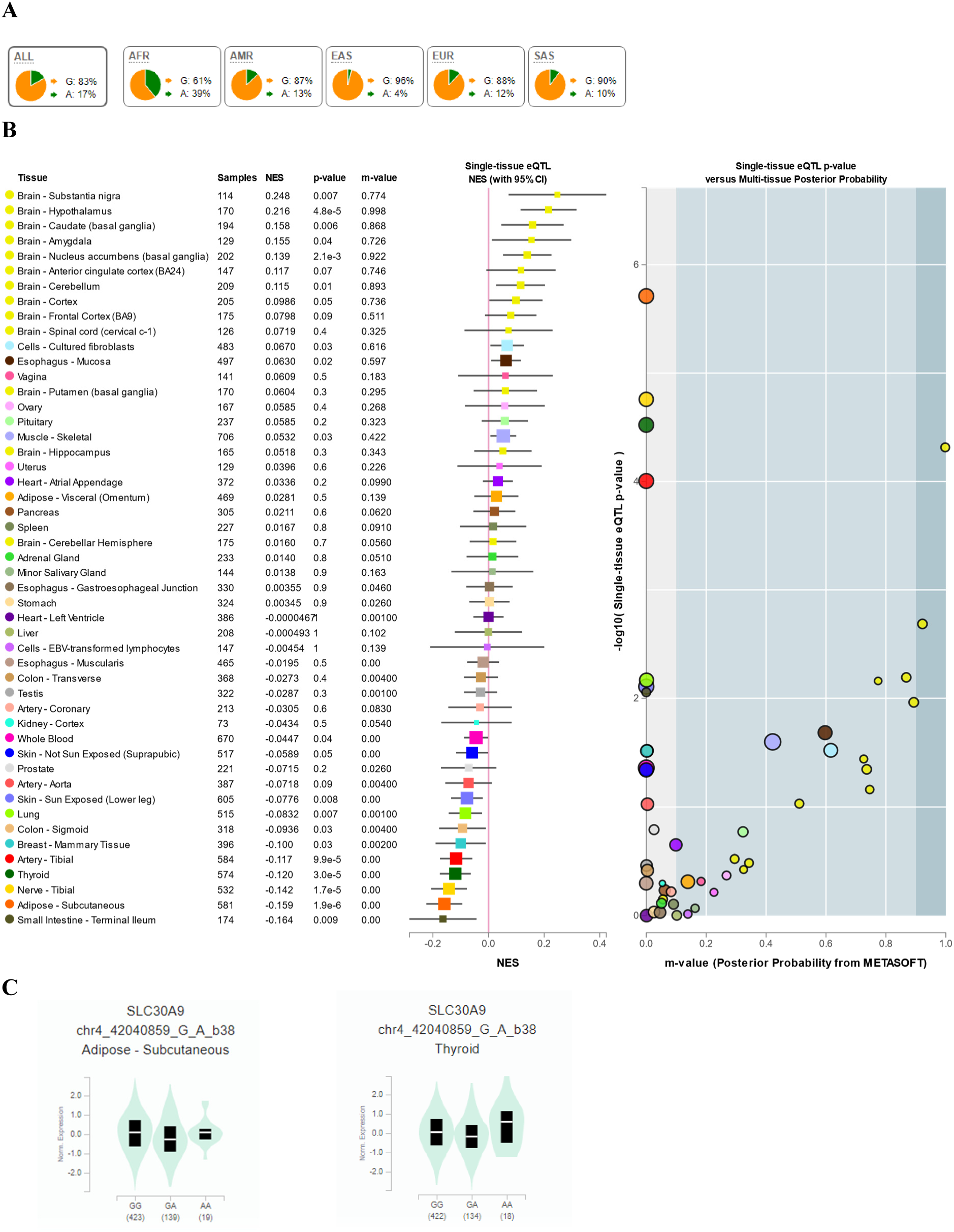
Continental allele frequencies and GTEX data for rs12510574. (A) Continental 1000 Genomes Project Phase 3 allele frequencies as retrieved from Ensembl. (B) Multi-tissue eQTL comparison for rs12510574. (C) Differential *SLC30A9* expression in the subcutaneous adipose tissue and the thyroid according to the rs12510574 genotypes as available at the GTEX portal (https://www.gtexportal.org/home/). NES, normalized effect sizes.

**Figure S12.**
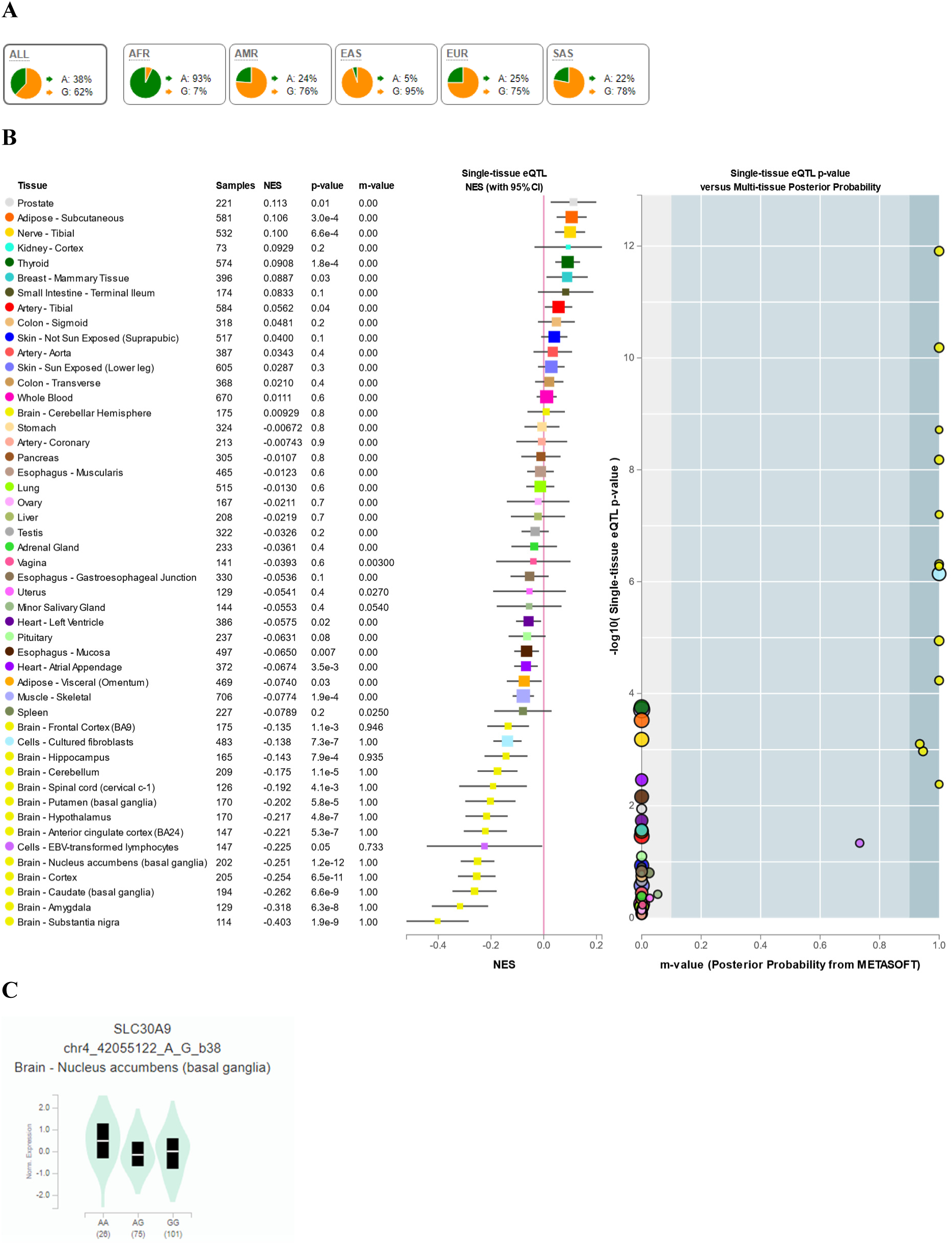
Continental allele frequencies and GTEX data for rs4861014. (A) Continental 1000 Genomes Project Phase 3 allele frequencies as retrieved from Ensembl (https://www.ensembl.org/index.html). (B) Multi-tissue eQTL comparison for rs4861014. (C) Differential *SLC30A9* expression in the nucleus accumbens according to the rs4861014 genotypes as available at the GTEX portal (https://www.gtexportal.org/home/). NES, normalized effect sizes.

**Figure S13.**
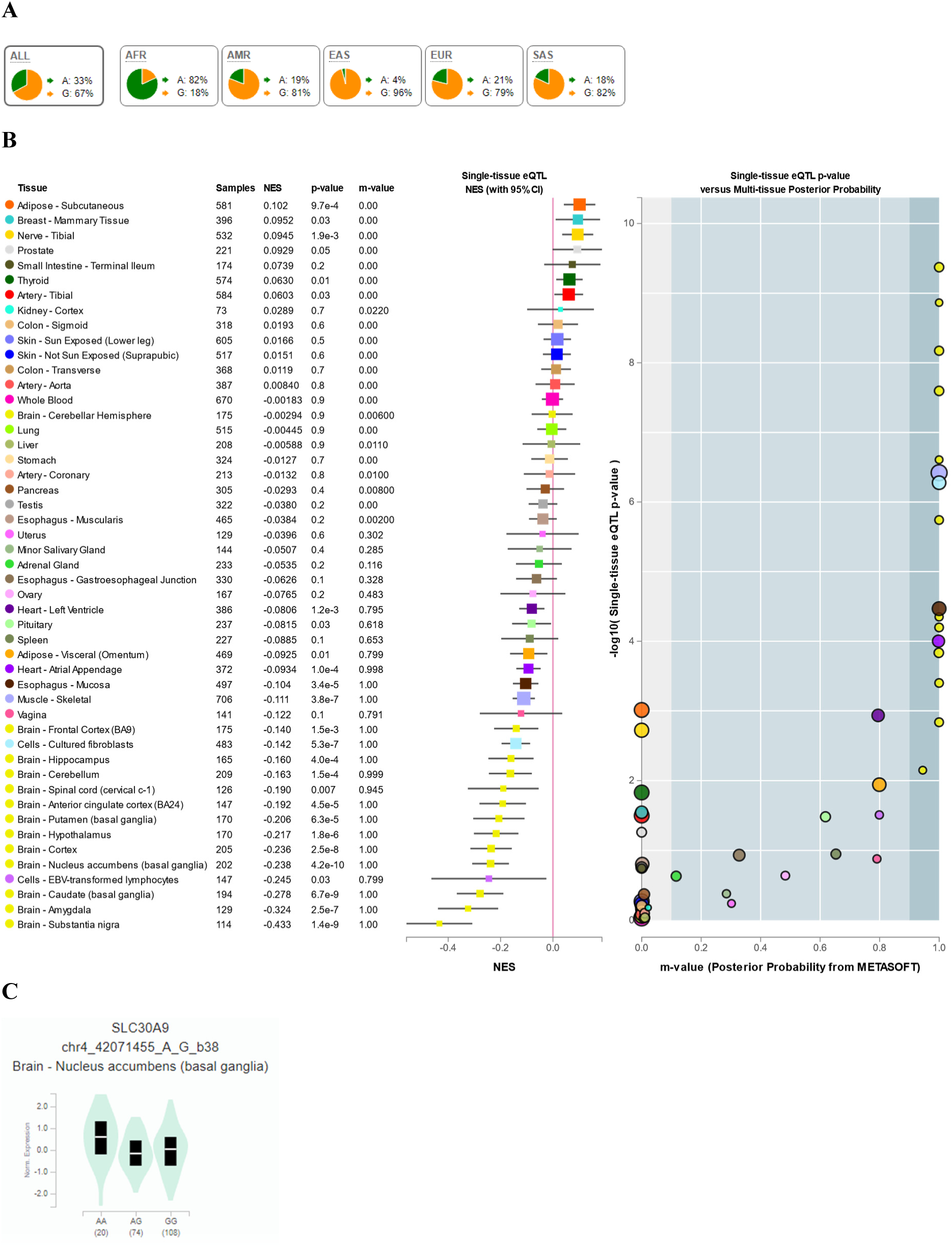
Continental allele frequencies and GTEX data for rs10019356. (A) Continental 1000 Genomes Project Phase 3 allele frequencies as retrieved from Ensembl (https://www.ensembl.org/index.html). (B) Multi-tissue eQTL comparison for rs10019356. (C) Differential *SLC30A9* expression in the nucleus accumbens according to the rs10019356 genotypes as available at the GTEX portal (https://www.gtexportal.org/home/). NES, normalized effect sizes.

**Figure S14.**
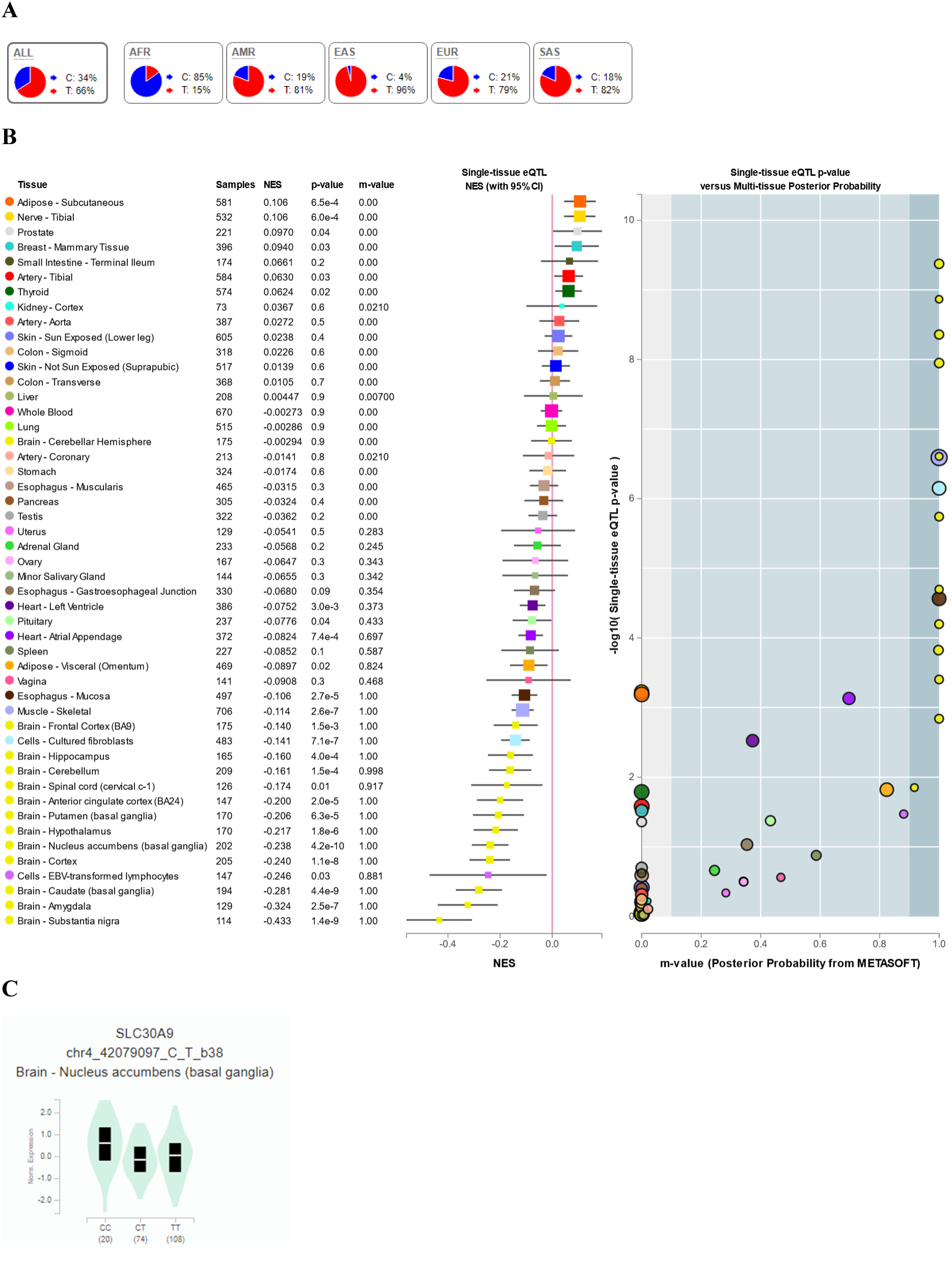
Continental allele frequencies and GTEX data for rs7660223. (A) Continental 1000 Genomes Project Phase 3 allele frequencies as retrieved from Ensembl (https://www.ensembl.org/index.html). (B) Multi-tissue eQTL comparison for rs7660223. (C) Differential *SLC30A9* expression in the nucleus accumbens according to the rs7660223 genotypes as

**Figure S15.**
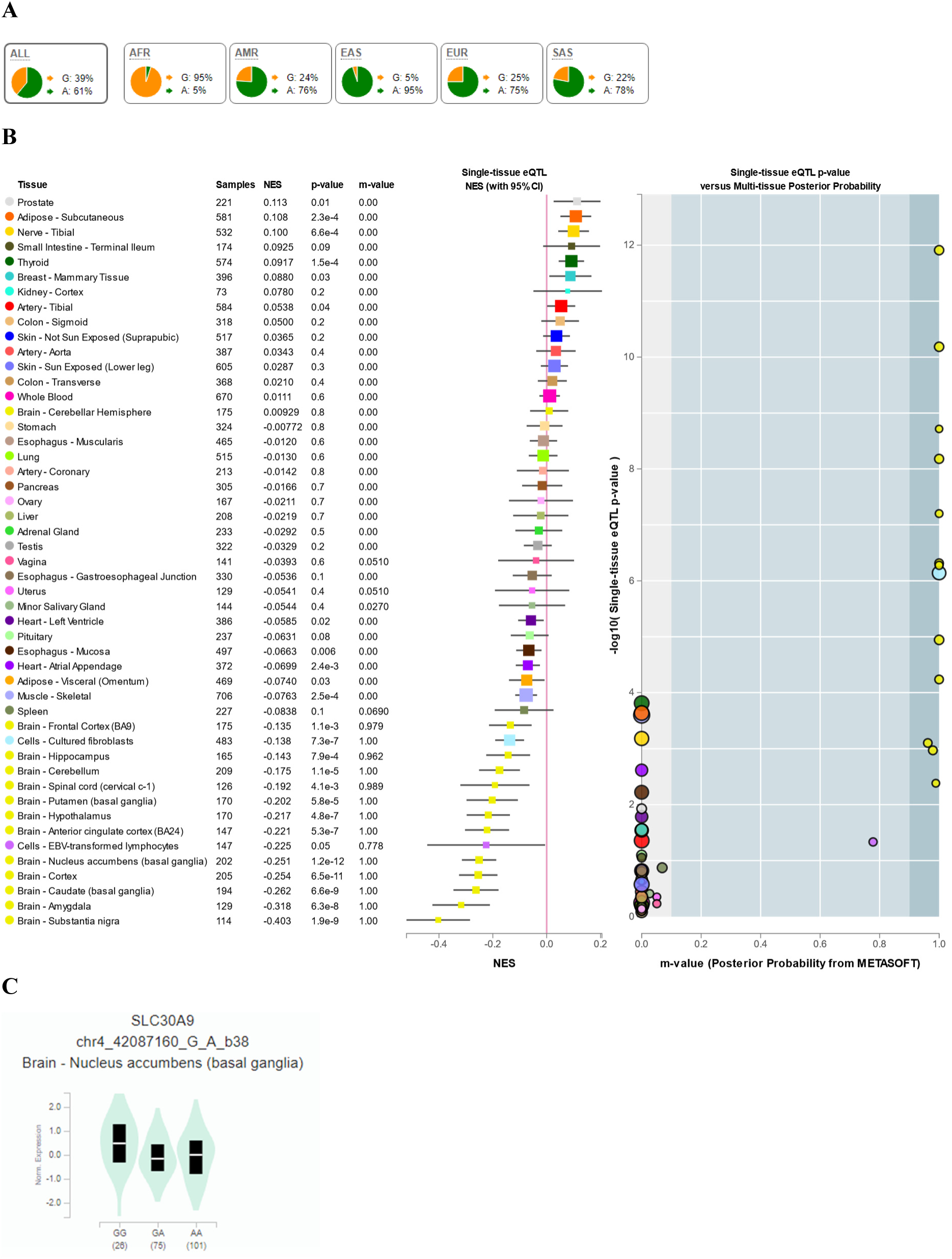
Continental allele frequencies and GTEX data for rs11051. (A) Continental 1000 Genomes Project Phase 3 allele frequencies as retrieved from Ensembl (https://www.ensembl.org/index.html). (B) Multi-tissue eQTL comparison for rs11051. (C) Differential *SLC30A9* expression in the nucleus accumbens according to the rs11051 genotypes as

**Figure S16.**
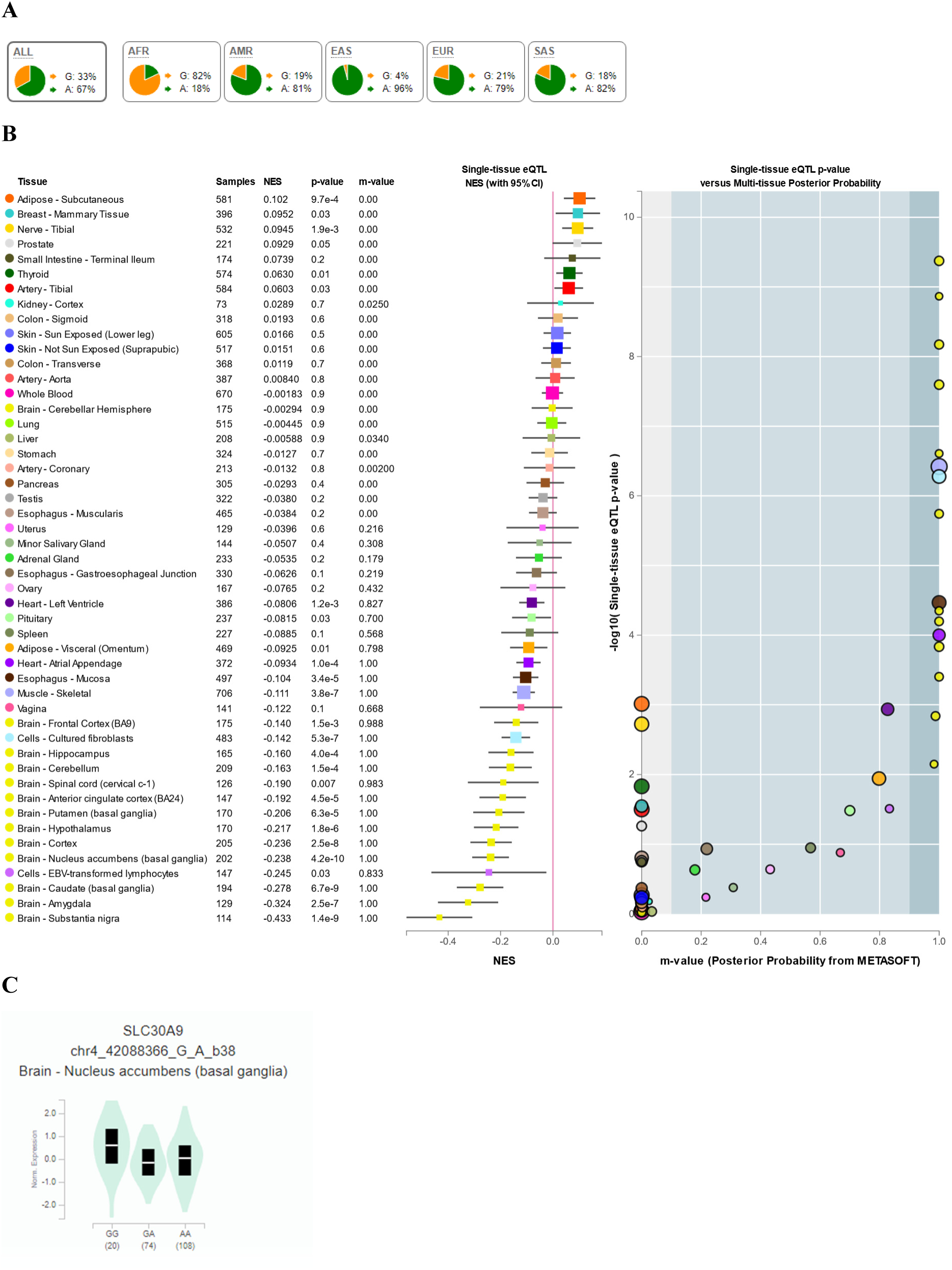
Continental allele frequencies and GTEX data for rs11935648. (A) Continental 1000 Genomes Project Phase 3 allele frequencies as retrieved from Ensembl (https://www.ensembl.org/index.html). (B) Multi-tissue eQTL comparison for rs11935648. (C) Differential *SLC30A9* expression in the nucleus accumbens according to the rs11935648 genotypes as

**Figure S17.**
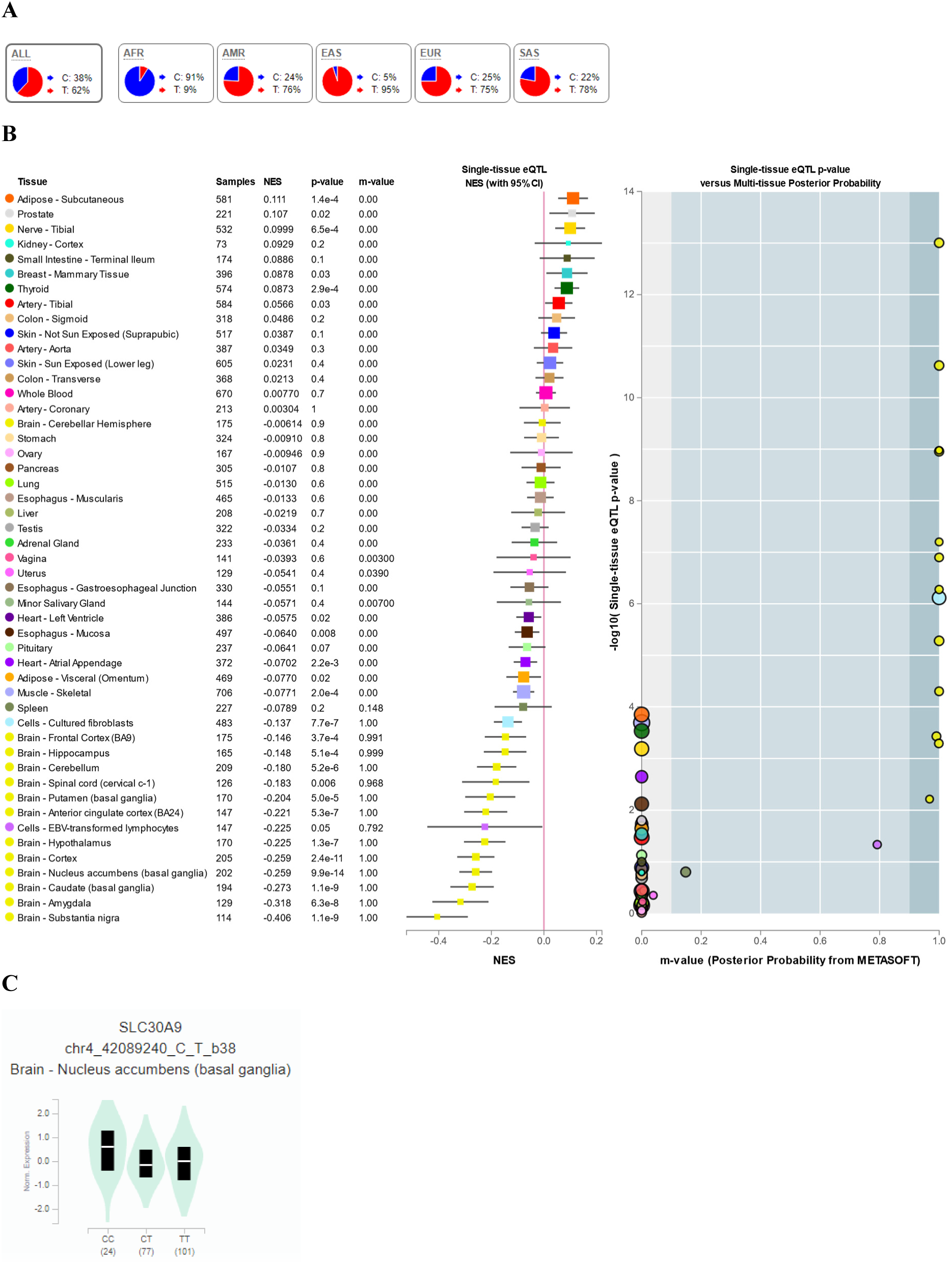
Continental allele frequencies and GTEX data for rs12511999. (A) Continental 1000 Genomes Project Phase 3 allele frequencies as retrieved from Ensembl (https://www.ensembl.org/index.html). (B) Multi-tissue eQTL comparison for rs12511999. (C) Differential *SLC30A9* expression in the nucleus accumbens according to the rs12511999 genotypes as

**Figure S18.**
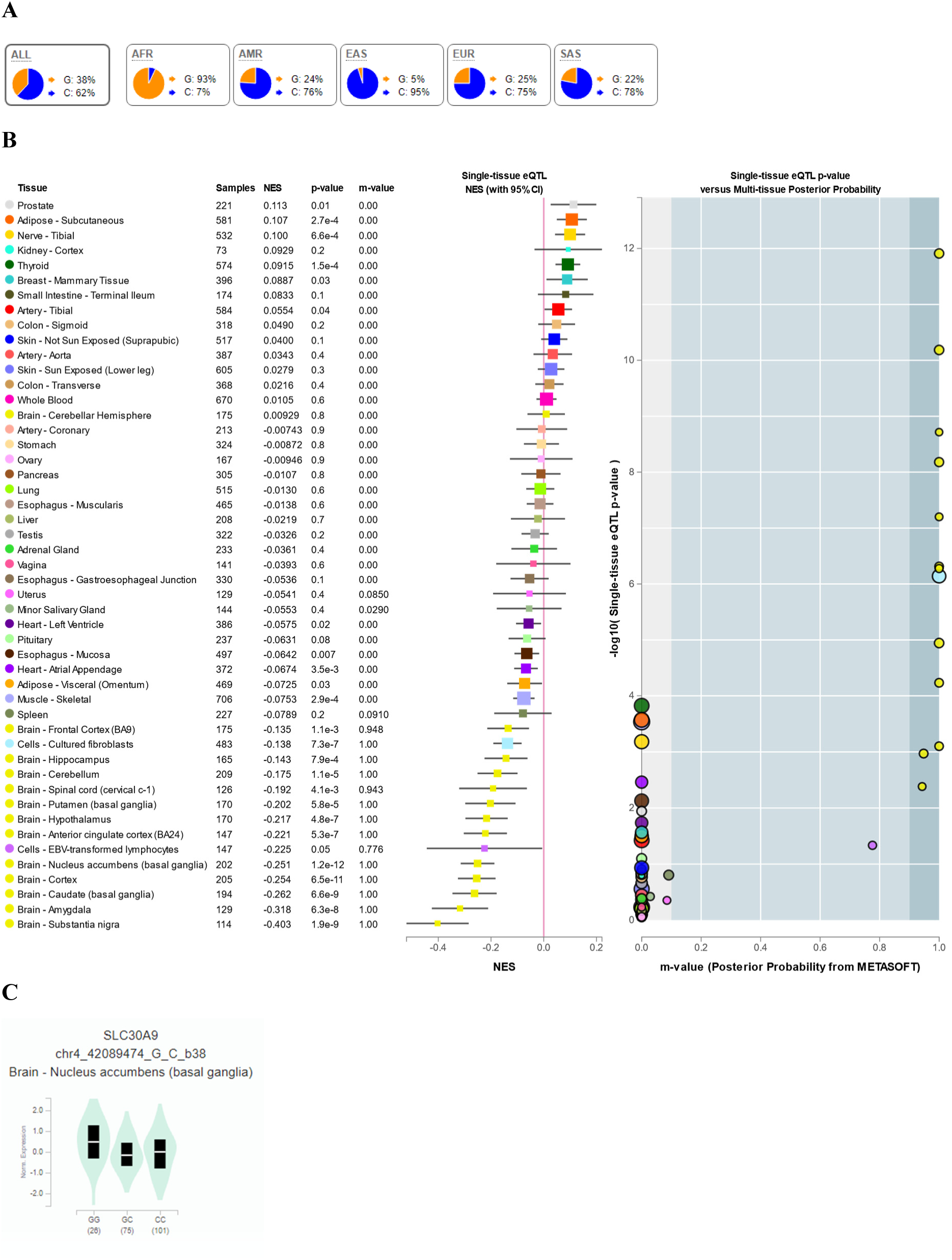
Continental allele frequencies and GTEX data for rs10938178. (A) Continental 1000 Genomes Project Phase 3 allele frequencies as retrieved from Ensembl (https://www.ensembl.org/index.html). (B) Multi-tissue eQTL comparison for rs10938178. (C) Differential *SLC30A9* expression in the nucleus accumbens according to the rs10938178 genotypes as

**Figure S19.**
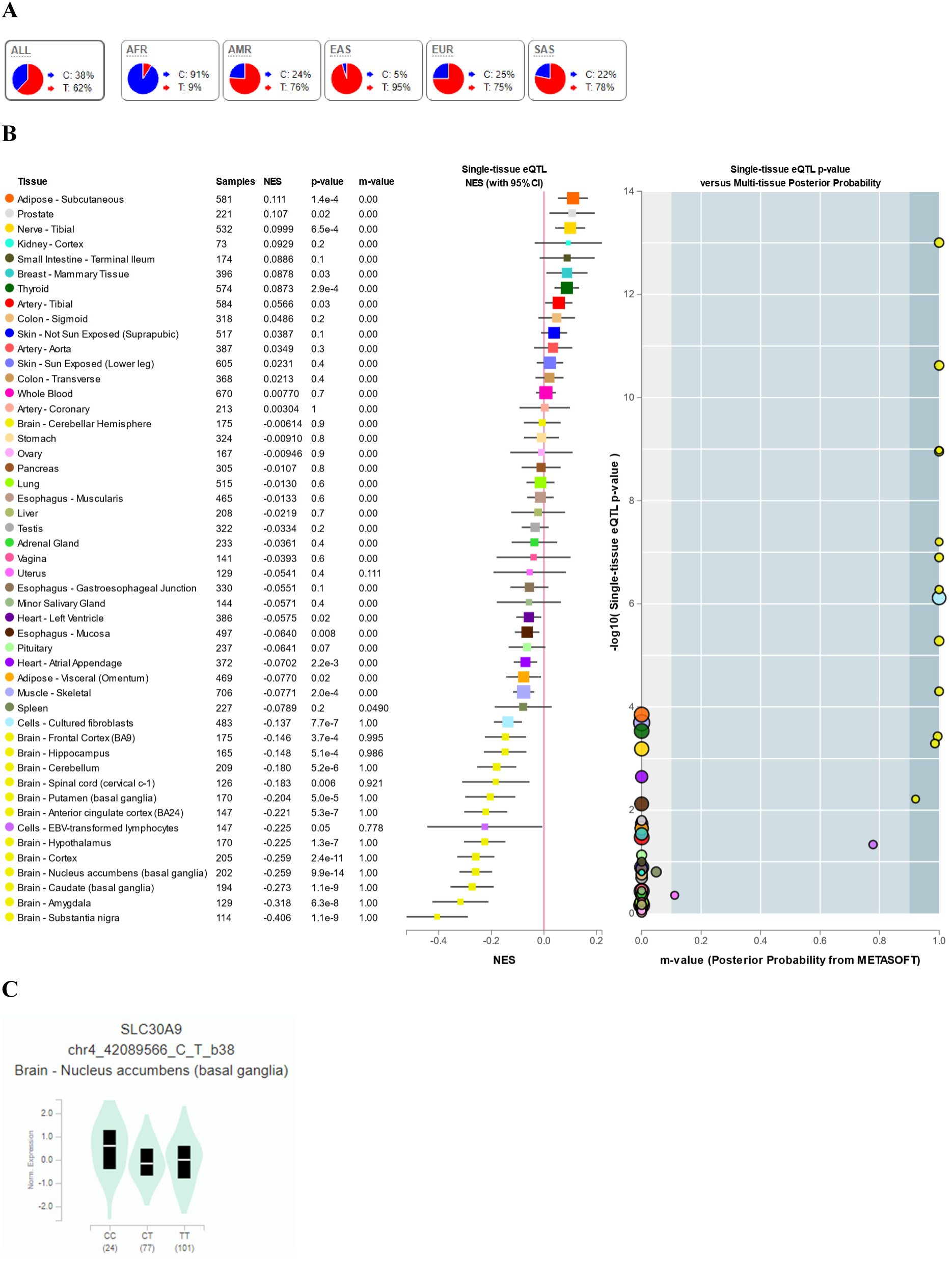
Continental allele frequencies and GTEX data for rs12512101. (A) Continental 1000 Genomes Project Phase 3 allele frequencies as retrieved from Ensembl (https://www.ensembl.org/index.html). (B) Multi-tissue eQTL comparison for rs12512101. (C) Differential *SLC30A9* expression in the nucleus accumbens according to the rs12512101 genotypes as

**Figure S20.**
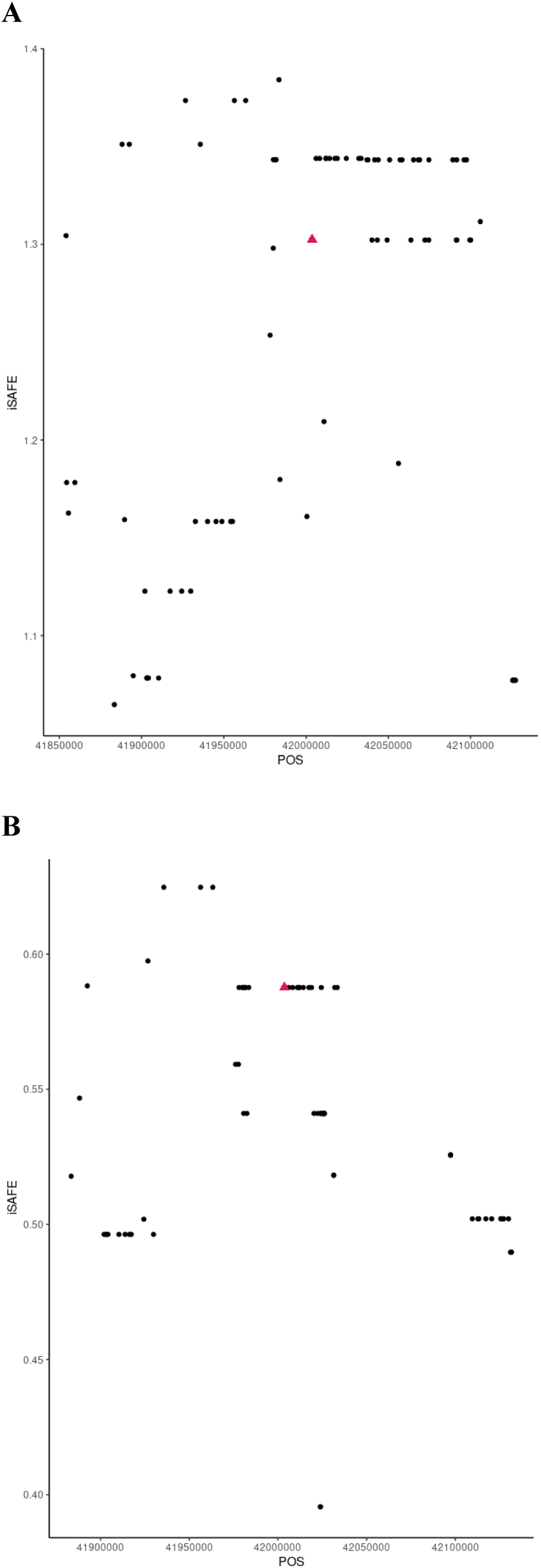
iSAFE values along the *SLC30A9* region. iSAFE values were computed within 300 kb (chr4: 41853671 – 42153671, GRCh37/hg19) centered on rs1047626, shown in red. A. iSAFE values in CHB. B. iSAFE values in CEU. Further details on the SNP annotation and iSAFE values are available in Tables S2-3.

**Figure S21.**
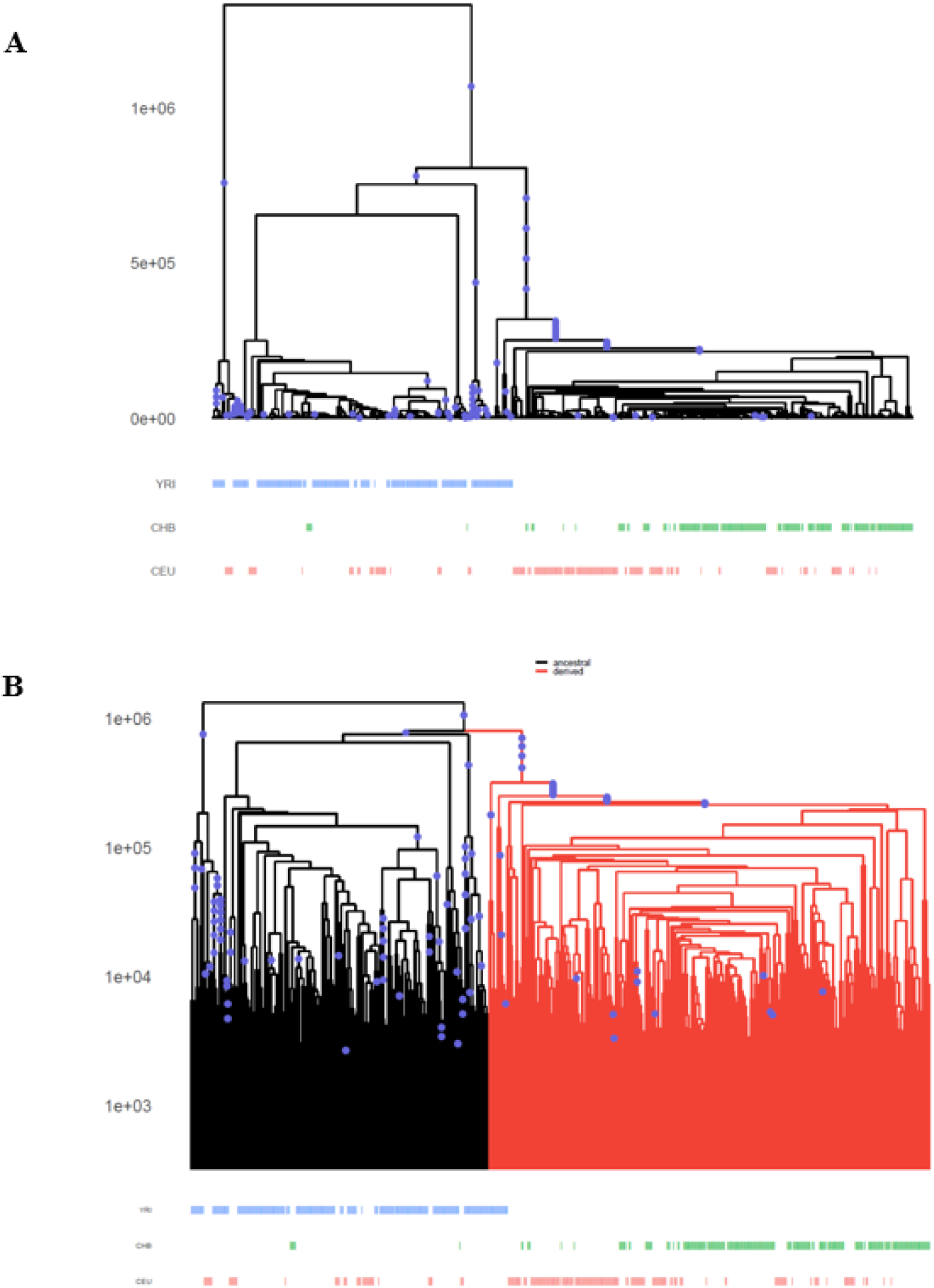
Positive selection acting on rs1047626. A) Marginal tree corresponding to the rs1047626 flanking region (chr4: 42002370 - 42004739; GRCh37/hg19) at *SLC30A9*. The derived allele at this SNP expanded rapidly in CHB and CEU, which is indicative of positive selection. B) Tree of interest highlighting those lineages carrying the derived allele at rs1047626.

**Figure S22.**
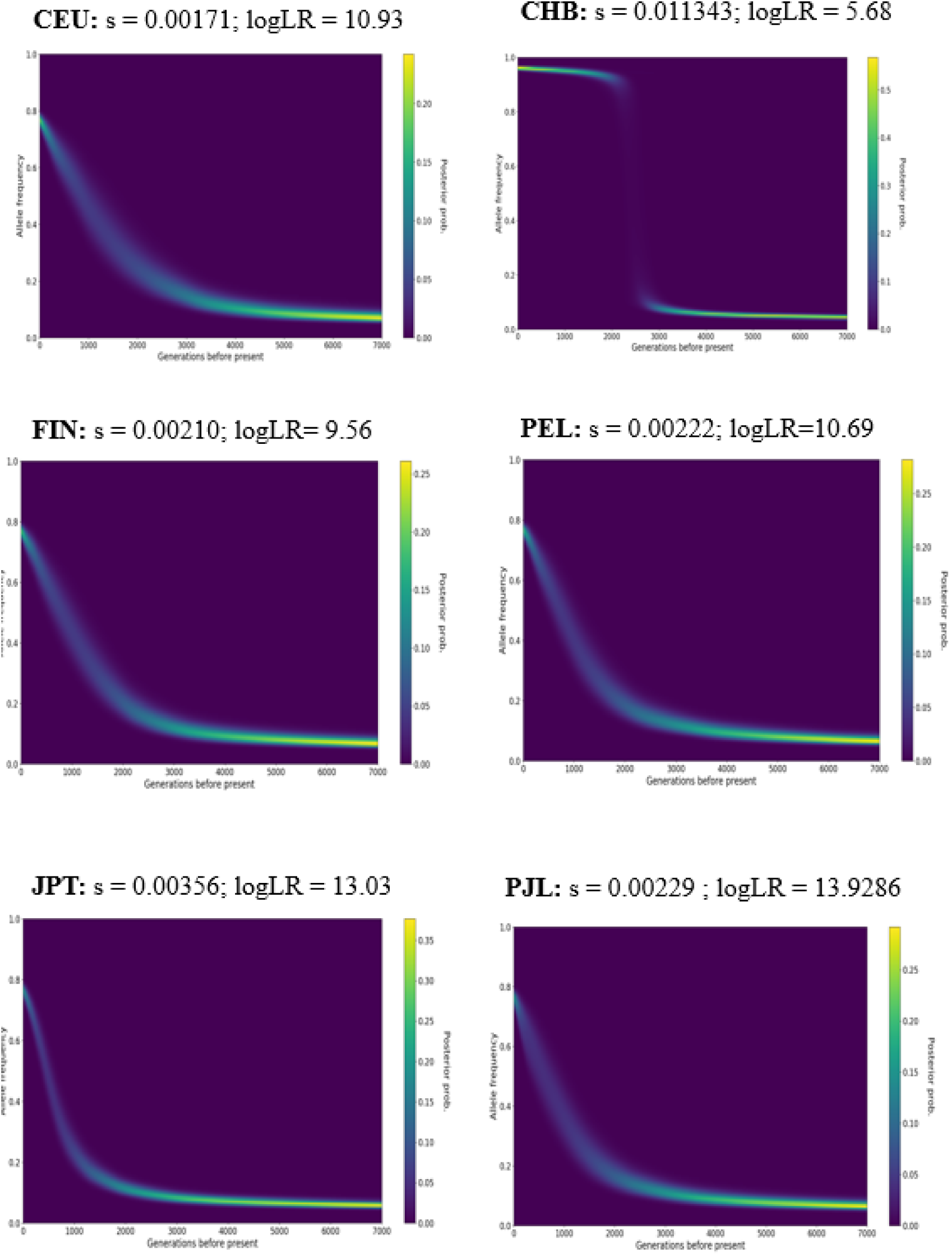
Allelic trajectories and selection coefficient for rs1047626 inferred by CLUES. CEU, Utah residents (CEPH) with Northern and Western European ancestry; CHB, Han Chinese in Beijing, China; FIN, Finnish in Finland; PEL, Peruvian in Lima, Peru; JPT, Japanese in Tokyo, Japan; PJL, Punjabi in Lahore, Pakistan.

**Figure S23.**
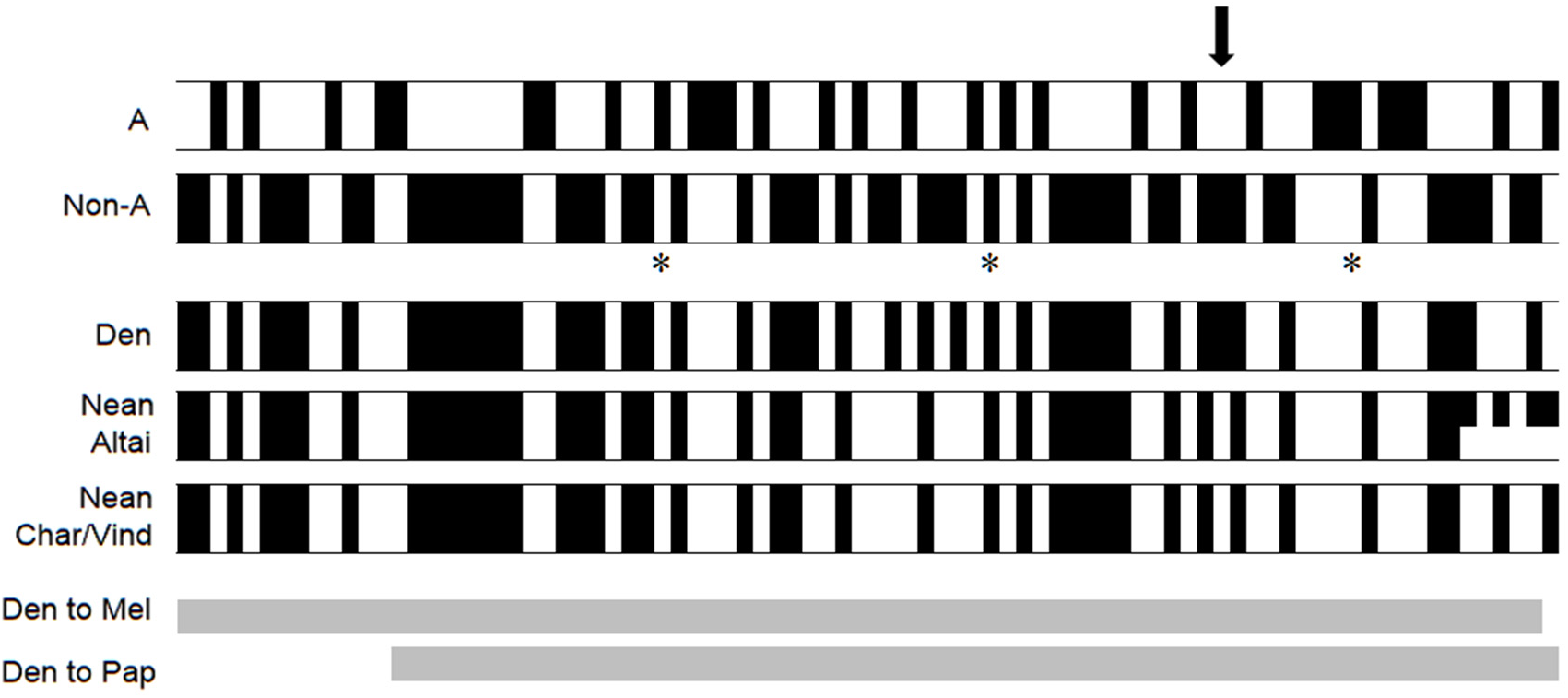
Haplotype structure of two major contrasting human haplotypes at *SCL30A9* and comparison with archaic humans. Schematic representation of the two major human haplotypes as defined by 84 SNPs in high linkage disequilibrium (r^2^>0.8) with rs1047626 in CEU and CHB along the putatively inferred introgressed region (chr4: 41,977,828-42,048,441; GRCh38) and the corresponding allele states found in four high coverage Denisovan and Neanderthal genomes. Derived states are indicated in black, and ancestral alleles in white. Den to Papuan and Den to Melanesian indicate Denisovan introgression segments previously described in Melanesians [3] and Papua New Guineans [4]. The black arrow point to the rs1047626 position. GWAS hits are indicated with an asterisk (see Table S4 for details).

**Figure S24.**
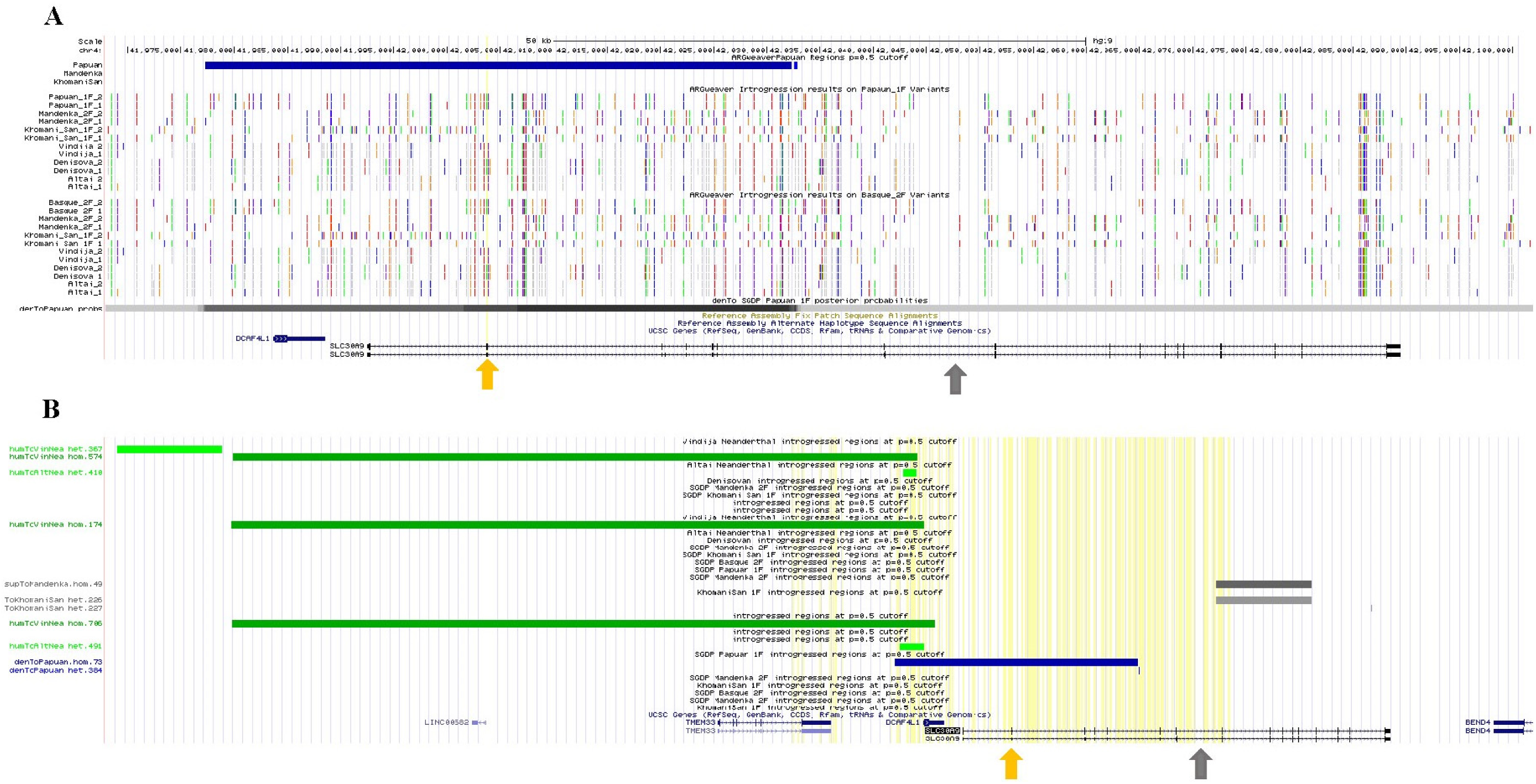
Introgression tracks around the *SLC30A9* region. A) ARGweaver Denisovan introgression results track in modern Papuans as available at the UCSC Genome Browser on Human GRCh37/hg19 and according to the methods and model described in Hubisz et al. (2020) [4]. Arrows in yellow and grey indicate the positions of rs1047626 and rs4861157, respectively. The “Region” track shows the predicted Denisovan introgressed regions in blue using a posterior probability cut-off of 0.5. For each individual, variants used in the analysis are shown above with alternating colours indicating variant alleles. When chimpanzee alignments are available, the non-chimp allele is coloured; otherwise the minor allele is coloured. B) Overview of introgression tracks surrounding the *SLC390A9* gene region as inferred by ARGweaver and available at the UCSC Genome Browser. In blue, Denisovan to Papuan introgression at p=0.5 cut-off (chr4: 41977301-42032370); in green, human to Vindija/Altaic Neanderthal introgression at p=0.5 cut-off (chr4: 41827151-41986310, chr4: 41801111-41824760 and chr4: 41978271-41983950); in dark grey, Super-Archaic hominin to Mandenka 2F introgression regions at p=0.5 cut-off (chr4:42050001-42071760); in light grey, Super-Archaic hominin to KhomaniSan 1F introgressed regions at p=0.5 cut-off (chr4:42050001-42071760).

**Figure S25.**
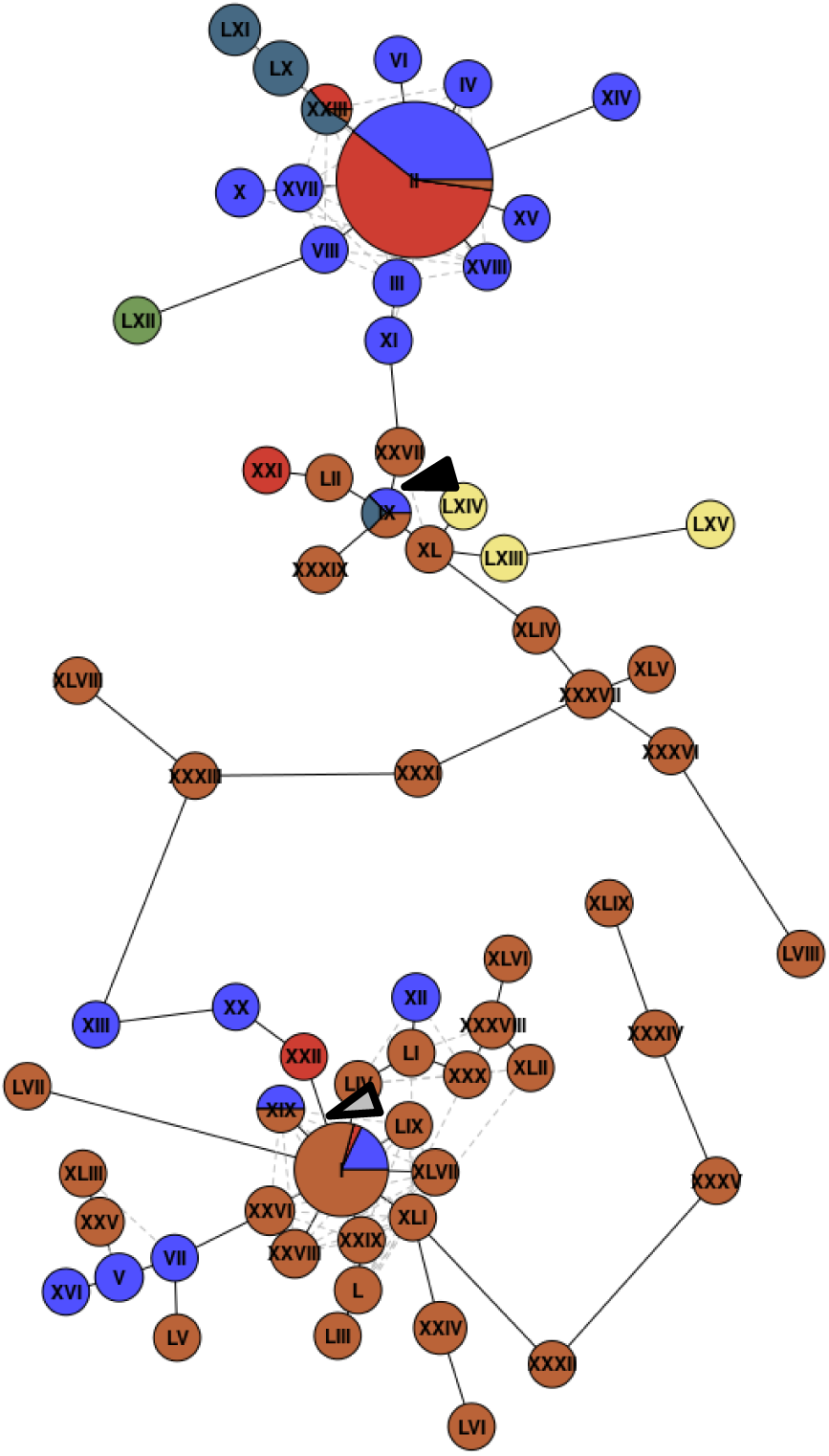
Network of modern and archaic human haplotypes. Haplotypes were defined by the complete set of SNPs available with a homozygous genotype in the Denisovan genome along the putatively introgressed region (chr4: 41,977,828-42,048,441; GRCh38). Pie charts show the frequency of each haplotype among YRI, CEU, CHB, and Oceanians (OCE). Denisovan and Neanderthal haplotypes are show in green and yellow, respectively. Arrow in black, position of the rs1047626 substitution; arrow in grey, position of the rs4861157 polymorphism. Haplotype frequencies, and polymorphic positions defining each haplotype are available in Table S5. For nucleotide pairwise distances between haplotypes, see Table S6.

**Figure S26.**
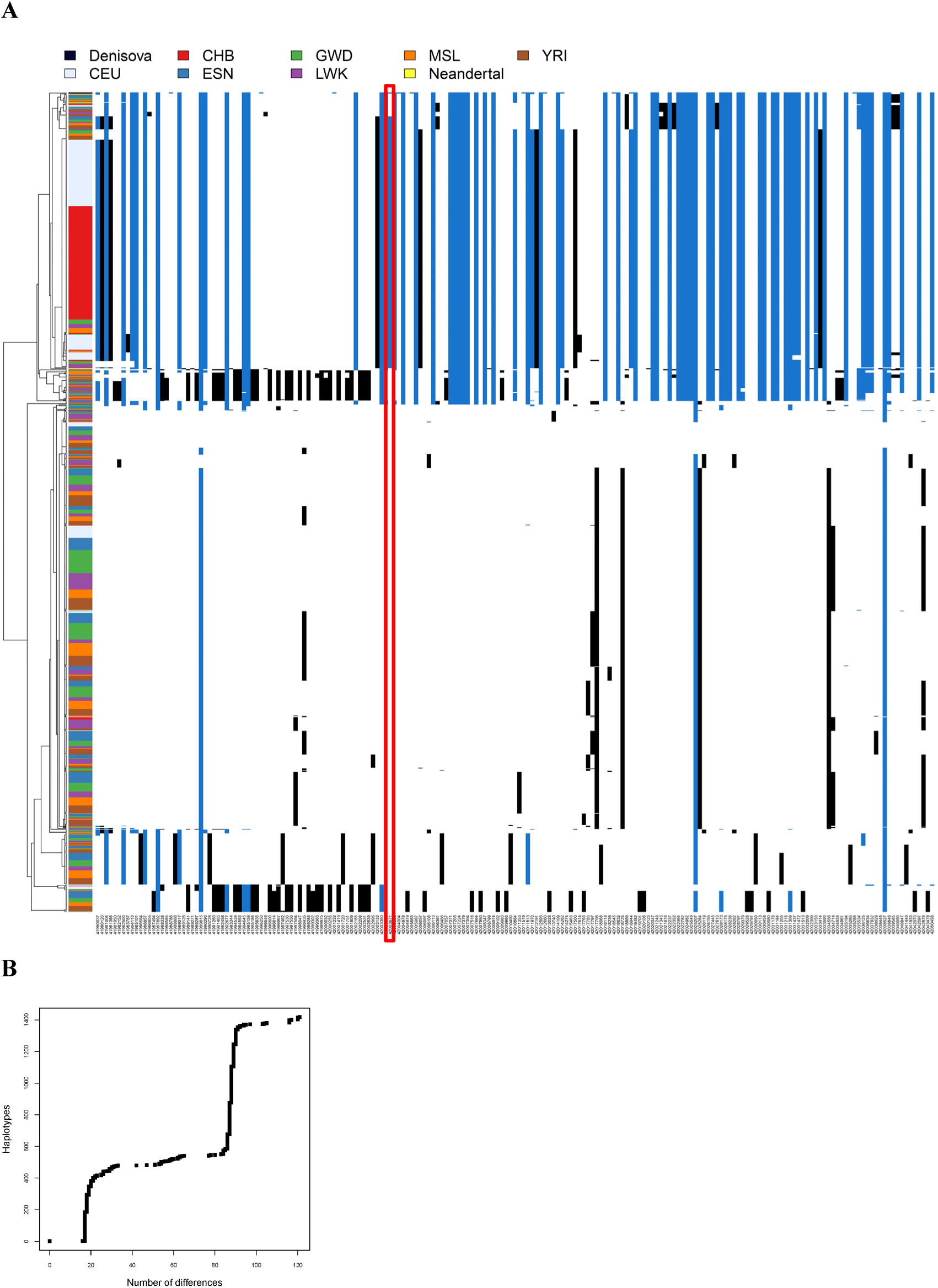
*SLC30A9* haplotype variation and structure when considering an extended panel of African populations. Note that Oceanians and Chagyrskaya were not included in the analysis to maximize the number of SNPs in the haplotypes. A total of 195 SNPs along the *SLC30A9* region (chr4: 41975811-42046424; GRCh37) defined 97 unique haplotypes when considering the three remaining high coverage archaic genomes (Altai, Vindija and Denisova), and the CEU and CHB populations together with an extended panel of African populations (ESN, Esan in Nigeria; YRI, Yoruba in Ibadan, Nigeria; MSL, Mende in Sierra Leone; GWD, Gambian Mandinka; LWK, Luhya in Webuye, Kenya). Note that all sites with a maximum within-population minor allele frequency below 0.05 were removed. A) Haplotypes depicted in decreasing order according to their similarity to the Denisovan genome, shown in the top. Each column corresponds to a SNP. The white colour in each cell indicates the ancestral allele; blue, indicates a derived allele shared with the Denisovan genome; whereas black indicates a derived allele that is not shared with the Denisova. The red box highlights the location of rs1047626. B) Number of differences observed to the Denisovan haplotype (see values and individual haplotype codes per population in Table S8).

**Figure S27.**
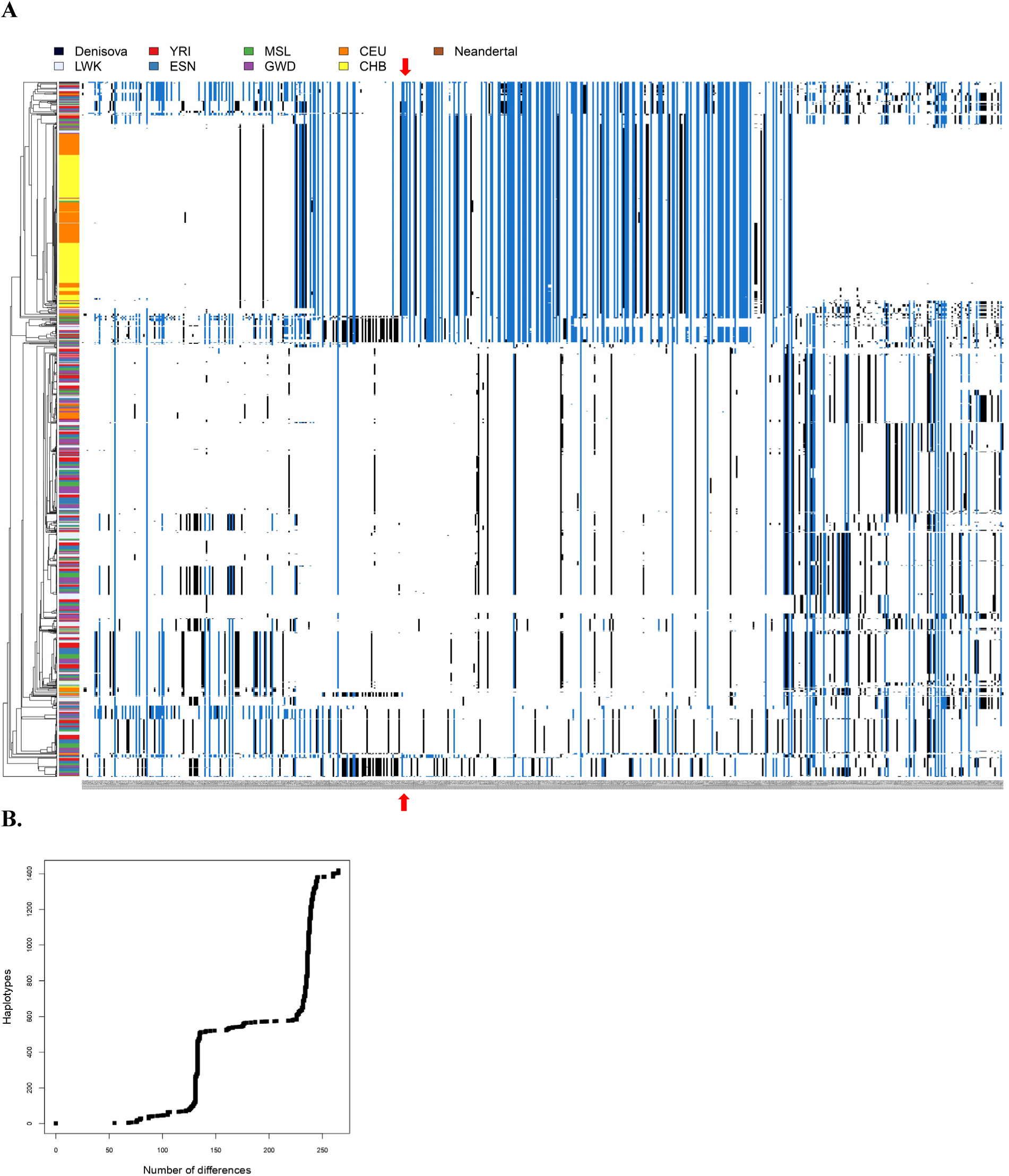
*SLC30A9* haplotype variation and structure in an extended genomic region. Note that Oceanians and Chagyrskaya were not included in the analysis to maximize the number of SNPs in the haplotypes. A total of 603 SNPs in an extended genomic region around the *SLC30A9* gene (chr4:41905811-42146424; GRCh37) defined 347 unique haplotypes when considering the three remaining high coverage archaic genomes (Altai, Vindija and Denisova), and the CEU and CHB populations together with an extended panel of African populations (ESN, Esan in Nigeria; YRI, Yoruba in Ibadan, Nigeria; MSL, Mende in Sierra Leone; GWD, Gambian Mandinka; LWK, Luhya in Webuye, Kenya). Note that all sites with a maximum within-population minor allele frequency below 0.05 were removed. A) Haplotypes depicted in decreasing order according to their similarity to the Denisovan genome, shown in the top. Each column corresponds to a SNP. The white colour in each cell indicates the ancestral allele; blue, indicates a derived allele shared with the Denisovan; whereas black indicates a derived allele that is not shared with the Denisova. The red arrow highlights the location of rs1047626.B) Number of differences observed to the Denisovan haplotype (see values and individual haplotype codes per population in Table S9).

**Figure S28.**
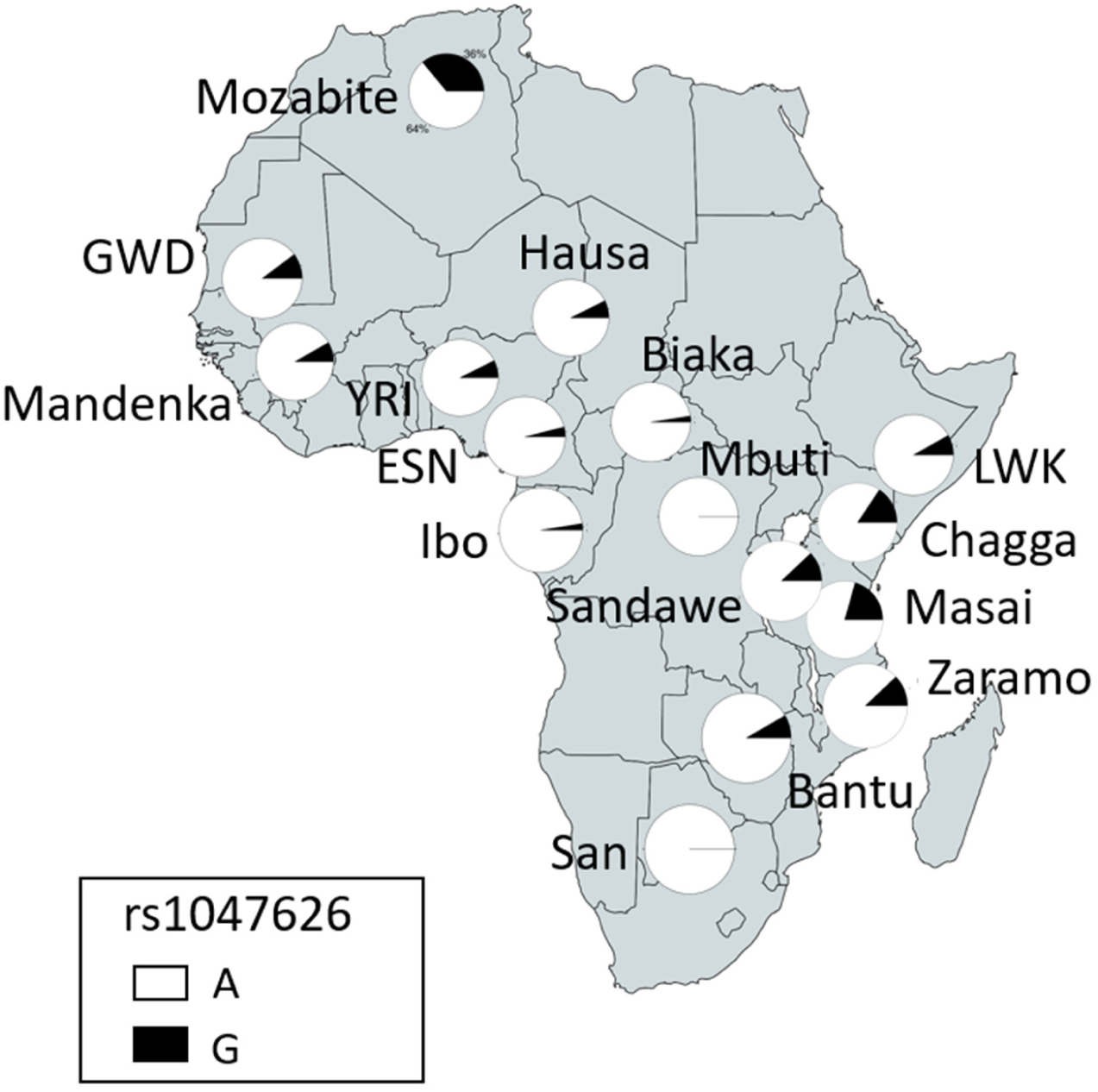
Allele frequencies for the rs1047626 polymorphism in Africa. Allele frequencies for the Mbuti, Mozabite, Hausa, San, Ibo, Chagga, Masai, Sandawe and Zaramo were compiled from the ALFRED database (https://alfred.med.yale.edu/alfred/index.asp)[5], for ESN, GWD, LWK, MSL and YRI from the 1000 Genomes Project [1] data available at Ensembl (https://www.ensembl.org/index.html), and for Biaka, Bantu, and Mandenka from the Human Genome Diversity Panel (HGDP-CEPH)[6].

**Figure S29.**
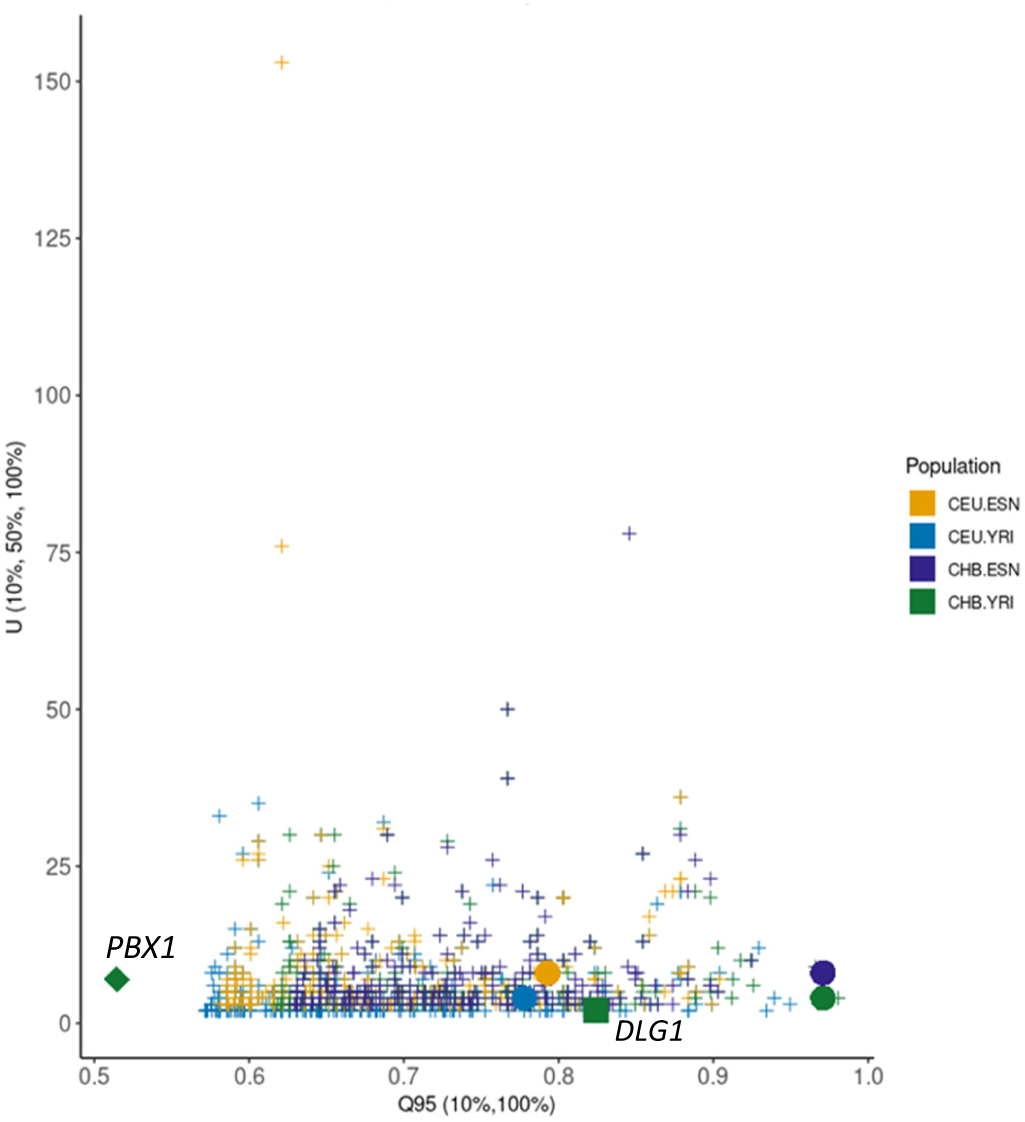
Joint distribution of the U (10%, 50%, 100%) and Q95 (10%, 100%) statistics in CEU and CHB when using either YRI or ESN as unadmixed group. Circles denote the U (10%, 50%, 100%) and Q95 (10%, 100%) values in the *SLC30A9* region, whereas the square and the rombe denote the statistics values in two candidate regions for Denisovan introgression in CHB [7].

**Figure S30.**
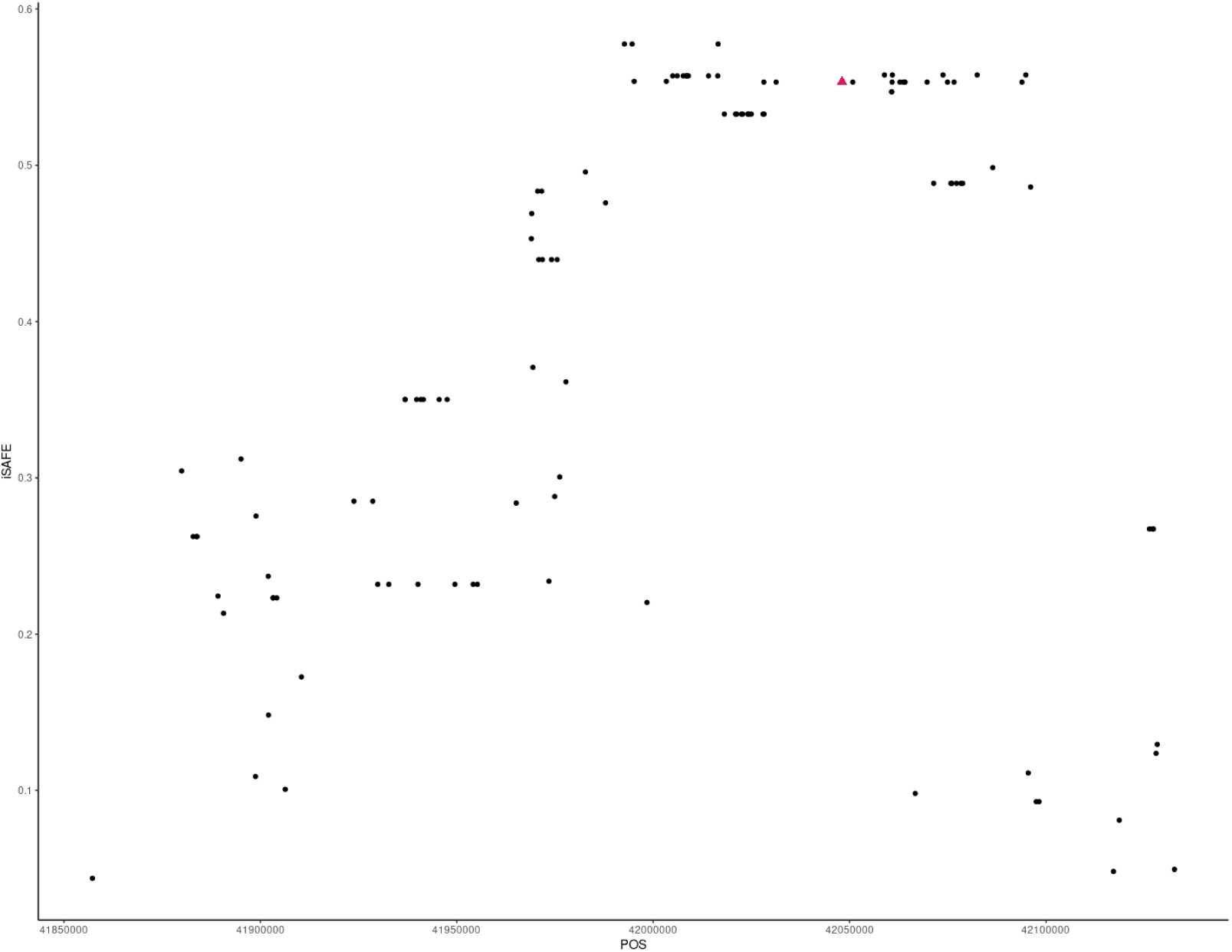
iSAFE values in YRI. iSAFE values computed along the *SLC30A9* candidate region for selection (chr4: 41853671 – 42153671, GRCh37/hg19) in the Yoruba population. In red, iSAFE value for rs4861157. Complete details on the SNP annotation and iSAFE values are available in Table S15.

**Figure S31.**
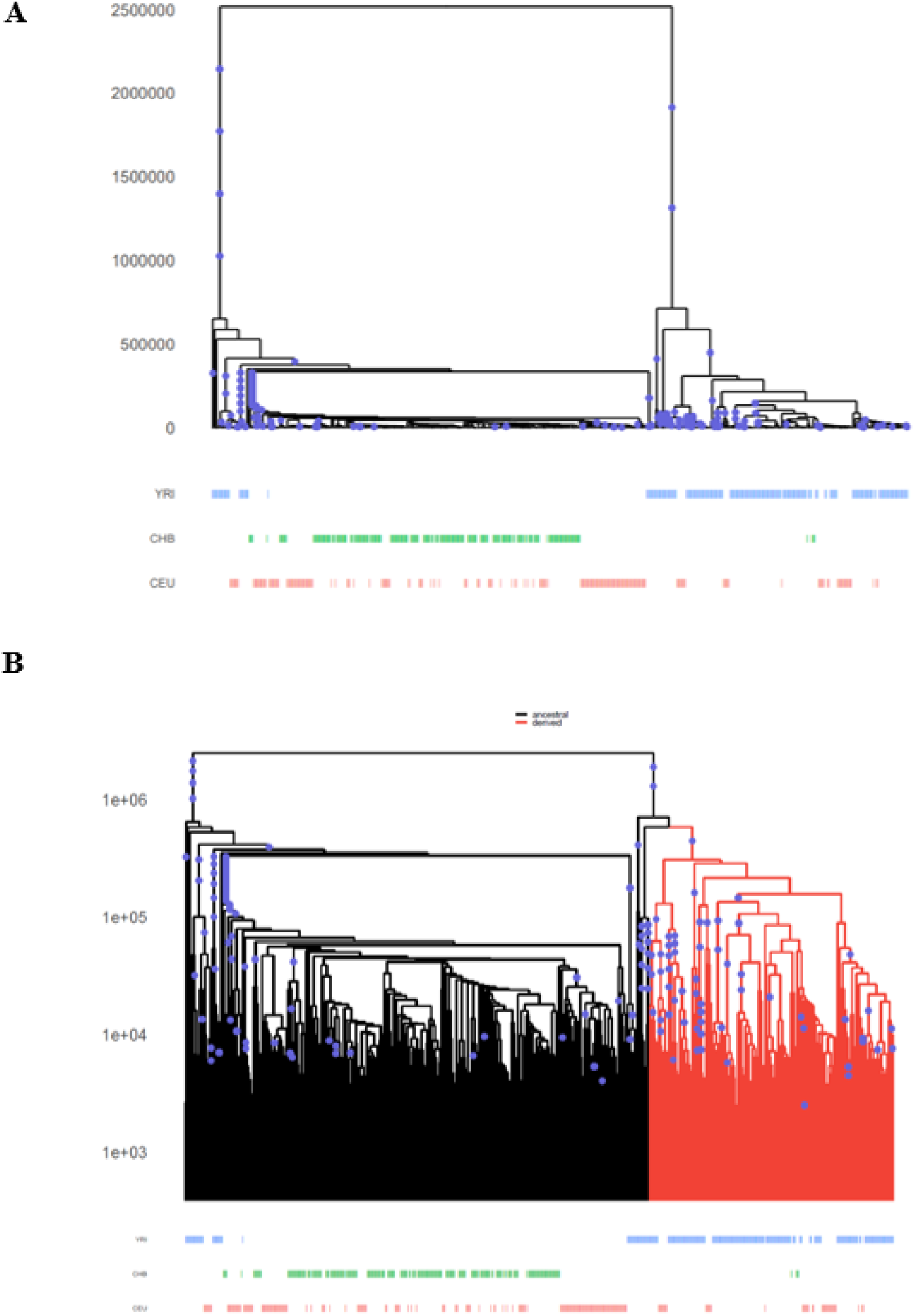
Positive selection acting on rs4861157. A) Marginal tree corresponding to the rs4861157 flanking region (chr4: 42022464-42025285; GRCh37/hg19) at *SLC30A9*. The derived allele at this SNP expanded rapidly in YRI, which is indicative of positive selection. B) Tree of interest highlighting lineages carrying the derived allele at rs4861157.

**Figure S32.**
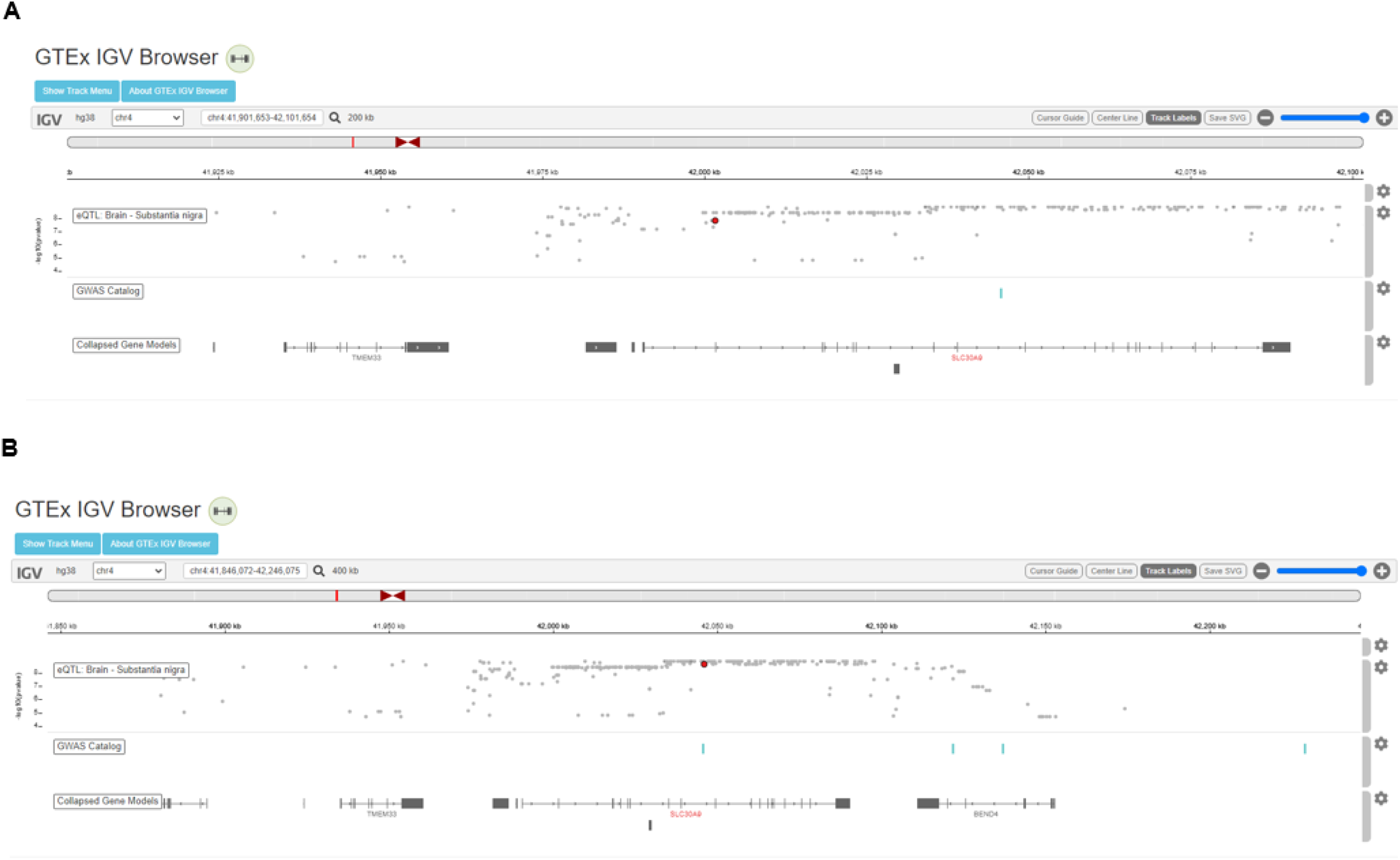
eQTLs affecting gene expression in the substantia nigra around the *SLC30A9* genomic region. A) GTEx IGV Browser image showing rs1047626 significance as an eQTL (red dot). B) GTEx IGV Browser image showing rs4861157 significance as an eQTL (red dot).

**Figure S33.**
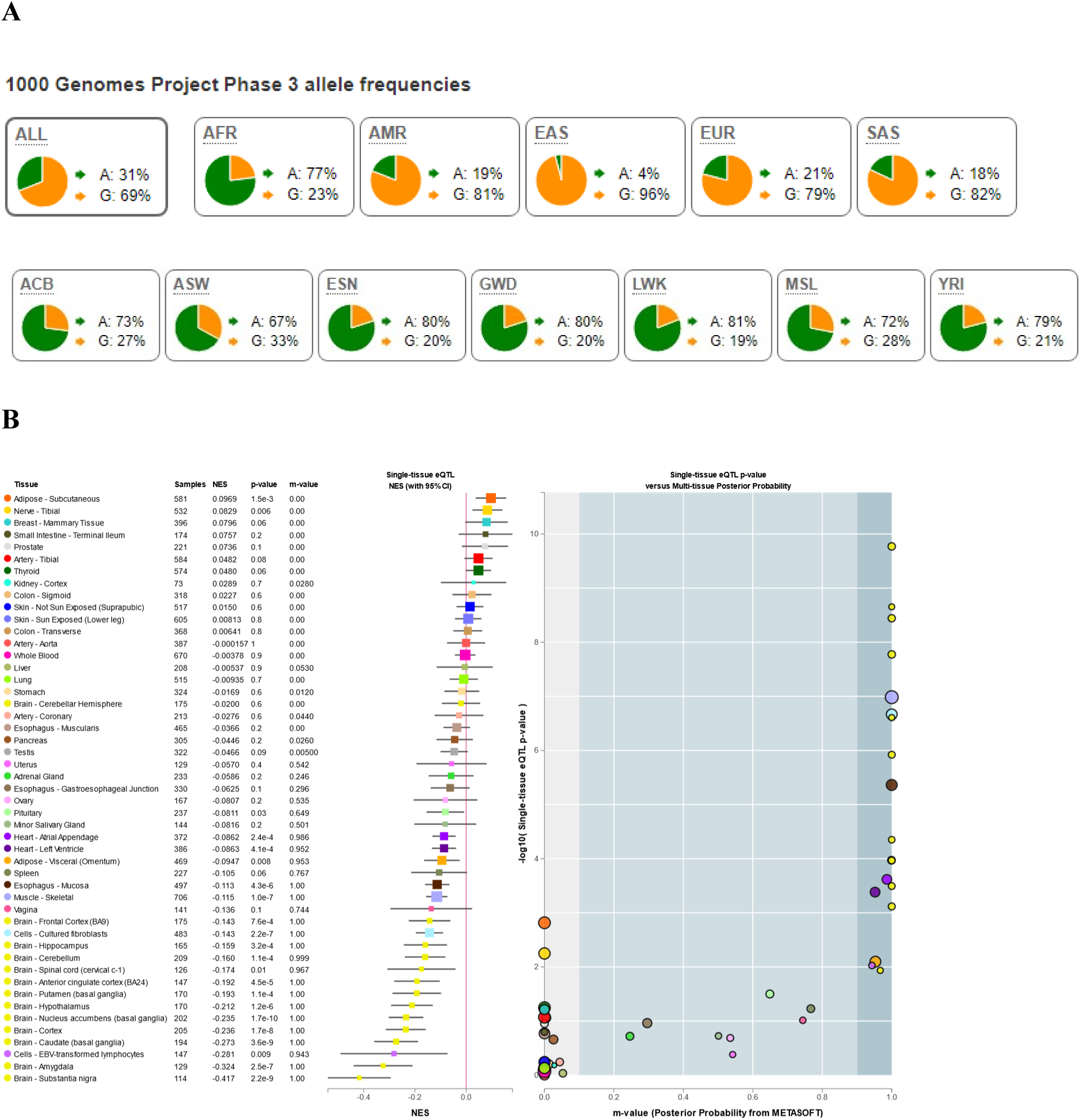
Continental allele frequencies and GTEX data for rs4861157. (A) Continental and African sub-populations 1000 Genomes Project Phase 3 allele frequencies as retrieved from Ensembl (https://www.ensembl.org/index.html). (B) Multi-tissue eQTL comparison for rs4861157.

**Figure S34.**
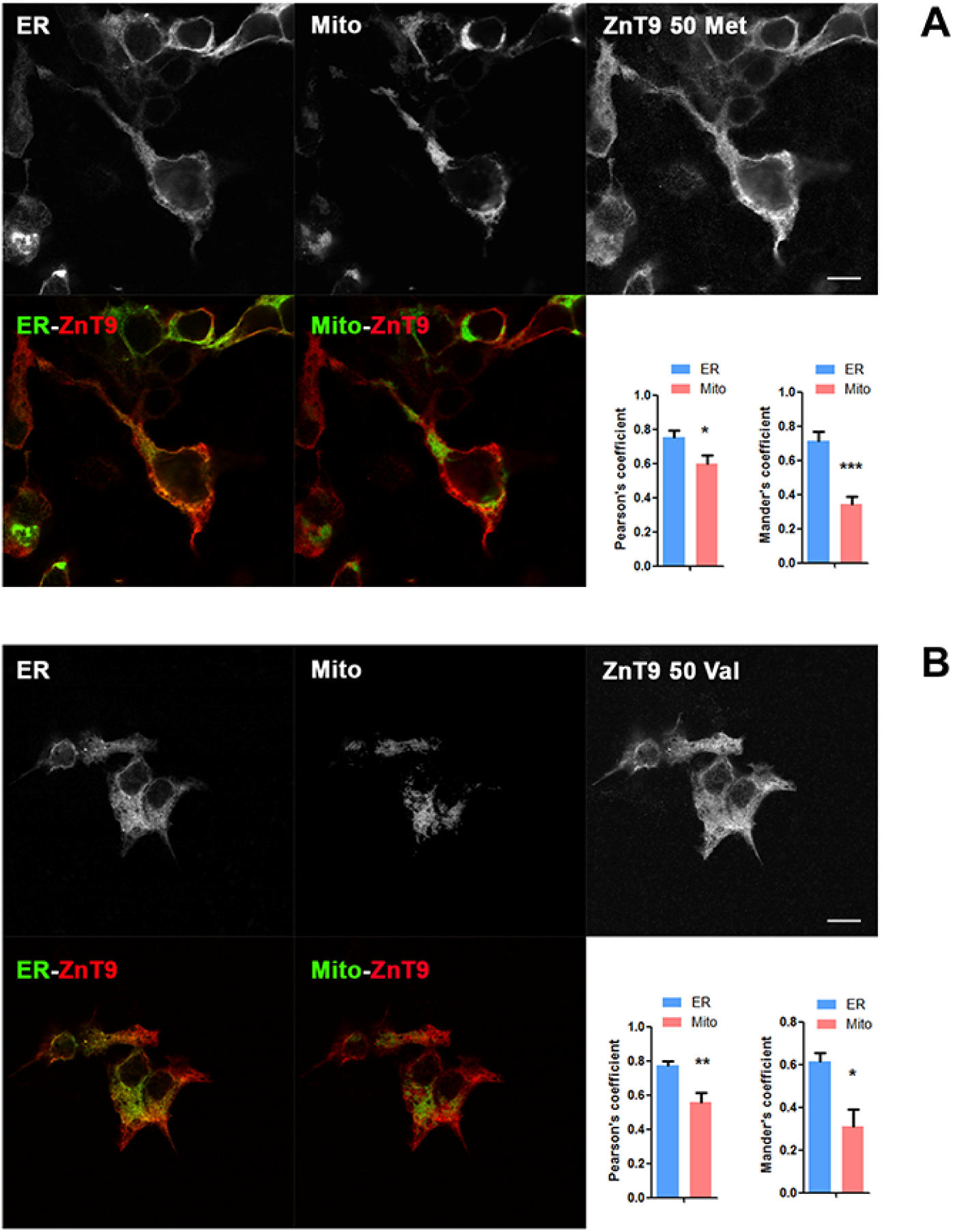
**Characterization of the localization of ZnT9 transporter in HEK293 cells**. Representative pictures of colocalization of ZnT9-50Met (A) and ZnT9-50Val (B) (both in red) with a double fluorescent reporter plasmid from the endoplasmic reticulum and the mitochondria (both in green in separate images) Scale bar = 10 µm. Representation of Pearson’s correlation coefficient and Mander’s overlap coefficient for quantifying the colocalization of ZnT9-50Met (n = 5) and ZnT9-50Val (n = 4) with endoplasmic and mitochondria. *** p < 0.001, ** p < 0.01, * p < 0.05. See further statistical details in File S1.

**Figure S35.**
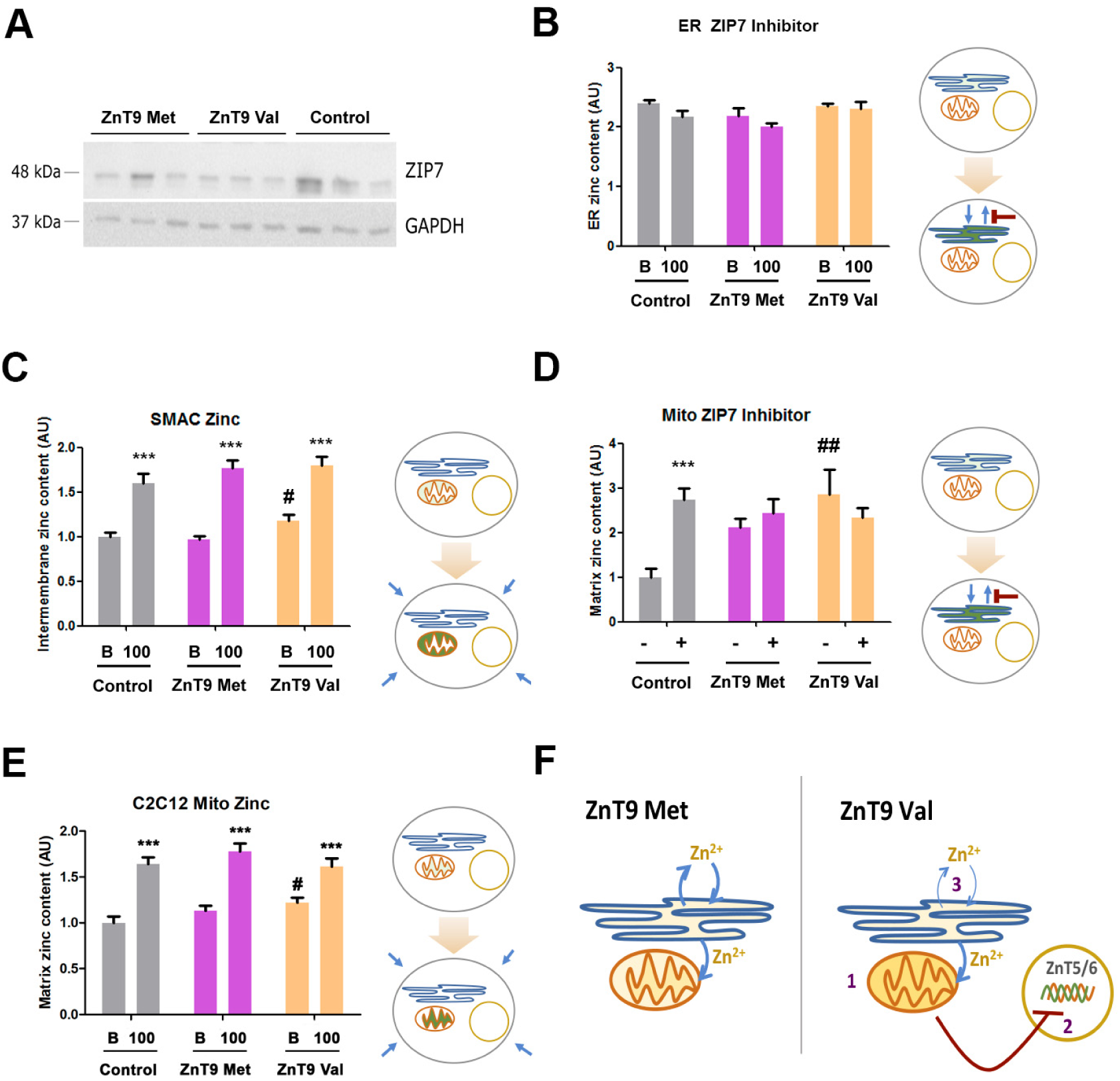
**Zip9 expression and zinc homeostasis**. (A) Representative Western blot against ZIP7 and GAPDH in HEK293 cells transfected with ZnT9-50Met, ZnT9-50Val, or an empty vector. (B) Evaluation of endoplasmic zinc content in HEK293 cells using an ER fluorescent zinc sensor (ER-ZapCY1) in basal and 100 µM ZnSO4 in the presence of DMSO and Zip7 blocker (n = 12-19). (C) Evaluation of zinc mitochondrial intermembrane space content in HEK293 cells using SMAC-Gn2Zn probe in conditions of 10 µM and 100 µM ZnSO4 (n = 8-9); *** p<0.001 using t-test between basal and 100 µM zinc conditions, # p<0.05 using Bonferroni-corrected ANOVA between transfection conditions. (D) Evaluation of zinc mitochondrial matrix content in HEK293 cells using Mito-cCherry-Gn2Zn incubating 40 min in the presence of DMSO or the Zip7 blocker (n = 6-8). *** p<0.001 using t-test between DMSO and blocker, ## p<0.05 using Bonferroni-corrected ANOVA between transfection conditions. (E) Evaluation of zinc mitochondrial matrix content in C2C12 cells using Mito-cCherry-Gn2Zn incubating 40min with basal and 100 µM ZnSO4 conditions (n = 21-28); *** p<0.01 using t-test between basal and 100 µM zinc conditions, # p<0.05 using Bonferroni-corrected ANOVA between transfection conditions. (F) Schematic model of the effect of the ZnT9 variants in zinc homeostasis. 1, ZnT9 Val expressing cells contain higher mitochondrial zinc content. Mitochondrial zinc is dependent of endoplasmic reticulum zinc content. 2, A higher mitochondrial zinc could be compensated in order to avoid overload with the transcriptional repression of the endoplasmic zinc importers ZnT5 and ZnT6. 3, ER zinc content is not altered because the zinc exporter Zip7 is the master regulator of ER zinc content. See further statistical details in File S1.

**Figure S36.**
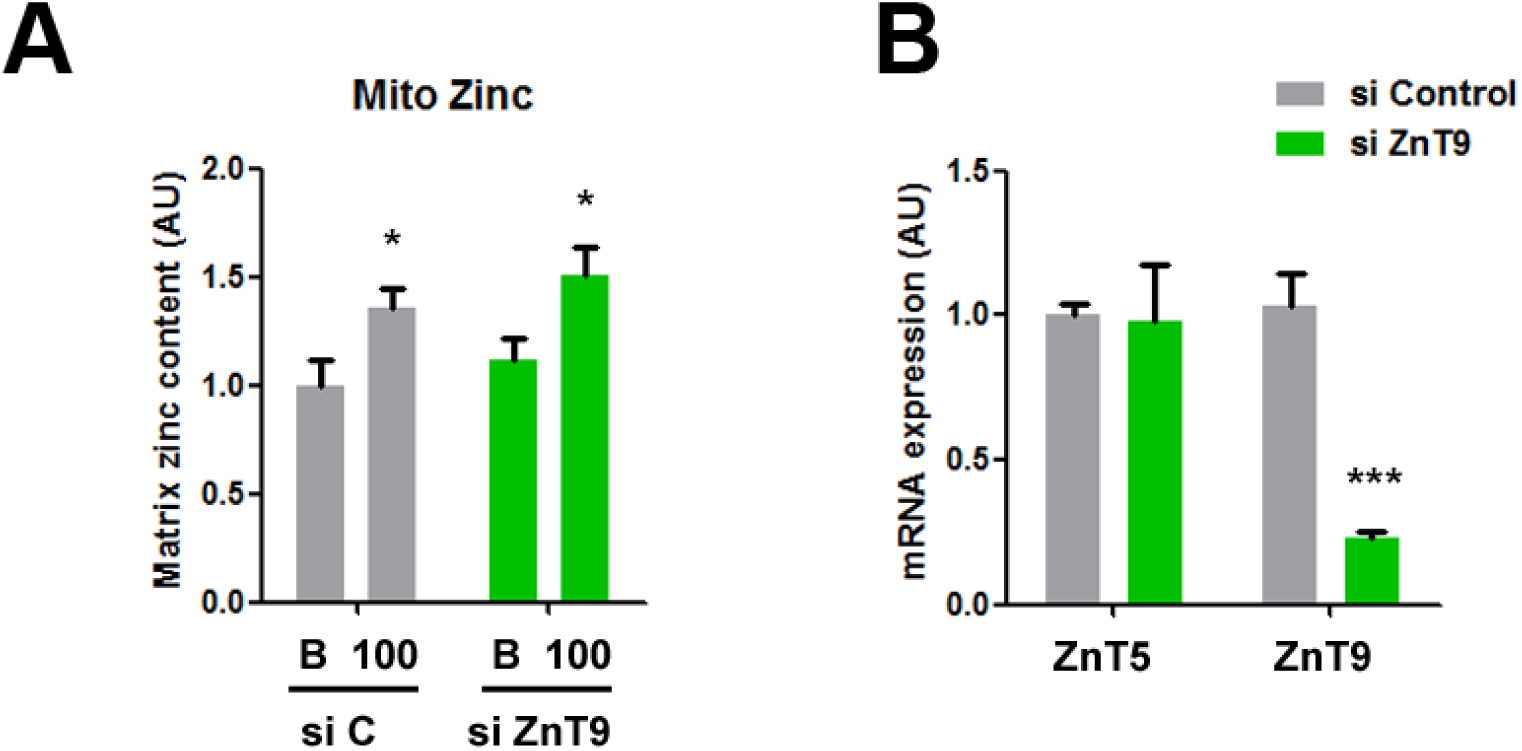
ZnT9 silencing experiments. (A) Evaluation of zinc mitochondrial matrix content in siRNAControl (siC) and siRNA ZnT9 (siZnT9) transfected cells using Mito-cCherry-Gn2Zn 40min with basal and 100 µM ZnSO4 conditions (n = 10-11); * p<0.05 using t-test. (B) RNA expression analysis of ZnT5 and ZnT9 cells transfected with siRNAControl (siC) and siRNA ZnT9 (siZnT9). 2-(DDCT) plotted using GAPDH as the housekeeping gene (n = 6); *** p<0.001 using t-test. See further statistical details in File S1.

**Figure S37.**
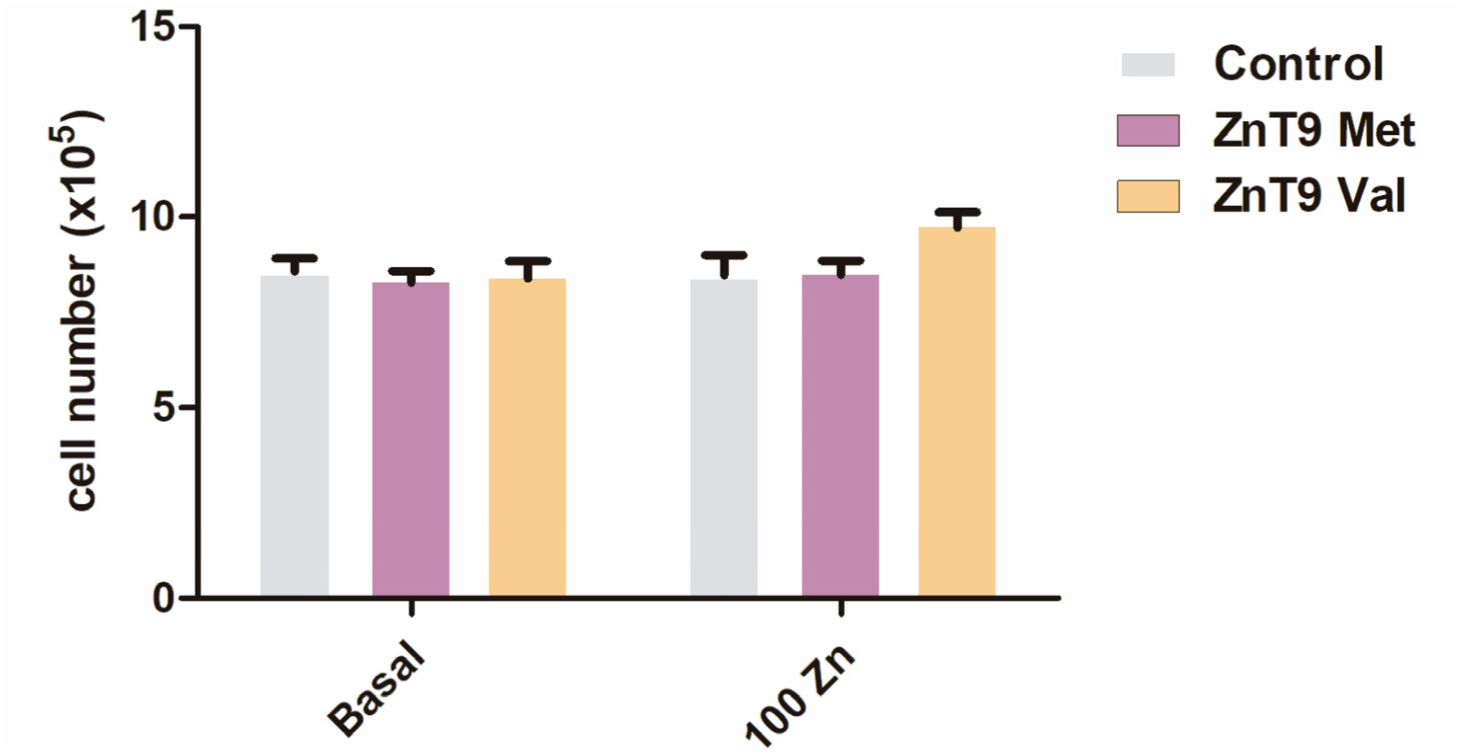
Toxicity evaluation in ZnT9 transfected cells. Cell counting assay in cells transfected with ZnT9-50Met, ZnT9-50Val, or an empty vector incubated for 24h at basal and 100 µM ZnSO4 conditions. Data are reported as mean ± SE (error bars) (n=8-12). No statistically significant differences were found. See further statistical details in File S1.

**Figure S38.**
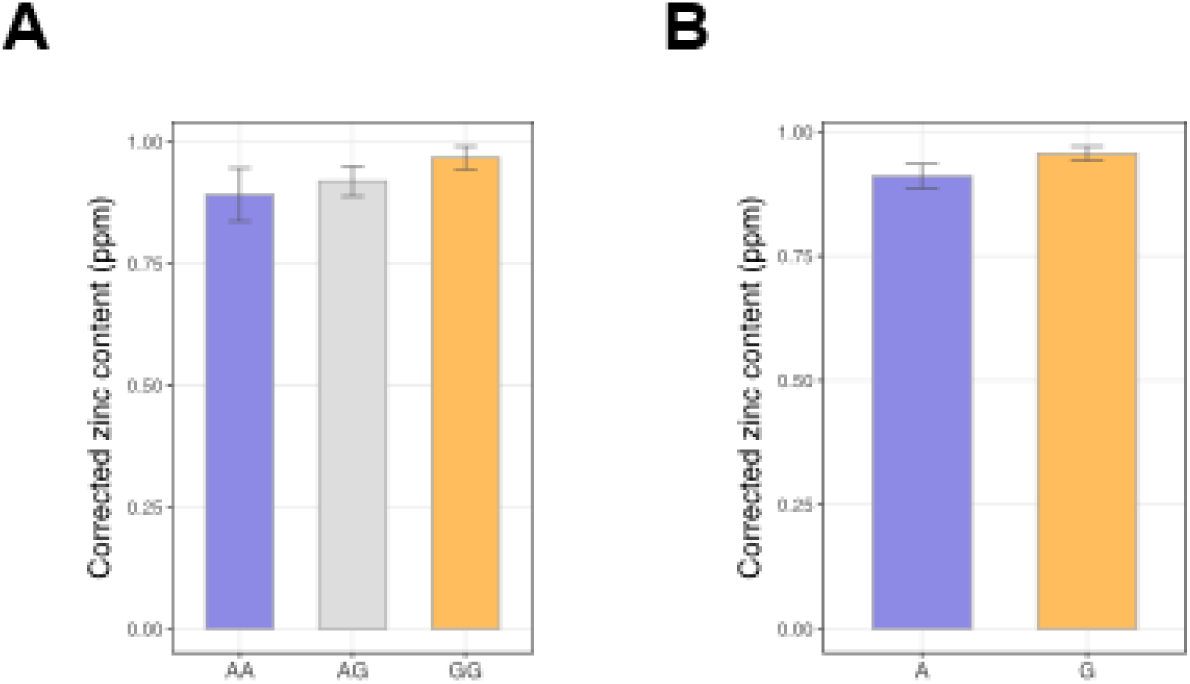
Zinc content in liver samples according to rs1047626 genotype. (A) Representation of liver zinc content comparing individuals with either the AA (n=11), AG (n=52) or GG (n=80) genotypes at rs1047626. (B) Representation of liver zinc content according to the A (n=74) or G (n=212) alleles at rs1047626. No statistically significant differences were found despite the observed tendency towards higher zinc concentrations in individuals carrying the derived G-allele. Data are reported as mean ± SE (error bars). Liver zinc content corresponds to the corrected zinc levels previously quantified [8] in the liver samples genotyped here for the rs1047626 polymorphism.

## Tables S1-S20 (separate excel file)

**Table S1.** List of functionally relevant candidate SNPs in the *SLC30A9* region.

**Table S2.** Top 40 candidate variants in CHB identified by iSAFE.

**Table S3.** Top 40 candidate variants in CHB identified by iSAFE.

**Table S4.** SNPs in high linkage disequilibrium (r^2^>0.8) with rs1047626 in CEU or CHB and their corresponding allelic states found in the Neanderthal and Denisovan genomes.

**Table S5.** Archaic and modern human haplotypes inferred along the putative *SLC30A9* introgressed region and their corresponding frequencies in YRI, CHB, CEU and Oceanians.

**Table S6**. Pairwise distances between haplotypes inferred along the putative *SLC30A9* introgressed region.

**Table S7.** Haplostrips distance output using four high-coverage archaic humans and the OCE, CHB, CEU and YRI populations (chr4: 41,977,828-42,048,441; GRCh38).

**Table S8**. Haplostrips distance output using three high-coverage archaic humans together with the CHB, CEU populations and an extended African dataset comprising ESN, YRI, MSL, GWD, and LWK (chr4: 41,975,811-42,046,424; GRCh37).

**Table S9.** Haplostrips distance output using three high-coverage archaic humans together with the CHB, CEU populations and an extended African dataset in a larger genomic region around *SLC30A9* (chr4: 41,905,811-42,146,424; GRCh37).

**Table S10**. Parameters for determining the probability of Incomplete Lineage Sorting (ILS).

**Table S11.** U and Q95 introgression statistics.

**Table S12**. eQTLS in high linkage disequilibrium (r^2^>0.8) with rs1047626.

**Table S13.** For the YRI population, Relate p-values in the *SLC30A9* region.

**Table S14.** Functional annotation of SNPs with Relate p-value < 0.05 at the *SLC30A9* region in the YRI population.

**Table S15.** Top 30 candidate variants in YRI identified by iSAFE.

**Table S16.** eQTLs with Relate p-value < 0.05 at the *SLC30A9* region in the YRI population.

**Table S17.** Linkage disequilibrium (r^2^) found in CEU and JPT (as proxy for Europeans and East Asians) between all putative candidate variants indicated in Table 1 and three nutriQTLS previously described at the 3’ region of *SLC30A9*.

**Table S18.** Zinc liver content and rs1047626 genotypes in individuals of European ancestry.

**Table S19.** PheWAS analysis for rs1047626 (Gene Atlas, last accessed 16/05/2022).

**Table S20**. List of primers used for RT-PCR analyses.

**Data File S1.** Raw data and statistics for the experimental data presented in Figs 4-5, and Figs S34-S37.

## References

1. Vitti JJ, Grossman SR, Sabeti PC. Detecting natural selection in genomic data. Annu Rev Genet. 2013; 47: 97–120. doi:10.1146/annurev-genet-111212-133526

2. Rees JS, Castellano S, Andrés AM. The Genomics of Human Local Adaptation. Trends Genet. 2020; 36: 415–428. doi:10.1016/j.tig.2020.03.006

3. Akey JM. Constructing genomic maps of positive selection in humans: Where do we go from here? Genome Res. 2009; 19: 711–722. doi:10.1101/gr.086652.108

4. Lachance J, Tishkoff SA. Population Genomics of Human Adaptation. Annu Rev Ecol Evol Syst. 2013; 44: 123–143. doi:10.1146/annurev-ecolsys-110512-135833

5. Fan S, Hansen MEB, Lo Y, Tishkoff SA. Going global by adapting local: A review of recent human adaptation. Science 2016; 354: 54–59. doi:10.1126/science.aaf5098

6. International Zinc Nutrition Consultative Group (IZiNCG). Assessment of the Risk of Zinc Deficiency in Populations and Options for Its Control. Hotz C, Brown KH, editors. Food and Nutrition Bulletin. 2004.

7. Kambe T, Tsuji T, Hashimoto A, Itsumura N. The Physiological, Biochemical, and Molecular Roles of Zinc Transporters in Zinc Homeostasis and Metabolism. Physiol Rev. 2015; 95: 749–784. doi:10.1152/physrev.00035.2014

8. Vallee BL, Falchuk KH. The Biochemical Basis of Zinc Physiology. Physiol Rev. 1993; 73: 79–118.

9. Hara T, Takeda T-A, Takagishi T, Fukue K, Kambe T, Fukada T. Physiological roles of zinc transporters: molecular and genetic importance in zinc homeostasis. J Physiol Sci. 2017; 67: 283–301. doi:10.1007/s12576-017-0521-4

10. Fukada T, Kambe T. Molecular and genetic features of zinc transporters in physiology and pathogenesis. Metallomics. 2011;3: 662–674. doi:10.1039/c1mt00011j

11. Huang L, Tepaamorndech S. The SLC30 family of zinc transporters-A review of current understanding of their biological and pathophysiological roles. Mol Aspects Med. 2013; 34: 548–560. doi:10.1016/j.mam.2012.05.008

12. Jeong J, Eide DJ. The SLC39 family of zinc transporters. Mol Aspects Med. 2013; 34: 612– 619. doi:10.1016/j.mam.2012.05.011

13. Zhang C, Li J, Tian L, Lu D, Yuan K, Yuan Y, et al. Differential natural selection of human zinc transporter genes between african and non-African populations. Sci Rep. 2015; 5: 9658. doi:10.1038/srep09658

14. Barreiro LB, Laval G, Ne Quach H, Patin E, Quintana-Murci L. Natural selection has driven population differentiation in modern humans. Nat Genet. 2008; 40: 340–345. doi:10.1038/ng.78

15. Engelken J, Carnero-Montoro E, Pybus M, Andrews GK, Lalueza-Fox C, Comas D, et al. Extreme population differences in the human zinc transporter ZIP4 (SLC39A4) are explained by positive selection in Sub-Saharan Africa. PLoS Genet. 2014; 10: e1004128. doi:10.1371/journal.pgen.1004128

16. Carlson CS, Thomas DJ, Eberle MA, Swanson JE, Livingston RJ, Rieder MJ, et al. Genomic regions exhibiting positive selection identified from dense genotype data. Genome Res. 2005; 15: 1553–1565. doi:10.1101/GR.4326505

17. Grossman SR, Andersen KG, Shlyakhter I, Tabrizi S, Winnicki S, Yen A, et al. Identifying Recent Adaptations in Large-Scale Genomic Data. Cell. 2013; 152: 703–713. doi:10.1016/J.CELL.2013.01.035

18. Pickrell JK, Coop G, Novembre J, Kudaravalli S, Li JZ, Absher D, et al. Signals of recent positive selection in a worldwide sample of human populations. Genome Res. 2009; 826– 837. doi:10.1101/gr.087577.108

19. Sabeti PC, Varilly P, Fry B, Lohmueller J, Hostetter E, Cotsapas C, et al. Genome-wide detection and characterization of positive selection in human populations. Nature. 2007; 449: 913–8. doi:10.1038/nature06250

20. Williamson SH, Hubisz MJ, Clark AG, Payseur BA, Bustamante CD, Nielsen R. Localizing Recent Adaptive Evolution in the Human Genome. PLoS Genet. 2007; 3: e90. doi:10.1371/journal.pgen.0030090

21. Deng H, Qiao X, Xie T, Fu W, Li H, Zhao Y, et al. SLC-30A9 is required for Zn2+ homeostasis, Zn2+ mobilization, and mitochondrial health. Proc Natl Acad Sci U S A. 2021; 118. doi:10.1073/pnas.2023909118

22. Ma T, Zhao L, Zhang J, Tang R, Wang X, Liu N, et al. A pair of transporters controls mitochondrial Zn 2+ levels to maintain mitochondrial homeostasis. Protein Cell. 2021; 13(3):180–202. doi:10.1007/S13238-021-00881-4

23. Engelken J, Espadas G, Mancuso FM, Bonet N, Scherr AL, Jimenez-Alvarez V, et al. Signatures of evolutionary adaptation in quantitative trait loci influencing trace element homeostasis in liver. Mol Biol Evol. 2016; 33: 738–754. doi:10.1093/molbev/msv267

24. Roca-Umbert A, Caro-Consuegra R, Londono-Correa D, Rodriguez-Lozano GF, Vicente R, Bosch E. Understanding signatures of positive natural selection in human zinc transporter genes. Sci Rep. 2022; 12: 4320. doi:10.1038/s41598-022-08439-y

25. Sabeti PC, Schaffner SF, Lohmueller J, Varilly P, Shamovsky O, Palma A, et al. Positive Natural Selection in the Human Lineage. Science 2006; 312: 1614–1620. doi:10.1126/science.1124309

26. Casillas S, Mulet R, Villegas-Mirón P, Hervas S, Sanz E, Velasco D, et al. PopHuman: the human population genomics browser. Nucleic Acids Res. 2018; 46: D1003–D1010. doi:10.1093/NAR/GKX943

27. Kircher M, Witten DM, Jain P, O ’roak BJ, Cooper GM, Shendure J, et al. A general framework for estimating the relative pathogenicity of human genetic variants. Nat Genet. 2014; 46: 310–315. doi:10.1038/ng.2892

28. Akbari A, Vitti JJ, Iranmehr A, Bakhtiari M, Sabeti PC, Mirarab S, et al. Identifying the favored mutation in a positive selective sweep. Nat Methods. 2018; 15: 279–282. doi:10.1038/nmeth.4606

29. Speidel L, Forest M, Shi S, Myers SR. A method for genome-wide genealogy estimation for thousands of samples. Nat Genet. 2019; 51: 1321–1329. doi:10.1038/s41588-019-0484-x

30. Curtis D, Amos W. The human genome harbours widespread exclusive yin yang haplotypes. Eur J Hum Genet. 2023; doi:10.1038/s41431-023-01399-5

31. Browning SR, Browning BL, Zhou Y, Tucci S, Akey JM. Analysis of Human Sequence Data Reveals Two Pulses of Archaic Denisovan Admixture. Cell. 2018; 173: 53–61.e9. doi:10.1016/j.cell.2018.02.031

32. Sankararaman S, Mallick S, Patterson N, Reich D. The Combined Landscape of Denisovan and Neanderthal Ancestry in Present-Day Humans. Curr Biol. 2016; 26: 1241–7. doi:10.1016/j.cub.2016.03.037

33. Skov L, Coll Macià M, Sveinbjörnsson G, Mafessoni F, Lucotte EA, Einarsdóttir MS, et al. The nature of Neanderthal introgression revealed by 27,566 Icelandic genomes. Nature. 2020; 582: 78–83. doi:10.1038/s41586-020-2225-9

34. Gower G, Picazo PI, Fumagalli M, Racimo F. Detecting adaptive introgression in human evolution using convolutional neural networks. Elife. 2021; 10. doi:10.7554/eLife.64669

35. Hubisz MJ, Williams AL, Siepel A. Mapping gene flow between ancient hominins through demography-aware inference of the ancestral recombination graph. Schierup MH, editor. PLoS Genet. 2020; 16: e1008895. doi:10.1371/journal.pgen.1008895

36. The 1000 Genomes Project Consortium. A global reference for human genetic variation. Nature. 2015; 526: 68–74. doi:10.1038/nature15393

37. Bergström A, McCarthy SA, Hui R, Almarri MA, Ayub Q, Danecek P, et al. Insights into human genetic variation and population history from 929 diverse genomes. Science. 2020; 367. doi:10.1126/science.aay5012

38. Marnetto D, Huerta-Sánchez E. *Haplostrips* : revealing population structure through haplotype visualization. Methods Ecol Evol. 2017; 8: 1389–1392. doi:10.1111/2041-210X.12747

39. Hodgson JA, Mulligan CJ, Al-Meeri A, Raaum RL. Early Back-to-Africa Migration into the Horn of Africa. PLoS Genet. 2014; 10: e1004393. doi:10.1371/journal.pgen.1004393

40. Vicente M, Schlebusch CM. African population history: an ancient DNA perspective. Curr Opin Genet Dev. 2020; 62: 8–15. doi:10.1016/j.gde.2020.05.008

41. Browning SR, Browning BL, Zhou Y, Tucci S, Akey JM. Analysis of Human Sequence Data Reveals Two Pulses of Archaic Denisovan Admixture. Cell. 2018; 173: 53–61.e9. doi:10.1016/j.cell.2018.02.031

42. Bergström A, McCarthy SA, Hui R, Almarri MA, Ayub Q, Danecek P, et al. Insights into human genetic variation and population history from 929 diverse genomes. Science 2020; 367(6484):eaay5012. doi:10.1126/science.aay5012

43. Racimo F, Marnetto D, Huerta-Sánchez E. Signatures of Archaic Adaptive Introgression in Present-Day Human Populations. Mol Biol Evol. 2017; 34(2):296–317. doi: 10.1093/molbev/msw216.

44. Eide DJ. Zinc transporters and the cellular trafficking of zinc. Biochim Biophys Acta. 2006; 1763: 711–722. doi:10.1016/J.BBAMCR.2006.03.005

45. Xue J, Xie T, Zeng W, Jiang Y, Bai XC. Cryo-EM structures of human ZnT8 in both outward-and inward-facing conformations. Elife. 2020; 9: 1–32. doi:10.7554/ELIFE.58823

46. Chen L, Wolf AB, Fu W, Li L, Akey JM. Identifying and Interpreting Apparent Neanderthal Ancestry in African Individuals. Cell. 2020; 180: 677–687.e16. doi:10.1016/j.cell.2020.01.012

47. Peyrégne S, Kelso J, Peter BM, Pääbo S. The evolutionary history of human spindle genes includes back-and-forth gene flow with Neandertals. Elife. 2022; 11. doi:10.7554/eLife.75464

48. Chen L, Wolf AB, Fu W, Li L, Akey JM. Identifying and Interpreting Apparent Neanderthal Ancestry in African Individuals. Cell. 2020; 180: 677–687.e16. doi:10.1016/j.cell.2020.01.012

49. Jumper J, Evans R, Pritzel A, Green T, Figurnov M, Ronneberger O, et al. Highly accurate protein structure prediction with AlphaFold. Nature. 2021; 596: 583–589. doi:10.1038/s41586-021-03819-2

50. Kowalczyk A, Gbadamosi O, Kolor K, Sosa J, Andrzejczuk L, Gibson G, et al. Evolutionary rate covariation identifies SLC30A9 (ZnT9) as a mitochondrial zinc transporter. Biochem J. 2021; 478: 3205–3220. doi:10.1042/BCJ20210342

51. Ishihara K, Yamazaki T, Ishida Y, Suzuki T, Oda K, Nagao M, et al. Zinc Transport Complexes Contribute to the Homeostatic Maintenance of Secretory Pathway Function in Vertebrate Cells. J Biol Chem. 2006; 281: 17743–17750. doi:10.1074/JBC.M602470200

52. Suzuki T, Ishihara K, Migaki H, Ishihara K, Nagao M, Yamaguchi-Iwai Y, et al. Two Different Zinc Transport Complexes of Cation Diffusion Facilitator Proteins Localized in the Secretory Pathway Operate to Activate Alkaline Phosphatases in Vertebrate Cells. J Biol Chem. 2005; 280: 30956–30962. doi:10.1074/JBC.M506902200

53. Taylor KM, Hiscox S, Nicholson RI, Hogstrand C, Kille P. Protein kinase CK2 triggers cytosolic zinc signaling pathways by phosphorylation of zinc channel ZIP7. Sci Signal. 2012; 5. doi:10.1126/SCISIGNAL.2002585

54. Fudge DH, Black R, Son L, Lejeune K, Qin Y. Optical Recording of Zn 2+ Dynamics in the Mitochondrial Matrix and Intermembrane Space with the GZnP2 Sensor. ACS Chem Biol. 2018; 13: 1897–1905. doi:10.1021/ACSCHEMBIO.8B00319

55. Rensvold JW, Shishkova E, Sverchkov Y, Miller IJ, Cetinkaya A, Pyle A, et al. Defining mitochondrial protein functions through deep multiomic profiling. Nature. 2022; 606: 382–388. doi:10.1038/s41586-022-04765-3

56. Prinz WA, Toulmay A, Balla T. The functional universe of membrane contact sites. Nat Rev Mol Cell Biol. 2020; 21: 7–24. doi:10.1038/s41580-019-0180-9

57. Perez Y, Shorer Z, Liani-Leibson K, Chabosseau P, Kadir R, Volodarsky M, et al. SLC30A9 mutation affecting intracellular zinc homeostasis causes a novel cerebro-renal syndrome. Brain. 2017; 140: 928–939. doi:10.1093/brain/awx013

58. Coleman JRI, Gaspar HA, Bryois J, Byrne EM, Forstner AJ, Holmans PA, et al. The Genetics of the Mood Disorder Spectrum: Genome-wide Association Analyses of More Than 185,000 Cases and 439,000 Controls. Biol Psychiatry. 2020; 88: 169–184. doi:10.1016/J.BIOPSYCH.2019.10.015

59. Wray NR, Ripke S, Mattheisen M, Trzaskowski M, Byrne EM, Abdellaoui A, et al. Genome-wide association analyses identify 44 risk variants and refine the genetic architecture of major depression. Nat Genet. 2018; 50: 668–681. doi:10.1038/s41588-018-0090-3

60. Primes G, Fieder M. Real-life helping behaviours in North America: A genome-wide association approach. Novelli G, editor. PLoS One. 2018; 13: e0190950. doi:10.1371/journal.pone.0190950

61. Hahne F, Ivanek R. Visualizing Genomic Data Using Gviz and Bioconductor. Methods Mol Biol. 2016; 1418: 335–351. doi:10.1007/978-1-4939-3578-9_16

62. Wang K, Li M, Hakonarson H. ANNOVAR: functional annotation of genetic variants from high-throughput sequencing data. Nucleic Acids Res. 2010; 38. doi:10.1093/NAR/GKQ603

63. Ionita-Laza I, McCallum K, Xu B, Buxbaum JD. A spectral approach integrating functional genomic annotations for coding and noncoding variants. Nat Genet. 2016; 48(2):214–20. doi:10.1038/ng.3477

64. Gulko B, Hubisz MJ, Gronau I, Siepel A. A method for calculating probabilities of fitness consequences for point mutations across the human genome. Nat Genet. 2015; 47: 276–283. doi:10.1038/ng.3196

65. Stern AJ, Wilton PR, Nielsen R. An approximate full-likelihood method for inferring selection and allele frequency trajectories from DNA sequence data. Hernandez RD, editor. PLoS Genet. 2019; 15: e1008384. doi:10.1371/journal.pgen.1008384

66. Auton A, Abecasis GR, Altshuler DM, Durbin RM, Abecasis GR, Bentley DR, et al. A global reference for human genetic variation. Nature. 2015; 526: 68–74. doi:10.1038/nature15393

67. Delaneau O, Zagury J-F, Robinson MR, Marchini JL, Dermitzakis ET. Accurate, scalable and integrative haplotype estimation. Nat Commun. 2019; 10: 5436. doi:10.1038/s41467-019-13225-y

68. Paradis E. pegas: an R package for population genetics with an integrated-modular approach. Bioinformatics. 2010; 26: 419–420. doi:10.1093/bioinformatics/btp696

69. Huerta-Sánchez E, Jin X, Asan, Bianba Z, Peter BM, Vinckenbosch N, et al. Altitude adaptation in Tibetans caused by introgression of Denisovan-like DNA. Nature. 2014; 512: 194–197. doi:10.1038/nature13408

70. Dannemann M, Andrés AM, Kelso J. Introgression of Neandertal- and Denisovan-like Haplotypes Contributes to Adaptive Variation in Human Toll-like Receptors. Am J Hum Genet. 2016; 98: 22–33. doi:10.1016/j.ajhg.2015.11.015

71. Prüfer K, Racimo F, Patterson N, Jay F, Sankararaman S, Sawyer S, et al. The complete genome sequence of a Neanderthal from the Altai Mountains. Nature. 2014; 505: 43–49. doi:10.1038/nature12886

72. Buniello A, MacArthur JAL, Cerezo M, Harris LW, Hayhurst J, Malangone C, et al. The NHGRI-EBI GWAS Catalog of published genome-wide association studies, targeted arrays and summary statistics 2019. Nucleic Acids Res. 2019; 47: D1005–D1012. doi:10.1093/nar/gky1120

73. Watanabe K, Stringer S, Frei O, Umićević Mirkov M, de Leeuw C, Polderman TJC, et al. A global overview of pleiotropy and genetic architecture in complex traits. Nat Genet. 2019; 51: 1339–1348. doi:10.1038/s41588-019-0481-0

74. Naon D, Zaninello M, Giacomello M, Varanita T, Grespi F, Lakshminaranayan S, et al. Critical reappraisal confirms that Mitofusin 2 is an endoplasmic reticulum–mitochondria tether. Proc Natl Acad Sci U S A. 2016; 113: 11249–11254. doi:10.1073/PNAS.1606786113

75. Qin Y, Dittmer PJ, Park JG, Jansen KB, Palmer AE, Valentine JS. Measuring steady-state and dynamic endoplasmic reticulum and Golgi Zn 2þ with genetically encoded sensors. Proc Natl Acad Sci U S A. 2011; 108: 7351–7356. doi:10.1073/pnas.1015686108/-/DCSupplemental

## References

1. Auton A, Abecasis GR, Altshuler DM, Durbin RM, Abecasis GR, Bentley DR, et al. A global reference for human genetic variation. Nature. 2015;526: 68–74. doi:10.1038/nature15393

2. Marcus JH, Novembre J. Visualizing the geography of genetic variants. Bioinformatics. 2017;33: 594–595. doi:10.1093/BIOINFORMATICS/BTW643

3. Gower G, Picazo PI, Fumagalli M, Racimo F. Detecting adaptive introgression in human evolution using convolutional neural networks. Elife. 2021;10. doi:10.7554/eLife.64669

4. Hubisz MJ, Williams AL, Siepel A. Mapping gene flow between ancient hominins through demography-aware inference of the ancestral recombination graph. Schierup MH, editor. PLoS Genet. 2020;16: e1008895. doi:10.1371/journal.pgen.1008895

5. Rajeevan H. ALFRED: the ALelle FREquency Database. Update. Nucleic Acids Res. 2003;31: 270–271. doi:10.1093/nar/gkg043

6. Bergström A, McCarthy SA, Hui R, Almarri MA, Ayub Q, Danecek P, et al. Insights into human genetic variation and population history from 929 diverse genomes. Science (1979). 2020;367. doi:10.1126/science.aay5012

7. Racimo F, Marnetto D, Huerta-Sánchez E. Signatures of Archaic Adaptive Introgression in Present-Day Human Populations. Mol Biol Evol. 2016; msw216. doi:10.1093/molbev/msw216

8. Engelken J, Espadas G, Mancuso FM, Bonet N, Scherr AL, Jimenez-Alvarez V, et al. Signatures of evolutionary adaptation in quantitative trait loci influencing trace element homeostasis in liver. Mol Biol Evol. 2016;33: 738–754. doi:10.1093/molbev/msv267

